# Multivariate Genome-wide Association Analyses Reveal the Genetic Basis of Seed Fatty Acid Composition in Oat (*Avena sativa* L.)

**DOI:** 10.1101/589952

**Authors:** Maryn O. Carlson, Gracia Montilla-Bascon, Owen A. Hoekenga, Nicholas A. Tinker, Jesse Poland, Matheus Baseggio, Mark E. Sorrells, Jean-Luc Jannink, Michael A. Gore, Trevor H. Yeats

## Abstract

Oat (*Avena sativa* L.) has a high concentration of oils, comprised primarily of healthful unsaturated oleic and linoleic fatty acids. To accelerate oat plant breeding efforts, we sought to identify loci associated with variation in fatty acid composition, defined as the types and quantities of fatty acids. We genotyped a panel of 500 oat cultivars with genotyping-by-sequencing and measured the concentrations of ten fatty acids in these oat cultivars grown in two environments. Measurements of individual fatty acids were highly correlated across samples, consistent with fatty acids participating in shared biosynthetic pathways. We leveraged these phenotypic correlations in two multivariate genome-wide association study (GWAS) approaches. In the first analysis, we fitted a multivariate linear mixed model for all ten fatty acids simultaneously while accounting for population structure and relatedness among cultivars. In the second, we performed a univariate association test for each principal component (PC) derived from a singular value decomposition of the phenotypic data matrix. To aid interpretation of results from the multivariate analyses, we also conducted univariate association tests for each trait. The multivariate mixed model approach yielded 148 genome-wide significant single-nucleotide polymorphisms (SNPs) at a 10% false-discovery rate, compared to 129 and 73 significant SNPs in the PC and univariate analyses, respectively. Thus, explicit modeling of the correlation structure between fatty acids in a multivariate framework enabled identification of loci associated with variation in seed fatty acid concentration that were not detected in the univariate analyses. Ultimately, a detailed characterization of the loci underlying fatty acid variation can be used to enhance the nutritional profile of oats through breeding.

## INTRODUCTION

Oat (*Avena sativa* L.) is a nutrient-rich human and animal food source. Recent studies have revealed the numerous beneficial effects of oat consumption on human health, from reduction in cardiovascular diseases risk (Grundy *et al.* 2018) to cancer prevention (Meydani 2009). These positive health effects are likely due to oat’s unique nutritional profile, which differs markedly from that of other cereals, notably in the complement of essential amino acids, fatty acids, β-glucan, and phenolic compounds (Butt *et al.* 2008). In particular, lipids account for as much as 18% of the oat grain (Halima *et al.* 2015). Moreover, these lipids are predominantly composed of unsaturated fatty acids, rendering oats a healthful energy source in human and animal diets. In response to the growing awareness of oat’s health-promoting properties, nutritional quality has become a key target for oat breeders. Maintaining and/or optimizing lipid composition is an important component of these efforts (Valentine *et al.* 2011).

To date, researchers have primarily investigated the genetic basis of variation in fatty acid composition by mapping quantitative trait loci (QTL) in biparental populations (Hizbai *et al.* 2012). However, the use of such QTL in marker-assisted selection (MAS) is likely only effective if the parents in the mapping population exhibit trait variation and are closely related to lines in the relevant breeding populations (Snowdon and Friedt 2004). A genome-wide association study (GWAS) can mediate this limitation if the population in which the GWAS is conducted captures the genetic variation present in the target breeding population(s) (Lipka *et al.* 2015). Indeed, a GWAS can identify allelic diversity associated with trait variation in complex plant pedigrees when both genotypic and phenotypic data are available. In several agricultural crops, GWAS have provided insight into the genetic architecture of oil composition in crops such as maize (*Zea mays* L.; Cook *et al.* 2012; Li *et al.* 2013), rapeseed (*Brassica napus* L.; Gacek *et al.* 2017), and soybean (*Glycine max* (L.) Merr.; Zhang *et al.* 2018). To our knowledge, researchers have not yet conducted a GWAS of seed oil traits in oat though, small-scale surveys suggest substantial phenotypic diversity is present in existing oat germplasm (Saastamoinen *et al.* 1989; Leonova *et al.* 2008). Thus, we aim to uncover the genetic basis of fatty acid composition in cultivated oat as a foundation for improvement of oat nutritional quality through genomics-assisted breeding.

Historically, a GWAS considered a single-trait or multiple traits independently. To capitalize on the increasing quantity and complexity of phenotypic data, GWAS methods were developed to analyze multiple traits simultaneously while accounting for correlations between traits. The primary benefit of multi-trait or multivariate GWAS is that phenotypic correlations can increase statistical power to detect association signals relative to univariate methods (O’Reilly *et al.* 2012; Korte *et al.* 2012; Stephens 2013, Zhou and Stephens 2014). Several univariate approaches leverage trait correlations in the linear mixed model (or mixed linear model) framework. When there is a direct relationship between two phenotypes, their ratios can be used as the phenotype in a univariate GWAS. This approach lowers variance and conceptually targets a mediator of the phenotypes, such as an enzyme that converts one metabolite to another (Gieger *et al.* 2008). Another approach combines test statistics from univariate GWAS of each trait in order to detect genetic variants with pleiotropic effects (Yang *et al.* 2010). In addition, dimension reduction techniques, such as principal component analysis (PCA), can be used to derive transformed phenotypes as inputs for univariate GWAS (PC-GWAS). By combining signals across many PCs, PC-GWAS captures genetic signal associated with both single trait and pleiotropic effects, resulting in increased statistical power (Aschard *et al.* 2014). Finally, the association between a genetic variant and multiple traits can be directly modeled in a multivariate linear mixed model (O’Reilly *et al.* 2012; Korte *et al.* 2012; Zhou and Stephens 2014). We refer to the multivariate mixed model approach as multi-GWAS in order to differentiate this method from PC-GWAS which ultimately relies on a univariate linear mixed model for testing genetic associations.

Multivariate GWAS approaches are well-suited for genetically dissecting metabolic networks, because the abundances of precursors, intermediates, and products are likely correlated due to metabolic flux. Here, we present a GWAS of seed fatty acid content and composition in a diverse germplasm collection of cultivated oats. We compared conventional single-trait univariate GWAS on the abundance of ten fatty acids and their total with two multivariate GWAS methods: GWAS of principal components (PC-GWAS) and GWAS with a multivariate model that accounts for the abundance of all fatty acids simultaneously (multi-GWAS). These multivariate GWAS methods detected several novel loci associated with distinct effects on fatty acid composition and previously identified loci. Conservation of fatty acid metabolism across plant species (Figure 1; Li-Beisson *et al.* 2013) allowed us to preliminarily interpret our GWAS results despite the lack of a reference genome sequence for hexaploid oat. Aside from providing new insights into the genetic control of fatty acid composition in oat germplasm, our results suggest that multivariate approaches will be useful for improving the power of GWAS of compositional or other highly correlated traits that share a common genetic basis.

**Figure 1.**
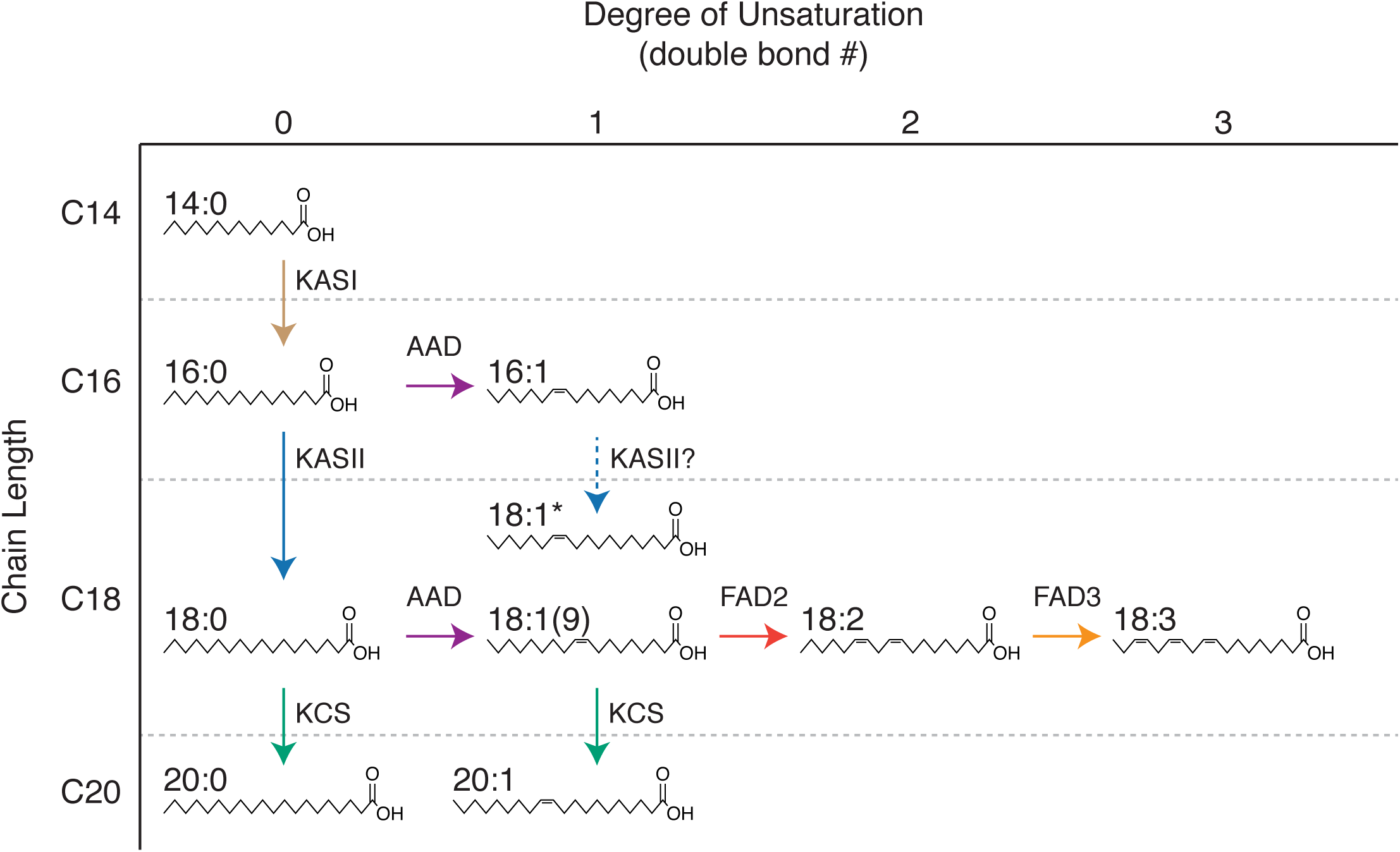
Inferred pathways of fatty acid synthesis and modification in oat seeds. Fatty acid abbreviations adhere to standard conventions, with chain length and degree of unsaturation (position[s] of double bond[s]) separated by a colon (for example, palmitic acid is denoted by 16:0). We detected two isomers of 18:1, 18:1(9) and another isomer, 18:1*, with unknown double-bond position (likely 18:1(11)). The 18:1(9) isomer was more abundant. Fatty acids up to 18 carbons in length are synthesized by a fatty acid synthase (FAS) complex. The nascent acyl chain is attached to acyl carrier protein (ACP) subunit of FAS and grows by two carbons per cycle through the action of distinct ketoacyl-ACP synthase (KAS) subunits of this complex (KASI and KASII are shown). Elongation is terminated either by a thioesterase that releases the fatty acid from ACP, or a double bond is introduced by an acyl-ACP desaturase (AAD) that typically acts with specificity for the Δ9 position and preference for C18 substrate. Thus, the initial fatty acid produced by FAS results from competition between one or more thioesterase and AAD isoforms. Subsequent elongation to ≥ 20C is catalyzed by the fatty acid elongase complex, in which the ketoacyl-CoA synthase (KCS) subunit determines chain length. Further desaturation is catalyzed by additional fatty acid desaturases (FAD2 and FAD3).

**Figure 2.**
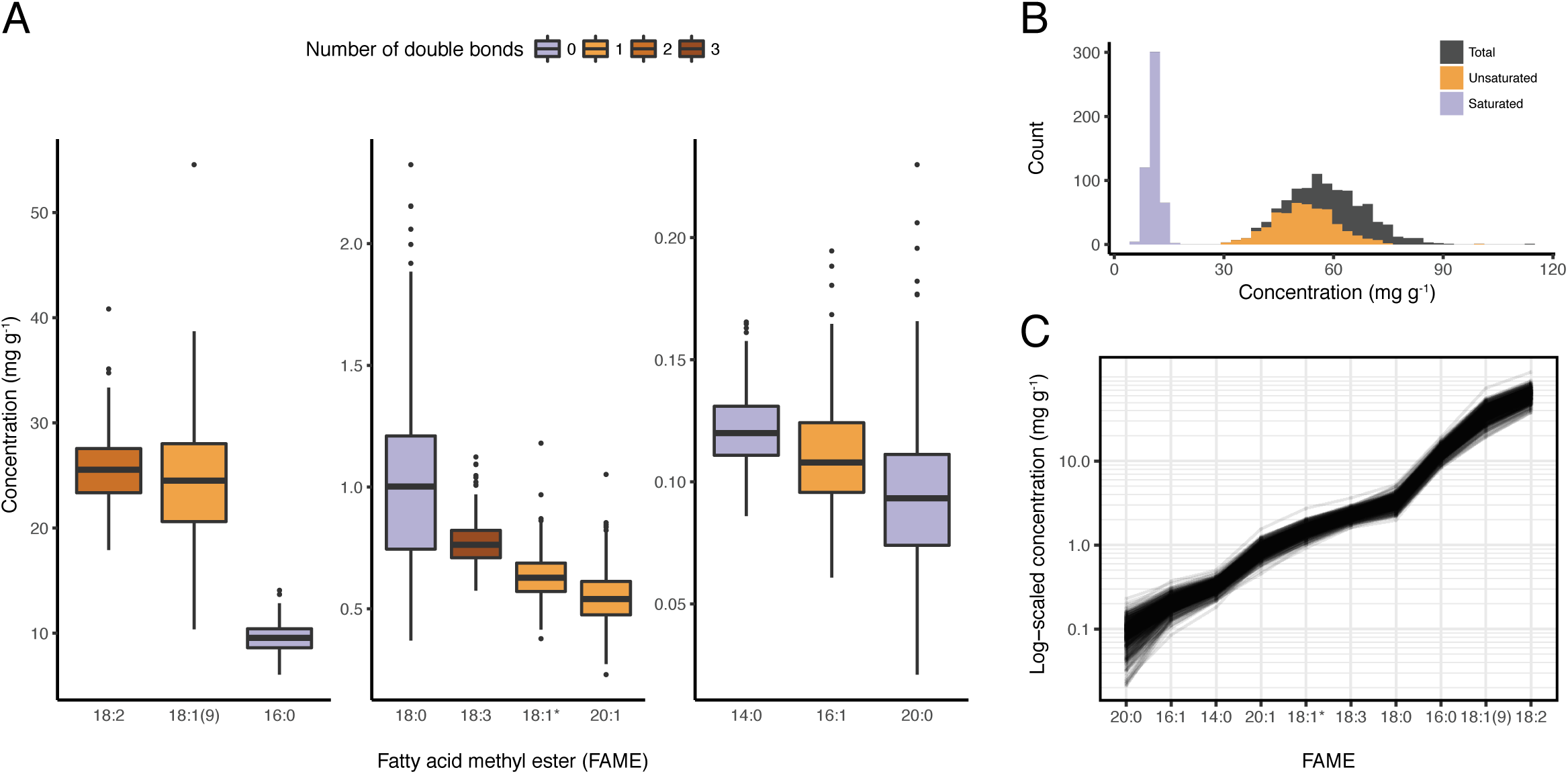
Variation in seed fatty acid concentration in a diverse oat panel. A. Box plots of the ten FAME best-linear unbiased predictor (BLUP) distributions measured in an oat diversity panel (*n* = 492). Compounds are divided into three groups based on mean concentration, each plotted with distinct y-axes. The color of the box denotes the number of double bonds in the corresponding FAME. B. Overlaid histograms of unsaturated, saturated, and total FAME BLUPs derived from the independent FAME measurements. C. The cumulative contribution of each FAME to total concentration plotted on a log-transformed scale.

## MATERIALS AND METHODS

### Germplasm

We assembled a 1,012 line diversity panel for oat, consisting of 391 inbred lines from the AFRI Core collection (Esvelt Klos *et al.* 2016), 219 inbred lines from Iowa State University (Newell *et al.* 2012), and 402 lines from a selection experiment conducted at Iowa State University composed of 20 inbred parents and 382 Cycle 1 or Cycle 2 individuals (Asoro *et al.* 2013). In 2014, the diversity panel was grown in an augmented field design with one replicate in each of two locations, with distinct soil types, near Ithaca, NY in 2014. Seeds were harvested, dried, and mechanically dehulled (Codema LLC, Maple Grove, MN, USA), as previously described (Montilla-Bascon *et al.* 2017). After visual inspection, any residual hull material was manually removed with a razor blade. Using the CDmean method of Rincent *et al.* (2012), which maximizes the genetic diversity relative to the larger source population, we selected a 500 line subset of the diversity panel for further analysis.

### Genotyping

A single seed from each line was planted in a greenhouse in Ithaca, NY. After four weeks, leaf tissue was collected for genotyping. DNA extraction and library preparation for genotyping-by-sequencing (GBS) were performed at Kansas State University (Poland and Rife 2012). Briefly, we extracted genomic DNA with the CTAB method as previously described (Esvelt Klos *et al.* 2016) and constructed 96-plex GBS libraries using the restriction enzymes *Pst*I and *Msp*I (Poland *et al.* 2012). Each library was then sequenced on a single Illumina HiSeq 2000 lane by the Cornell University Biotechnology Resource Center Genomics Facility.

Genotypes were called using the genotyping pipeline Haplotag, as this pipeline does not require a reference genome sequence (Tinker *et al.* 2016). Subsequently, single-nucleotide polymorphisms (SNPs) were mapped to a published consensus genetic linkage map generated from 15 bi-parental mapping populations (Chaffin *et al.* 2016, Bekele *et al.* 2018). We filtered SNPs using the following criteria: 1) biallelic; 2) minor allele frequency (MAF) > 2%; 3) site missingness < 60%; and 4) site heterozygosity < 10%. After initial SNP filtering, eight lines with more than 80% missing data and/or 10% heterozygous genotype calls were excluded from further analysis. To remove SNPs providing redundant information, we calculated Pearson’s correlation coefficient (*r*) for all pairwise combinations of SNPs, where missing data were coded as a fourth state. Only one SNP was retained in a group of SNPs with *r*_*2*_ = 1, leaving 29,320 SNPs. For the GWAS, we imputed missing data with the mean genotype.

### Kinship and population structure

To minimize the influence of missing data on pairwise relationship estimates, we estimated the relatedness matrix utilizing a 15,228 SNP subset (of the 29K filtered SNP set) with low missing data (<20%) with the *A.mat* function in the R package *rrBLUP* (Endelman 2011). Prior to conducting principal component analysis (PCA) with EIGENSTRAT (Price *et al.* 2006), we also removed SNPs that were not anchored to the genetic linkage map, resulting in a set of 12,585 SNPs.

### Fatty acid chemical analysis

Dehulled oat seeds were stored at −20°C prior to gas chromatography-mass spectrometry (GC-MS), performed at the Proteomics and Metabolomics Facility at Colorado State University. Oat seeds (300 mg) were ground to a fine powder in a 5 mL polypropylene vial containing a tungsten bead. A 6:3:1 (by volume) solution of cold methyl tert-butyl ether, methanol, and water (3 mL) was added to the vial prior to one hour of shaking at 4°C. Subsequently, 750 µL of water was added, the vial was vortexed again and centrifuged for 15 min at 4°C to induce phase separation. The upper organic layer, containing non-polar lipids, was transferred to a new glass vial and stored at −80°C until analysis.

Thawed samples were centrifuged at 3,750 rpm at 4°C for 10 min. The solvent was removed from 100 µL aliquots of each sample by nitrogen evaporation at room temperature. Once dried, 200 µL of toluene containing 1.25 mg/mL of internal standard (glyceryl triheptadecanoate), 200 µL of methanol, and 200 µL of 3N methanolic HCl, were added to the sample, and the mixture was incubated at 60°C for 1 h. Then, 1 ml of hexane and 300 µL of water were added to the cooled sample. After brief vortexing, the sample was centrifuged at 2,000 rpm for 5 min at 4°C.

One microliter of the upper hexane layer containing the fatty acid methyl esters (FAMEs) was injected onto a TG-WaxMS column (30 m x 0.25 mm x 0.25 µm, Thermo) in a TRACE 1310 Gas Chromatograph (Thermo) coupled to a Thermo ISQ LT GC-MS. The injector temperature and split ratio were 260°C and 30:1, respectively. The column was eluted with a constant flow of He carrier gas (1.2 mL min^-1^). The oven initial temperature was 200°C and held for 1 min, then increased to 260°C at 10°C min^-1^ and held for 3 min. Detection was completed under electron impact mode, with a scan range of 50-650 atomic mass units and scan rate of 5 scans s^-1^. Transfer line and source temperatures were 260°C and 230°C, respectively.

Quality control samples, consisting of pooled experimental samples, were injected after every 20 samples. Batches of 100 samples were prepared and analyzed simultaneously. Ion source, inlet liner, and septa were either cleaned or replaced after analysis of 300 samples. Standard curves were established for 16:0, 18:0, 18:1(9), 18:2, and 18:3. Two standard FAME mix samples (Nucheck GLC-85 and Sigma 47080-U) were used to confirm the retention times. In the case of 18:1, two peaks were consistently observed. The major peak in all samples corresponded to 18:1(9) based on retention time and authentic standards. The minor peak we designate 18:1* since assignment of double bond position was not possible based on spectra or authentic standards (Figure S1), although 18:1* is most likely the methyl ester of cis-vaccenic acid, 18:1(11) (Cahoon *et al.* 1998). Data processing was completed with Chromeleon 7 software (Thermo).

### Phenotypic data analysis

Ten fatty acids were measured as their respective FAMEs from the 500-line diversity panel (see Figure 1 for fatty acid structures and abbreviations). We observed 19:0 and 15:0 in several samples, but as values of these compounds were largely below the limit of detection (LOD), we excluded 19:0 and 15:0 from analysis. Samples that had levels below the LOD (zero values) for 14:0 (n=1), 16:1 (n=2), 20:0 (n=50), and 20:1 (n=1) were replaced with a uniform random variable between zero and the LOD, where the minimum non-zero value was used a proxy for LOD (Lipka *et al.* 2013). The ten individual FAME traits and total FAME were inspected for outliers in ASReml-R 3.0 (Butler *et al.* 2009) by examination of the Studentized deleted residuals (Kutner *et al.* 2004) from linear mixed models fitted with environment, block, line, and GC-MS batch as random effects and heading date as a fixed effect. To assess the effect of the two environments on FAME concentrations, we calculated pairwise Spearman’s rank correlations (ρ) and performed hierarchical clustering on all trait-environment combinations.

Given that fatty acids comprise a chemical family, we also considered the relative contribution of each member to total concentration. Such a compositional analysis imposes a unit-sum constraint, i.e. all proportions must sum to one, which, in turn, induces spurious negative correlations between variables (Aitchison 1983). Therefore, to relieve the constraints of this sample space (the unit simplex), we employed Aitchison’s log-contrast transformation. Specifically, we divided each FAME by an arbitrarily selected compound (here, 18:1*) prior to calculating pairwise correlations (Wei and Simko 2017).

As the FAME concentrations were approximately normally distributed, we used the raw phenotype values to estimate a best linear unbiased predictor (BLUP) for each line in ASReml-R. Our BLUP model (Equation 1) incorporated environmental and technical covariates, as well as variation in heading date (Gilmour *et al.* 2008):

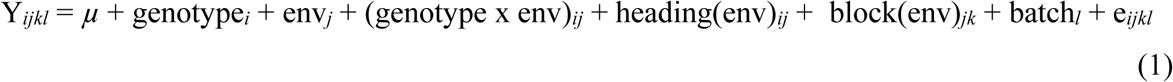

in which Y_*ijkl*_ is a plot-level average; *µ* is the grand mean; genotype_*i*_ is the effect of the *i*th genotype; env_*j*_ is the effect of the *j*th environment; (genotype x env)_*ij*_ is the effect of the *i*th genotype in the *j*th environment; heading(env)_*ij*_ is the effect of days to heading of the *i*th genotype within the *j*th environment; block(env)_*jk*_ is the effect of the *k*th block within the *j*th environment; batch_*l*_ is the effect of the *l*th GC-MS batch; and, e_*ijkl*_ is the random error term, assumed to follow a normal distribution with mean zero and variance *σ*^*2*^. Only heading(env)_*ij*_ was fitted as a (continuous) fixed effect; all other terms were fitted as random effects.

Variance component estimates from the fitted model were used to calculate line-mean heritability for each trait (*hl*^*2*^ as per Holland *et al.* (2002) and Hung *et al.* (2012)). Standard errors of the line-mean heritabilities were estimated using the delta method (Holland *et al.* 2002).

To identify axes of variation in the FAME data set, we performed PCA on the centered and scaled (to unit-variance) BLUPs using the R package *pcaMethods* (Stacklies *et al.* 2007). We extracted the principal component (PC) loadings to determine the contribution of each compound to each PC.

### Network analysis with a Gaussian graphical model (GGM)

To assess the relationships between BLUPs for the 11 FAME measures (ten individual, plus the total FAMEs), hereafter referred to as FAME BLUPs, we first estimated pairwise Pearson’s correlation coefficients (*r*) between each trait with the R package *Hmisc* (Harrell 2018). We next constructed a Gaussian graphical model (GGM). As GGMs reflect the conditional dependencies between variables, they are more likely to capture causality and precursor/product relationships in metabolic networks relative to standard correlation analyses (Krumsiek *et al.* 2011). We employed the R package *ppcor* for calculation of partial correlations and significance testing (Kim 2015). An edge was drawn between two metabolites if the partial correlation was significant after applying a Bonferroni correction ((*α* = 0.05 / ((*m*(*m*-1)) / 2)), where *m* is equal to the number of metabolites (*m* = 10). We visualized both the *r* and *pr* networks with the R packages *network* and *ggplot2* (Butts 2008; Wickham 2016).

### Genome-wide association studies (GWAS)

We leveraged the correlations between FAME traits in a multivariate GWAS with 492 lines and 29,320 SNPs. Specifically, we fitted a multivariate linear mixed model, accounting for population structure and relatedness, with all ten FAME concentrations in GEMMA (Zhou and Stephens 2012). To determine the optimal number of genetic principal components (PCs) based on the genotype matrix to include in the GWAS model, we used the Bayesian information criterion (BIC), comparing models with zero to five PCs (Schwarz 1978; Lipka *et al.* 2012). To minimize deviation from multivariate normality, we quantile transformed each phenotype to a standard normal distribution (Stephens 2013). Specifically, the raw BLUPs were rank normalized using the *qnorm* function of R and scaled to unit variance. To inform our multivariate analysis, we also conducted univariate association analyses on each of the non-transformed FAME BLUPs and the PCs derived from singular value decomposition of the FAME BLUPs data matrix (PC-GWAS, see *Phenotypic data analysis*). In each association test, we employed the univariate analog of our multivariate GWAS model in GEMMA. To account for multiple testing, we computed a Bonferroni correction (*α* = 0.05 / 29,320 = 1.7 x 10^-6^). Alternatively, we controlled the false-discovery rate (FDR) at 5% and 10% (Benjamini and Hochberg 1995) using the R package *qvalue* (Storey *et al.* 2019).

Pairwise LD between SNPs was estimated using squared allele frequency correlations (*r*^2^), excluding missing and double heterozygous genotypes (Pritchard and Przeworski 2001). We defined the *significant SNP set* as the union of SNPs identified as significantly associated with a phenotype at a 10% FDR in any of the univariate or multivariate analyses. To remove likely false positives and focus on GWAS signals with the most statistical support, we further analyzed only those significant SNPs in LD (*r*^2^ > 0.5) with at least two other significant SNPs.

The amount of phenotypic variation explained by each SNP in the univariate analyses was approximated using a likelihood-ratio-based *R*^2^ statistic (*R*^2^_LR_), as defined by Sun *et al.* (2010) (Table S2). The *R*^2^_LR_ value of models with or without a significant SNP was calculated using the maximum log-likelihood of the model of interest fitted in GEMMA and the maximum log-likelihood of an intercept-only model fitted with the *lm* function in R. For the SNPs detected to be significant in the multi-GWAS, we approximated the relative contribution of each trait to the multivariate signal by calculating *R*^2^_LR_ for each SNP in each of the ten respective univariate models developed for the quantile-transformed FAME BLUPs (Table S3).

To qualitatively summarize the effects of SNPs identified as statistically significant in the multi-GWAS on individual FAME levels, we compared the BLUP phenotypic means of the distinct homozygous classes (major and minor) and conducted a pairwise t-test (*pairwise.t.test* function in R) for each SNP. Phenotypes were mean centered to zero and standardized to unit variance prior to testing. We excluded heterozygous and imputed genotypes.

### Data availability

All genotype and phenotype data are freely available from the Triticeae Toolbox/Oat database (https://triticeaetoolbox.org/oat/).

## RESULTS

### Genotyping and population structure

We assembled a panel of 1,012 oat lines from three existing germplasm collections. This panel consists of inbred lines from around the world, some of which were selected based on variability of beta-glucan, including selections from a beta-glucan improvement program (see *Materials and Methods*). We genotyped this panel with genotyping-by-sequencing (GBS), a reduced-representation sequencing technique (Poland and Rife 2012). After genotype calling with the non-reference pipeline Haplotag (Tinker *et al.* 2016), we identified an optimal and diverse subset of 500 lines using the CDmean method of Rincent *et al.* (2012). Within this subset, we applied standard quality control filters, resulting in a data set of 29,320 SNPs and 492 individuals in the diversity panel. Of these 29,320 SNPs, 19,558 (67%) were anchored to the consensus genetic map (Chaffin *et al.* 2016; Bekele *et al.* 2018). Principal component analysis (PCA) on a reduced set of 12,585 SNPs (see *Materials and Methods*) revealed minimal structure with respect to line origin (Figure S2).

### Variation in seed fatty acid content

Oats are especially high in healthful mono- and di-unsaturated fatty acids, relative to other cereals. To investigate variation in nutritional quality in oat, we measured the levels of ten FAMEs derived from non-polar seed lipids in the 492-line diversity panel grown in two environments (referred to as ENV1 and ENV2). In both environments, 18:2, 18:1(9), and 16:0 were the most abundant FAME species (Figure S3C), in accordance with previous reports (Banaś *et al.* 2007; Leonova *et al.* 2008). In particular, 18:2 and 18:1(9) accounted for >70% of the total FAME content in all lines (Figure S3F).

FAME concentrations were consistently higher in ENV1 compared to ENV2 (pairwise t-tests detected significant differences for all traits except 18:3). To assess the impact of this consistent environmental difference on the statistical relationships between traits with respect to environment, we calculated Spearman’s rank correlations between each trait-environment combination. Hierarchical clustering revealed grouping with respect to environment and to a lesser extent compound (Figure S3B). For example, the three FAMEs in highest abundance, 18:2, 18:1(9), and 16:0, clustered together in each environment, along with 20:1. In addition, we observed clustering between 20:0 and 18:0, and 16:1 and 18:1*, respectively.

Having observed that trait-environment clusters were predominately defined by environment (and secondarily by compound), we were interested in assessing the differential effect of environment on FAME composition, defined as the percent contribution of each FAME to total concentration. We observed significantly higher proportions of 14:0, 18:0, 18:2, and 20:0, and lower proportions of 18:1(9), 18:3, and 20:1 in ENV1 as compared to ENV2 (Figure S3F; *p*-value of pairwise t-test < 0.05). The proportions of other compounds, and the sum of the saturated FAMEs, were comparable across environments.

Assessing the correlation structure in this compositional sample space necessitated relieving the unit-sum constraint imposed by the fact that proportions must sum to one. After applying the log-contrast transformation (Aitchison 1983) to FAME concentrations, i.e. taking the logarithm of each FAME divided by arbitrarily selected FAME (18:1*), trait rather than environment defined clustering (Figure S3D-E). Yet, we still observed positive correlations between all trait pairs, excluding the pairwise comparisons that featured 16:1.

Consistent effects of environment on concentrations across lines corresponded to high line-mean heritabilities for all FAMEs, ranging from 0.67 to 0.91 (Table S1). Thus, we conclude that genetic determinants have a greater impact than environmental factors on variation in FAME concentrations between lines.

### Correlation networks

Using data from the two environments, we calculated BLUPs of each FAME for the 492 lines in the diversity panel. Consistent with our analysis of the raw phenotypes, we observed only positive correlations between FAME BLUPs (Figure S3), translating to a dense, highly connected network (Figure 3A).

**Figure 3.**
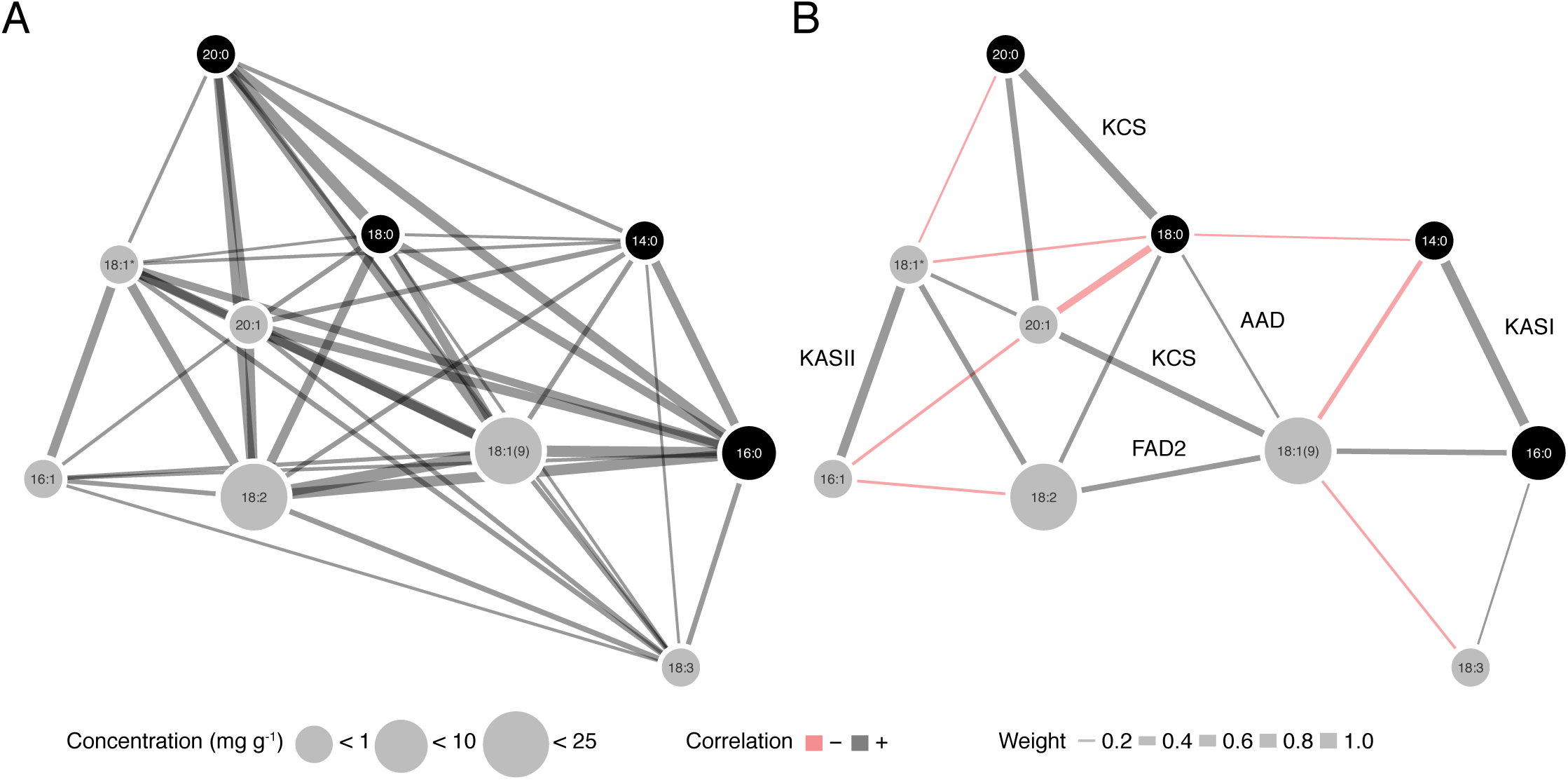
Fatty acid methyl ester (FAME) correlation networks. A. A network constructed from Pearson’s correlations (*r*) between FAMEs. B. An analogous network constructed from pairwise partial correlations (*pr*) with edges corresponding to biosynthetic steps annotated (See Figure 1 for abbreviations). In both A and B, an edge was drawn between compounds if the pairwise *r* or *pr* value was significant given a threshold of α = 0.05, after applying a Bonferroni correction for multiple-testing. Edge width is proportional to the magnitude of the correlation, and edge color indicates a positive or negative correlation. Node size corresponds to the mean concentration (mg g^-1^), with compounds grouped into three categories: less than 1, 10, and 25 mg g^-1^. Node color differentiates saturated from unsaturated FAMEs.

As FAMEs participate in shared biochemical pathways, we hypothesized that many of the observed correlations reflected indirect rather than direct interactions. Our rationale was that metabolic networks are generally sparse due to the limited reactivity of organic compounds and stepwise nature of metabolic pathways. Specifically, fatty acid biosynthetic pathways exhibit this property (Li-Beisson *et al.* 2013). Therefore, akin to Krumsiek *et al.* (2011), we constructed a Gaussian graphical model (GGM) to account for the conditional dependencies between FAME BLUPs. We estimated partial correlations (*pr*s), defined as the correlation between the residuals of two compounds after accounting for the other *n*-2 compounds. As hypothesized, the *pr* distribution was shifted toward zero relative to its *r* counterpart (Figure S4). This translated to a far sparser network, such that every node was no longer connected to nearly every other node by an edge, and to the introduction of negative correlations (Figure 3B). Notably, the large positive correlations between compounds with a difference of two carbons, but the same number of double bonds (e.g. 18:0 and 20:0), persisted in the *pr* network. While the significant pairwise *pr*s were not restricted to only precursor/product relationships in fatty acid biosynthesis, some of the strongest correlations in this network are between such pairs (Figure 3B).

### Decomposition of FAME variance

To identify the contribution of each fatty acid to the major axes of variation in the phenotypic data set, we performed PCA on the centered and scaled BLUPs for the ten FAMEs (Figure S5). All FAMEs exhibited negative loadings along the first PC (explaining 56.8% of the total variance), which tracked with variation in total fatty acids (adjusted *R*^2^ of total with PC 1 = 0.952; Figure S5). Thus, most of the variation in individual FAME concentrations can be explained by variation in fatty acid totals. The second PC (explaining 15.2% of variance) primarily distinguished saturated from unsaturated FAMEs, whereas the third and fourth PCs differentiated 14:0 and 18:3 from all other FAMEs, respectively. While 20:1 contributed disproportionately to both the fifth and sixth PCs, these two PCs were less obviously interpretable.

### Genetic mapping for fatty acids

We employed several mapping approaches to capitalize on the correlations between traits. First, we performed a multivariate GWAS of all ten FAME concentrations simultaneously (multi-GWAS). Second, we conducted GWAS with each PC derived from decomposition of the phenotypic data matrix (of dimension *n* x *m* = 492 lines x 10 traits) as phenotypes (PC-GWAS). Third, we implemented univariate tests of association for each FAME separately, as estimated by their BLUPs. To account for population structure, we included the first three PCs of the non-imputed genotype matrix as covariates in all of our GWAS models. These PCs were selected based on application of the BIC model selection procedure in the multi-GWAS, and explained 6.7, 4.3, and 3.2 percent of the total variance, respectively.

In the multi-GWAS, a total of 25 SNPs were significant after accounting for multiple testing with a 5% Bonferroni significance threshold (*p*-value ≤ 1.7 x 10^-6^). We also considered SNPs passing the two less conservative significance thresholds of genome-wide (FDR of 5% and 10%) At these thresholds, 93 and 148 SNPs were detected, respectively.

After applying a 10% FDR threshold independently in each association test, and combining results across all ten PCs, 129 SNPs were significantly associated with FAME variation in the PC-GWAS (Figure 4C and Figure S6). Approximately half of these 129 SNPs (66 SNPs) were represented in the multivariate 10% FDR significant set (Figure 4). The majority of SNPs uniquely identified in PC-GWAS were associated with PC10 and not anchored to a linkage group. This PC explained only 0.4 percent of total variance across the ten FAMEs, with 18:1(9) weighted most heavily along this PC (Figure S5).

**Figure 4.**
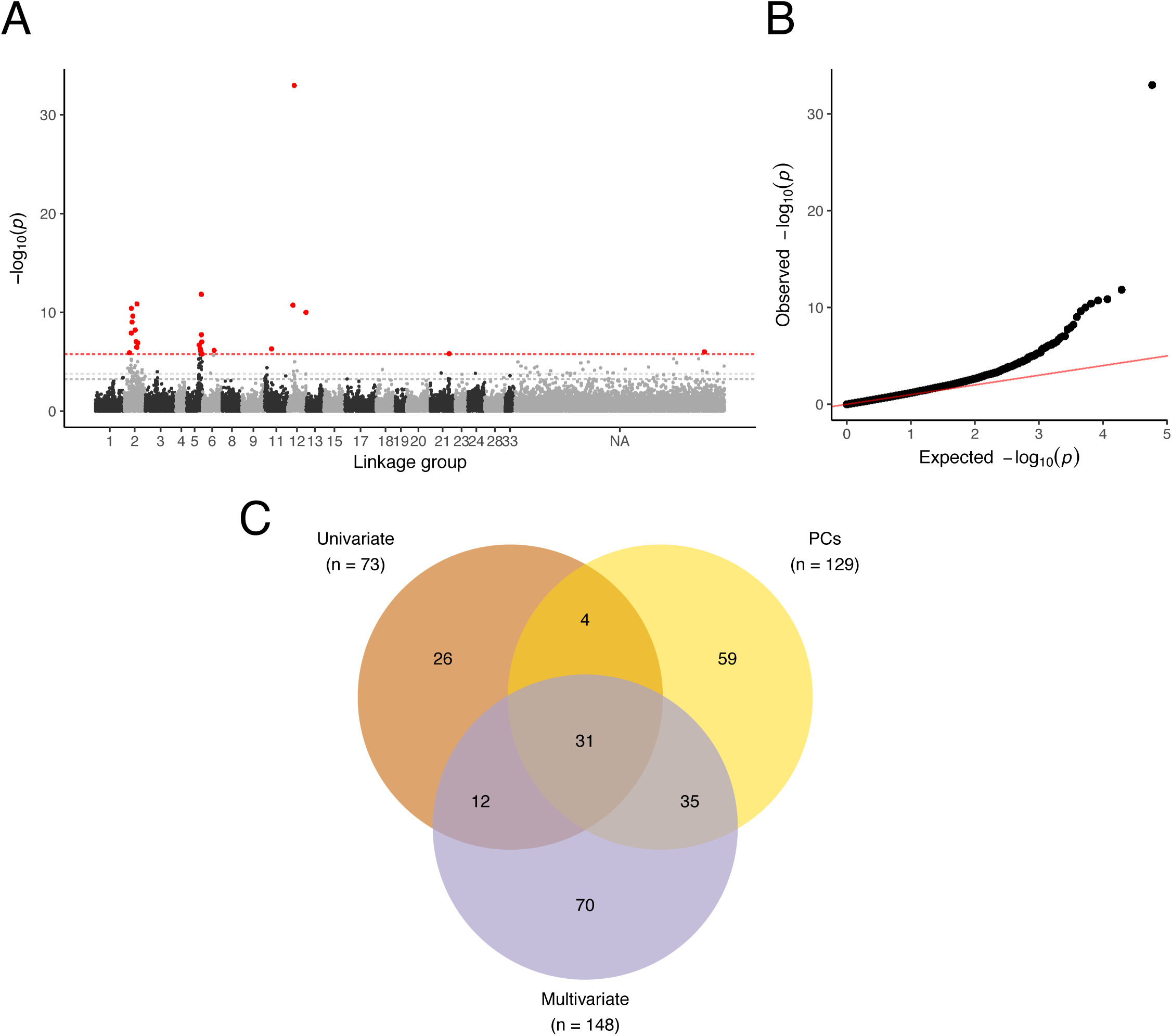
Multivariate genome-wide association study (multi-GWAS). A. Negative log_10_ *p*-values from a multi-GWAS of ten FAME BLUPs plotted against genetic position in the consensus linkage map (Chaffin *et al.* 2016; Bekele *et al.* 2018). Dotted lines denote the three significance thresholds considered, with a Bonferroni-corrected threshold of 5% in red, and 5 and 10% false-discovery rate (FDR) thresholds in light and dark gray, respectively. Markers with a *p*-value passing the Bonferroni threshold are shown in red. B. A quantile-quantile plot of the multi-GWAS *p*-values. C. Venn diagram comparing significantly associated SNPs (at a 10% FDR) from combined univariate analyses of ten individual and total FAME untransformed BLUPs (Univariate), combined univariate analyses of ten principal components (PCs) of the ten FAME BLUP data matrix, and multi-GWAS of the ten FAME quantile-transformed BLUPs (Multivariate).

In the univariate GWAS of the ten FAMEs and total FAME concentration, we identified 73 significantly associated SNPs at a 10% FDR (Figure 4C and Figure S7). Thus, univariate GWAS had the least power to detect significant associations. Furthermore, the majority of these SNPs were identified by either multi- or PC-GWAS (47 SNPs). The uniquely identified SNPs in the univariate GWAS were primarily associated with less abundant fatty acids, e.g. 18:0, 20:0, and 20:1. To provide a more apt comparison between the univariate and multivariate analysis, which employed quantile-transformed traits, we also performed the univariate GWAS with quantile-transformed traits. This analysis identified 62 SNPs at a 10% FDR (Figure S8). All but two of these SNPs were also found in GWAS of the non-quantile transformed BLUPs, suggesting that deviations from normality in the untransformed BLUPs had minimal influence on the results (Figure S8; Table S2).

### Integration of GWAS results

To focus on GWAS signals with the most statistical support, we only considered significantly associated SNPs with *r*^2^ > 0.5 with at least two other significant SNPs (at a 10% FDR threshold) across all analyses. This high confidence SNP set consisted of a 152 SNP subset of the 237 significant SNPs. The majority of SNPs in the high confidence SNP set (101 SNPs) spanned eight of the 21 consensus linkage groups, while 51 SNPs were unmapped (Figure 5). In the absence of a reference genome sequence, we used pairwise LD between significantly associated SNPs to putatively define the boundaries of independent signals (Figure 5). We further refined these boundaries such that SNPs within a signal interval had concordant estimated effects on the FAME abundances. Here, we assumed that if SNPs were in LD with the same causal locus they should exhibit correlated effects on the ten FAME traits. To interpret SNP effects, we inspected the mean difference between homozygous genotypic classes excluding lines with missing data or heterozygous genotypes (Figure 6). While this simple summary does not account for population structure and unequal relatedness, it provides qualitative insight into the effects of SNPs on individual FAMEs, which are obscured by multivariate GWAS.

**Figure 5.**
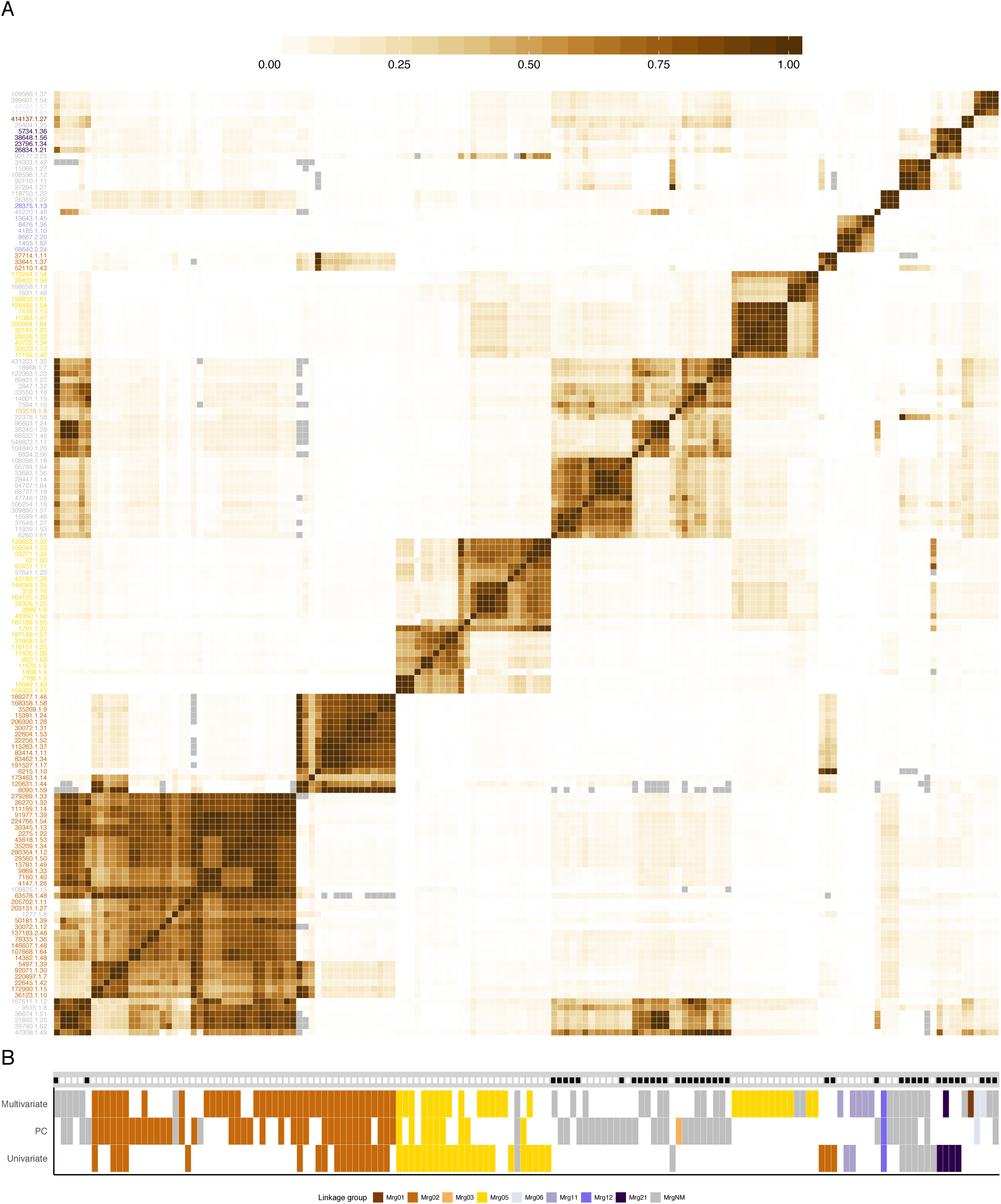
Linkage disequilibrium between markers associated with fatty acid methyl ester (FAME) variation identified in multivariate, principal component (PC), and univariate analyses. A. Linkage disequilibrium (LD), defined as the squared allele frequency correlation coefficient (*r*^*2*^), between markers identified as significantly associated with phenotypic variation at a 10% false-discovery rate (FDR) threshold in any of the multivariate, PC, or marginal analyses. (Here, we show only those significant markers that were in LD with at least two other significantly associated markers (see *Materials and Methods*). Single-nucleotide polymorphisms (SNPs) are ordered by hierarchical clustering, with abbreviated SNP name on the y-axis colored by linkage group assignment (see B) in the consensus genetic map (Chaffin *et al.* 2016; Bekele *et al.* 2018). Gray boxes denote missing values (see *Materials and Methods*). B. An incidence matrix of results from the multivariate, PC, and univariate association analyses. The x-axis mirrors that of A, with each vertical tract corresponding to the same SNP pictured in A. A colored rectangle indicates that the SNP was identified as significant at an FDR threshold of 10%, with the PC and Univariate tracks representing the union of significant SNPs across all ten and 11 traits, respectively. The color of the rectangle corresponds to the linkage group. A black dot along the gray track indicate that the SNP has a minor allele frequency < 5%.

**Figure 6.**
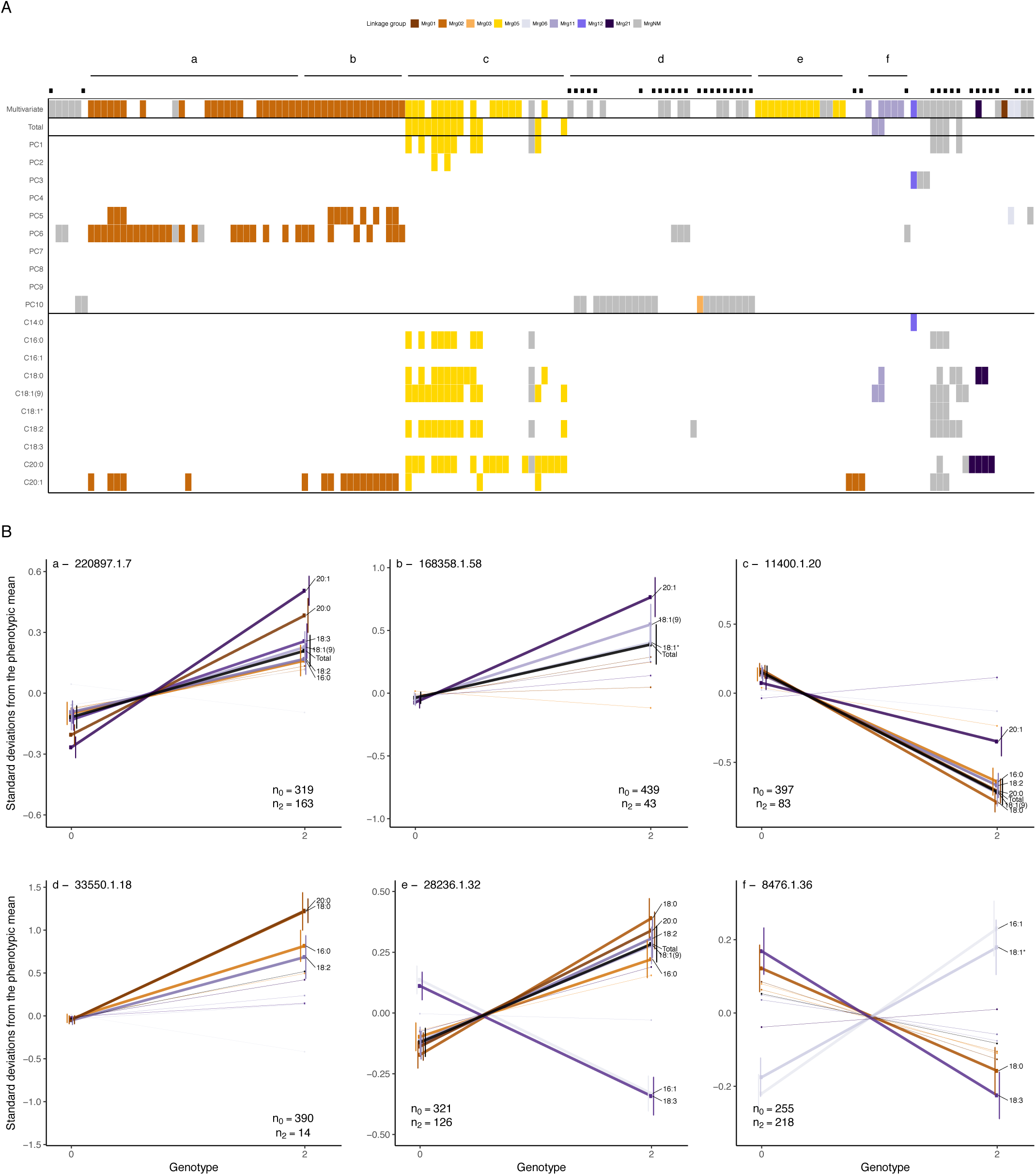
Comparison of multivariate, principal component (PC), and univariate genome-wide association study (GWAS) results. A. An incidence matrix of results from the multivariate, PC, and univariate GWAS. SNPs are ordered by hierarchical clustering of pairwise linkage disequilibrium (LD) estimates, as in Figure 5. A colored rectangle indicates that a SNP was significant in the given test at a 10% false-discovery rate (FDR) threshold, with color corresponding to a linkage group in the consensus genetic map (Chaffin *et al.* 2016; Bekele *et al.* 2018). Presence of a black dot indicates that the SNP has a minor allele frequency < 5%. Labeled (*a-f*) line segments span the SNPs in six LD clusters, defined by visualization of the pairwise LD matrix (see Figure 5). B. Mean phenotypic differences between distinct homozygote classes at a significantly associated SNP within each of the six LD clusters (*a-f*) defined in A. Genotype is plotted on the x-axis, with the mean number of phenotypic standard deviations away from the mean on the y-axis. Each centered and unit-variance scaled FAME BLUP (including total) is plotted, with warm colors for saturated and cool colors for unsaturated FAMEs. Total is indicated by a black line. The bold lines indicate that the pairwise t-test between genotype class means was significant at α = 0.05 after correcting for multiple testing with a Bonferroni correction. To simplify the plots, only traits with significantly different means between genotypes are labeled. As lines with missing genotypes are excluded from these calculations, we present the genotype counts in the bottom right or left-hand corner of each plot.

Using these criteria, we identified six main clusters, consisting of 124 SNPs in the high confidence SNP set (*a*-*f* in Figure 6). Five of the clusters localized to three linkage groups: Mrg02, Mrg05, and Mrg01, while one cluster was not anchored to the linkage map (Figure 6). The remaining 28 markers formed smaller clusters that were either unanchored or primarily supported by SNPs with MAF < 5% (Figures 5, 6, and S9). One notable small cluster is anchored at 36.0 cM on Mrg12 by a SNP with a very small *p-*value (1.9 x 10^-11^). Although the two other SNPs in this cluster were unanchored, the overall peak SNP in the multi-GWAS (*p* = 1.1 x 10^-33^) was mapped to a nearby position (40.9 cM) on this linkage group but excluded from the high confidence SNP set by our *r*^2^ clustering criteria. This SNP and the LD cluster SNPs were associated with 14:0 and PC3 (which exhibits a large 14:0 loading, Figure S5B), suggesting at least one gene in this Mrg12 region underlies variation in seed 14:0 content.

We observed two primary clusters in Mrg02, spanning 37.5 to 107.5 cM (cluster *a*) and 67.4 to 85.3 cM (cluster *b*), respectively. These large intervals are consistent with previous reported of long-range LD in this linkage group (Chaffin *et al.* 2016; Esvelt Klos *et al.* 2016; Sunstrum *et al.* 2019). We suspect that long-range LD, coupled with a causal locus in Mrg02, inflated test statistics for SNPs in Mrg02 resulting in the over representation of this linkage group in the high confidence SNP set. The peak SNPs from the two clusters (*a* and *b*) had markedly different minor allele frequencies (0.33 vs. 0.09; Figure 6B, *a* and *b*). As such, the *r*^2^ for these two SNPs is constrained to a maximum value of *r*^2^ = 0.19, reflecting the sensitivity of *r*^2^ to differences in allele frequency (VanLiere and Rosenberg 2008). Therefore, while weak LD between these SNPs (*r*^2^ = 0.14) resulted in assignment to different clusters by our criteria (Figure 5), the two clusters are likely tagging the same locus, or different alleles at the same locus. Note that without a physical map we cannot identify signal intervals with certainty. Furthermore, SNPs within both clusters exhibit similar patterns of phenotypic associations. In both clusters, SNPs were associated with increases in low concentration 20:1 and the biosynthetically-related, and abundant 18:1(9) (Figure 6B, *a* and *b*). SNPs in cluster *a* were also associated with higher levels of 20:0, 18:2, 18:3, and 16:0, while SNPs in cluster *b* also exhibited association with higher levels of 18:1*. Notably, in the univariate GWAS, the SNPs within these two clusters were only associated with 20:1, highlighting the increased statistical power of the multi- and PC-GWAS approaches to detect markers in LD with causal polymorphisms underlying correlated traits.

Two clusters were also observed in Mrg05, mapping to non-overlapping intervals between 124.1 and 135.5 cM (cluster *c*) and 114.7 to 122.7 cM (cluster *e*). The minor alleles in cluster *c* were associated with significant decreases in most FAMEs as well as total concentration. All three approaches (univariate, PCs, and multivariate) detected significant SNPs in the interval associated with cluster *c*. In contrast, cluster *e* was only detected by the multi-GWAS. In this cluster, the minor allele was associated with higher total FAMEs and concentrations of the most abundant FAMEs, as well as, decreases in 18:3 and 16:1. The correlated estimated effects of SNPs in cluster *e* on 16:1 and 18:3 were not expected *a priori* based on known biochemical pathways (Figure 1) or the observed correlation networks (Figure 3). To further probe the major drivers of the multi-GWAS signals, for each significant SNP in the multi-GWAS we calculated *R*^2^_LR_ in the univariate models for each trait. The difference between *R*^2^_LR_ with or without a given SNP included in the model, Δ*R*^2^_LR_, provides an approximation of phenotypic variance explained by that SNP. Thus, we used univariate Δ*R*^2^_LR_ of each component trait as a means of qualitatively dissecting the multi-GWAS signal (Table S3). By this analysis, detection of cluster *e* SNPs associate with 18:3 variation, despite the fact that we failed to detect significant associations in the univariate GWAS. Thus, it appears that unanticipated correlations between 18:3 and other fatty acids, such as 16:1, underlie the enhanced sensitivity of the multi-GWAS model.

The cluster identified on Mrg11 spanned 3.7 to 8.8 cM. This cluster (*f*) contained two SNPs identified in the univariate analyses of total and 18:1(9) FAMEs, and four SNPs uniquely identified in the multi-GWAS. The main effects associated with the peak SNP in this cluster were increased 16:1 and 18:1* and decreased 18:0 and 18:3 (Figure 6B). However, other SNPs in this cluster were associated with decreases in the majority of FAMEs, including total concentration (Figure S9). Thus, as with cluster *a* and *b*, the SNPs of cluster *f* may be tagging multiple alleles or loci and our interpretation is limited by the present lack of a genome sequence in hexaploid oat.

The final cluster (*d*) consisted of 29 SNPs of low minor allele frequency (< 0.05). The only mapped SNP in cluster *d* was located at 80.3 cM on Mrg03 of the consensus map and a significant association of this SNP was only detected in GWAS of the last PC. SNPs in this cluster were predominantly associated with higher levels of saturated fatty acids (16:0, 18:0, and 20:0).

## Discussion

We report the first GWAS of seed fatty acid composition in oat. Our results reveal several large effect loci underlying this important seed quality trait. At least four loci on four linkage groups contribute to variation in the content and composition of seed fatty acids in oat. Ongoing efforts to sequence the ~12.5 Gb hexaploid oat genome will likely enable higher resolution mapping of these GWAS hits. On the other hand, inability to define distinct GWAS support intervals may, in part, reflect the complex genetics of oat germplasm. Specifically, translocations and other chromosomal rearrangements are common in oat germplasm (Jellen *et al.* 1994), and these are thought to underlie distortions of LD within the consensus linkage groups as observed here and in other studies (Chaffin *et al.* 2016; Esvelt Klos *et al.* 2016; Sunstrum *et al.* 2019).

Previous studies used linkage analysis in biparental populations to map QTL for fatty acid composition or oil amount in oat seeds. Kianian *et al.* (1999) mapped QTL for oil content in two RIL populations, “Kanota” x “Ogle” (KO) and “Kanota” x “Marion” (KM). In both the KO and KM populations, alleles of a gene encoding Acetyl-CoA Carboxylase, *ACCaseA*, were found to associate with a QTL accounting for up to 48% of phenotypic variance in these populations (Kianian *et al.* 1999). This gene is likely to underlie seed oil content because its respective enzyme catalyzes the synthesis of Malonyl-CoA, the major substrate of fatty acid synthase. Colocalizing oil content QTL were later discovered in additional populations: “Ogle” x “MAM17-5” (Zhu *et al.* 2004), “Aslak” x “Matilda” (Tanhuanpää *et al.* 2012), and “Dal” x “Exeter” (Hizbai *et al.* 2012). *ACCaseA* is tightly linked with the RFLP marker *cdo665B*, which is placed at a position of 135.5 cM on Mrg05 of the consensus map of Chaffin *et al.* (2016). Thus, these QTL, and their putative causal locus, *ACCaseA*, likely correspond to cluster c (Figure 6) in our current work.

Clusters *a* and *b*, mapping to overlapping intervals on Mrg02 (Figure 6), may correspond to two oil content QTL discovered in the “Dal” x “Exeter” population, *oPt-11790* and *oPt-16384* (Hizbai *et al.* 2012). Another QTL identified in Hizbai *et al.* 2012, *oPT-17489*, colocalizes with cluster *f* on Mrg11 in our work (Figure 6). Mechanistically, the effects associated with cluster *b* and *f* may be attributed to changes in acyl-ACP desaturase (AAD) activity. In the case of cluster *b*, an increase in AAD activity could lead to increases in both 18:1(9) and 18:1* as well the elongation of 18:1(9) to 20:1. The correlated change in 16:1 and 18:1* (assumed to be 18:1(11)), on the other hand, would be consistent with an altered substrate specificity for AAD. In plants, AAD typically has a strong preference for 18:0-ACP as its substrate, and the conversion of 16:0-ACP to 16:1-ACP is typically a minor side reaction (Cahoon *et al.* 1997). In some cases, such as the seeds of cat’s claw (*Doxantha unguis-cati* L.), AAD isoforms with preference for 16:0-ACP as a substrate can lead to substantial production of 16:1(9) and its elongation product, 18:1(11) (Cahoon *et al.* 1998). While these interpretations are speculative, they illustrate how the increased sensitivity of multivariate GWAS models can link relatively subtle perturbations of metabolic networks to their genetic and biochemical bases.

One objective of this study was to identify loci associated with variation in 18:3 content, as these omega-3 fatty acids are a large determinant of oil quality. Increased consumption of omega-3 fatty acids is associated with improved cardiovascular health, although there is uncertainty regarding the benefits of consuming omega-3 fatty acids relative to omega-6 fatty acids, such as 18:2 (Ludwig *et al.* 2018). In oat seeds, the very low abundance of 18:3 relative to 18:2 in germplasm (Figure 2A) renders enhanced 18:3 content an unlikely breeding target for improving the healthfulness of oats. On the other hand, 18:3 is exceptionally prone to oxidation, resulting in rancidity. Therefore, decreased 18:3 content may improve postharvest stability (Zhou *et al.* 1999). In univariate GWAS, we were unable to detect loci associated with 18:3 variation (Figure 6A, Table S2). However, multi-GWAS that leveraged unexpected correlations between 18:3 and other fatty acids revealed a locus in Mrg05 (cluster *e,* Figure 6) associated with substantial differences in 18:3 content relative to the more abundant fatty acids (Figure 6B). Markers in cluster *e* may predict variation in 18:3 content associated with postharvest stability in oat breeding programs.

In conclusion, we have demonstrated the power of multivariate models to discover genetic associations of metabolites that share a biosynthetic pathway. By simultaneously accounting for variation in all ten fatty acids, we were able to detect SNPs that were otherwise not detectable in our population. The improved sensitivity of multivariate GWAS may be particularly advantageous for polyploid species like oat, where the effect of a single locus may be buffered by the activity of its homeologs (Santantonio *et al.* 2019). On the other hand, the poorly characterized structural variation within oat haplotypes in combination with long-range LD currently presents a significant challenge for resolving GWAS signals to individual genes. The present study provides a foundation for future investigations of the genetic basis of fatty acid variation in oat. With anticipated improvements in genomic resources and the implementation of LD-independent approaches such as transcriptome-wide association studies (Gusev *et al.* 2016) we expect further resolution of the genetic control of fatty acid composition.

## ACKNOWLEDGMENTS

The authors wish to thank the Proteomics and Metabolomics Facility at Colorado State University for FAME analysis. This research was supported by PepsiCo Global R&D, the USDA– ARS, USDA-NIFA-AFRI 2011-68002-30029 (Triticeae-CAP), USDA-NIFA-AFRI 2017-67007-25939 (Wheat-CAP), USDA-NIFA-AFRI 2017-67007-26502, and Hatch Project 149-447. Disclaimer: The views expressed in this article are those of the authors and do not necessarily reflect the position or policy of PepsiCo, Inc. Mention of a trademark or proprietary product does not constitute a guarantee or warranty of the product by the USDA and does not imply its approval to the exclusion of other products that may also be suitable. The USDA is an equal opportunity provider and employer.

## SUPPLEMENTAL FIGURE LEGENDS

**Figure S1. Representative GC-MS chromatogram of oat FAMEs.** The identity of peaks that were quantified is shown in black. Minor peaks that were frequently below the limit of detection are shown in red. 18:1* designates an unidentified isomer of 18:1.

**Figure S2. Principal component analysis (PCA) of genotype data matrix.** A. The 492 oat lines plotted along the first two principal components (PCs), colored with respect to line origin (AFRICore, Asoro, South America, North America, Europe, and Others). B. Analogous to A, except here the third and fourth PCs are plotted.

**Figure S3. Fatty acid phenotypes measured in two environments.** A. A heatmap of pairwise Spearman’s rank correlations (ρ) between FAME measurements in each of the two environments, referred to as trait-environment combinations. Trait-environment combinations are ordered by hierarchical clustering, as pictured in B. Environments are differentiated by axis label color, with Env 1 in blue and Env 2 in gray. B. Hierarchical clustering of trait-environment combinations with environment indicated by label color as in A. C. A comparison of FAME distributions for measurements in the two environments. A heatmap (D) and dendrogram (E) of log-contrast transformed traits. F. A comparison of FAME proportions for measurements in the two environments.

**Figure S4. Correlations (*r*) and partial correlations (*pr*) between FAME BLUPs.** A. Overlaid histograms of the pairwise *r* and *pr* values between untransformed BLUPs, and heatmaps displaying pairwise *r* (B) and pairwise *pr* (C).

**Figure S5. Principal component analysis (PCA) of FAME BLUPs.** A. Oat lines plotted along the first and second (left) and third and fourth (center) principal components (PCs) from a PCA on the mean-centered and unit-variance scaled FAME best-linear unbiased predictors. Phenotypic variance explained plotted as a function of PC (right). Dots are shaded based on the total FAME BLUP of each line as indicated. B. Loadings of each FAME with respect to PC, with each FAME distinguished by a unique color (see legend). Note, saturated and unsaturated FAMEs are portrayed in warm and cool colors, respectively.

**Figure S6. Manhattan and quantile-quantile (QQ) plots for GWAS of the ten principal components (PCs).** Dotted lines denote the three significance thresholds considered, with a Bonferroni-corrected threshold of 5% in red, and 5 and 10% false-discovery rate (FDR) thresholds in light and dark gray, respectively. If a *p*-value passed the Bonferroni threshold, then the corresponding marker was portrayed in red.

**Figure S7. Manhattan and quantile-quantile (QQ) plots for GWAS of the ten FAME BLUPs.** Dotted lines denote the three significance thresholds considered, with a Bonferroni-corrected threshold of 5% in red, and 5 and 10% false-discovery rate (FDR) thresholds in light and dark gray, respectively. If a *p*-value passed the Bonferroni threshold, then the corresponding marker was portrayed in red.

**Figure S8. Manhattan and quantile-quantile (QQ) plots for GWAS of the ten quantile-transformed FAME BLUPs.** Dotted lines denote the three significance thresholds considered, with a Bonferroni-corrected threshold of 5% in red, and 5 and 10% false-discovery rate (FDR) thresholds in light and dark gray, respectively. If a *p*-value passed the Bonferroni threshold, then the corresponding marker was portrayed in red.

**Figure S9. Mean phenotypic differences between distinct homozygote classes at 152 significantly associated SNPs.** Genotype is plotted on the x-axis, with the mean number of phenotypic standard deviations away from the mean on the y-axis. Each centered and unit-variance scaled FAME BLUP (including total) is plotted, with warm colors for saturated and cool colors for unsaturated FAMEs. Total is indicated by a black line. The bold lines indicate that the pairwise t-test between genotype class means was significant at α=0.05 after correcting for multiple testing with a Bonferroni correction. To simplify the plots, only traits with significantly different means between genotypes are labeled. As lines with missing genotypes are excluded from these calculations, we present the genotype counts in the bottom right or left-hand corner of each plot.

## SUPPLEMENTAL TABLE LEGENDS

**Supplemental Table S1. Line-mean heritabilities (*h*_*l*_ ^2^) for untransformed BLUPs of 11 FAME traits.**

**Supplemental Table S2. Combined GWAS results for all significant SNPs detected at FDR of 10%.** For the traits, qn denotes quantile-normalized BLUPs, nt denotes non-transformed BLUPs. Linkage disequilibrium (LD) Subset, if 1, denotes SNPs that are in LD with at least 2 other significant SNPs across all analyses (see *Materials and Methods*). MAF = minor allele frequency. Δ*R*^2^ *LR* is an approximation of the percent phenotypic variance explained (PVE) for each SNP expressed as the difference between the *R*^2^ *LR* statistics (Sun *et al.* 2010) with or without the SNP included as a covariate in the GWAS model. See Supplemental Table 3 for *R*^2^ *LR* statistics related to the multi-GWAS hits.

**Supplemental Table S3. Decomposition of variance explained by 148 SNPs detected using a ten trait multivariate GWAS model.** Linkage disequilibrium (LD) Subset, if 1, denotes SNPs that are in LD with at least two other significant SNPs across all analyses (see *Materials and Methods*). MAF = minor allele frequency. Δ*R*^2^ *LR* is an approximation of the percent phenotypic variance explained (PVE) for each SNP expressed as the difference between the *R*^2^ *LR* statistics (Sun *et al.* 2010) with or without the SNP included as a covariate in each trait’s univariate model.

