## Supplementary figures and images for "Multivariate Genome-wide Association Analyses Reveal the Genetic Basis of Seed Fatty Acid Composition in Oat (*Avena sativa* L.)"

### Supplemental Figure S1

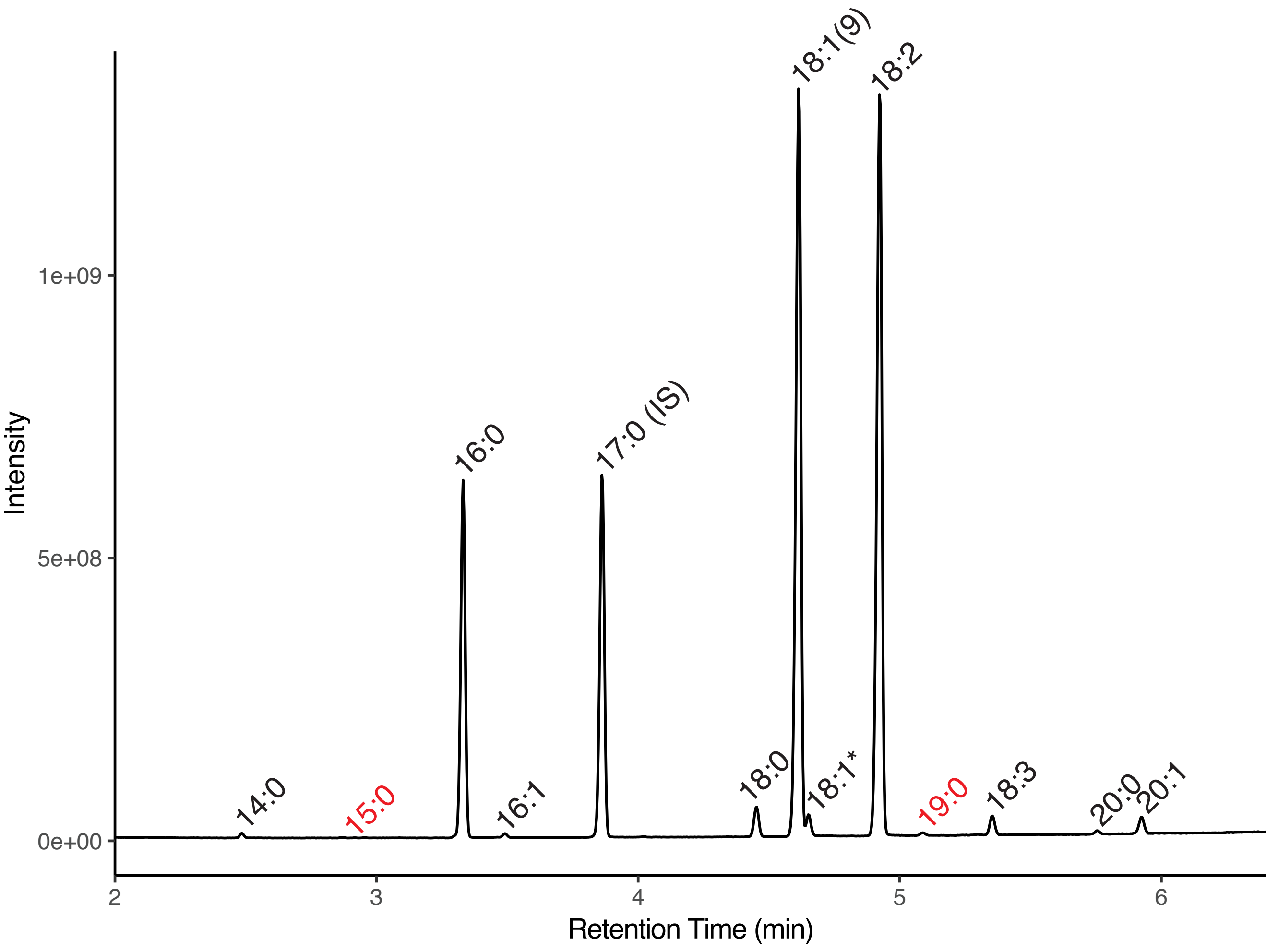

### Supplemental Figure S2

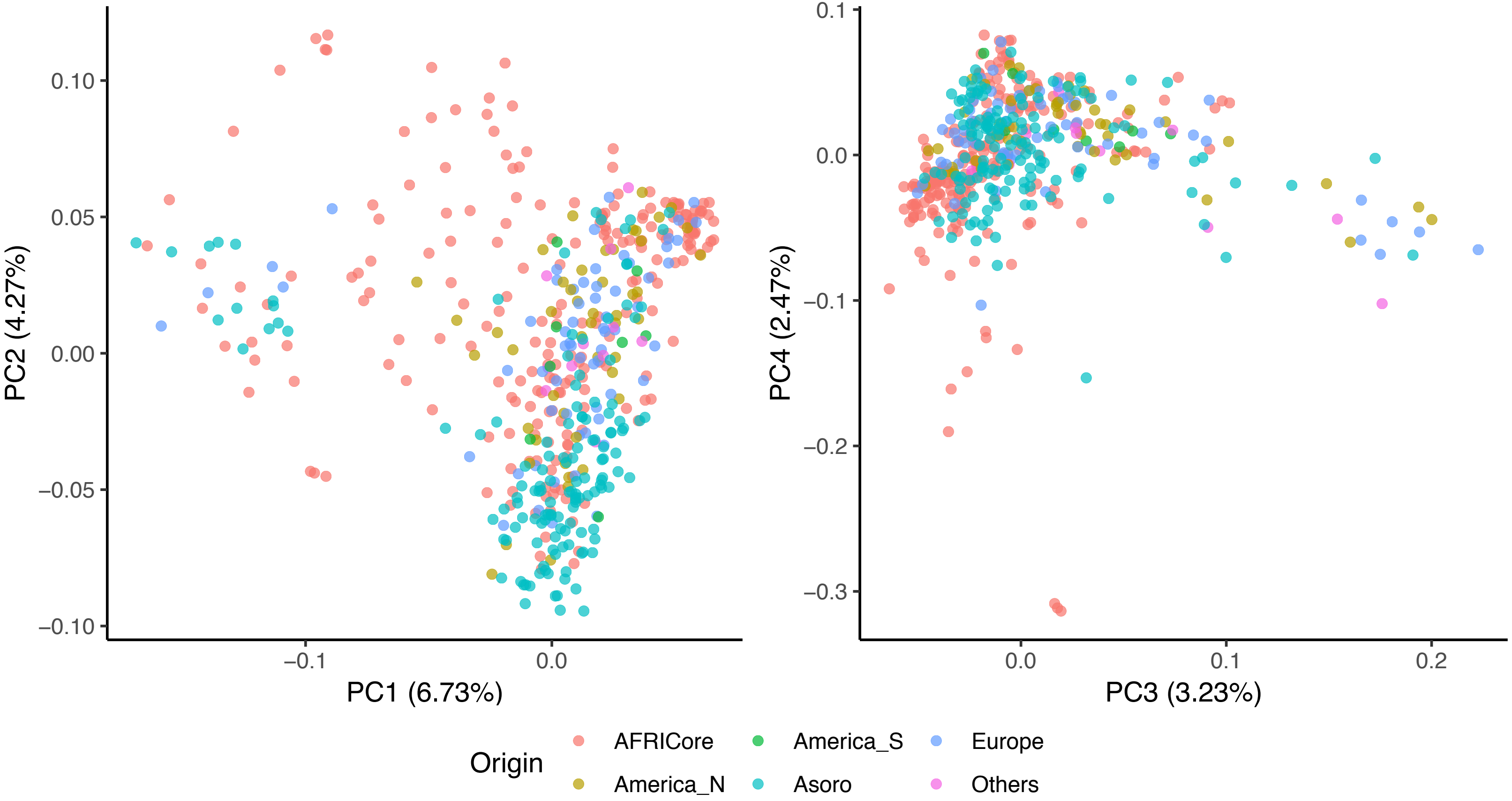

### Supplemental Figure S3

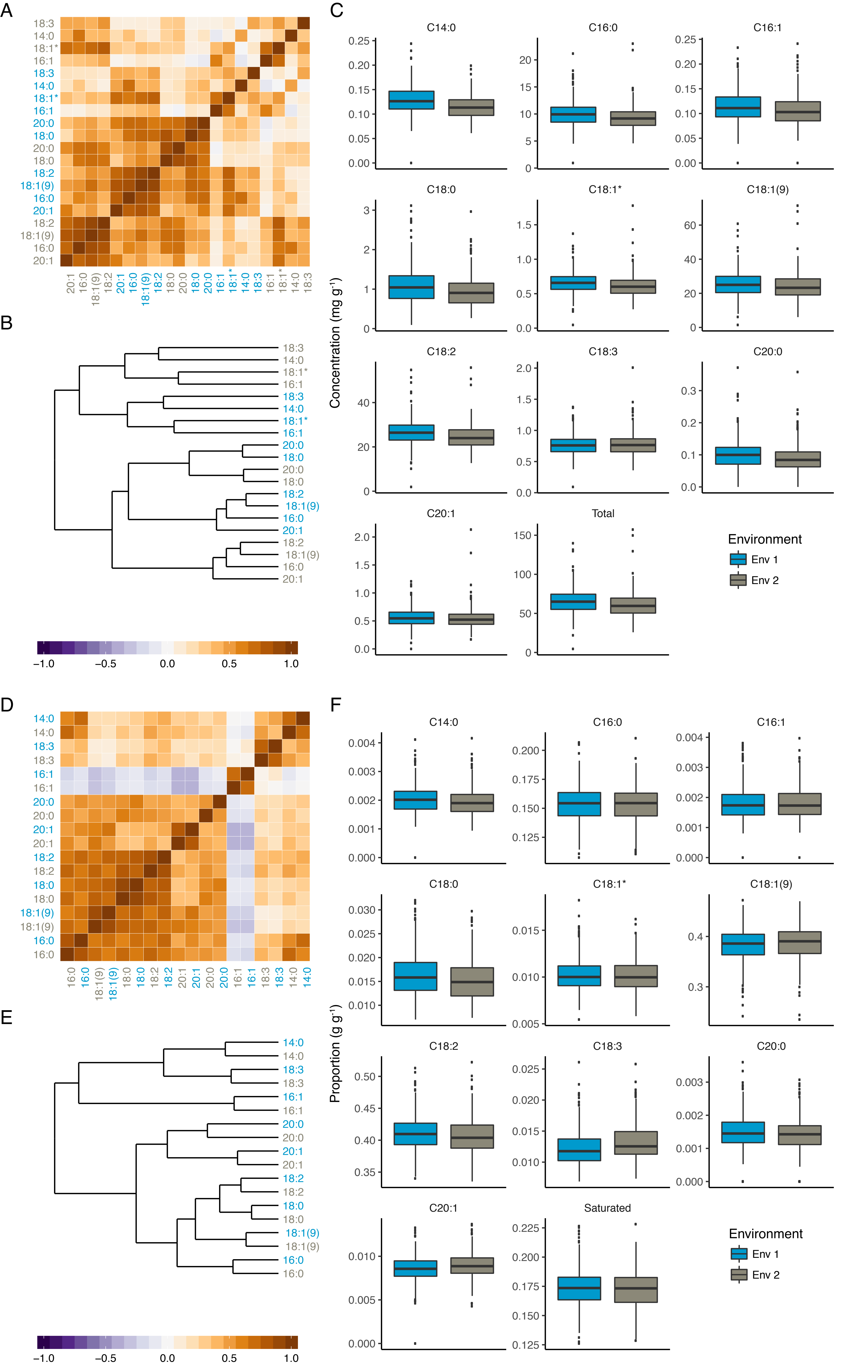

### Supplemental Figure S4

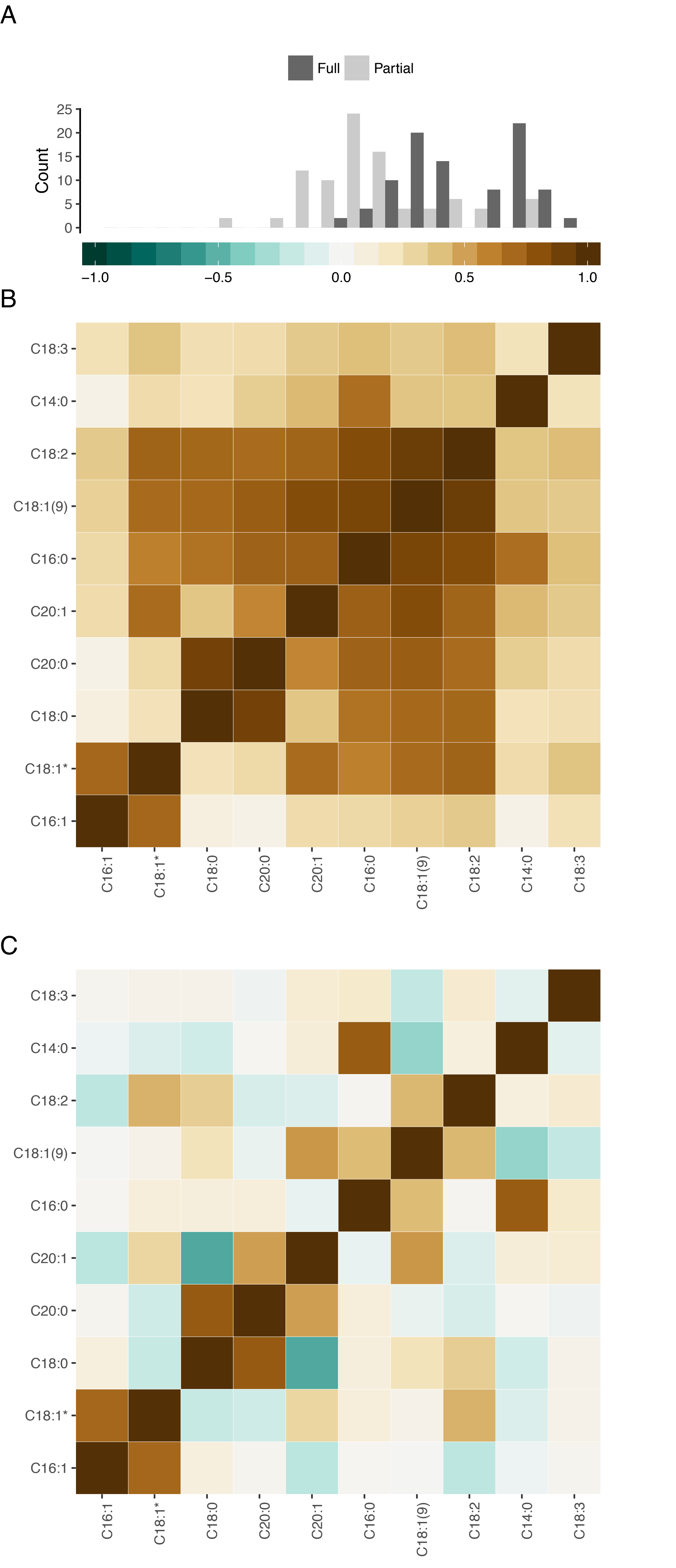

### Supplemental Figure S5

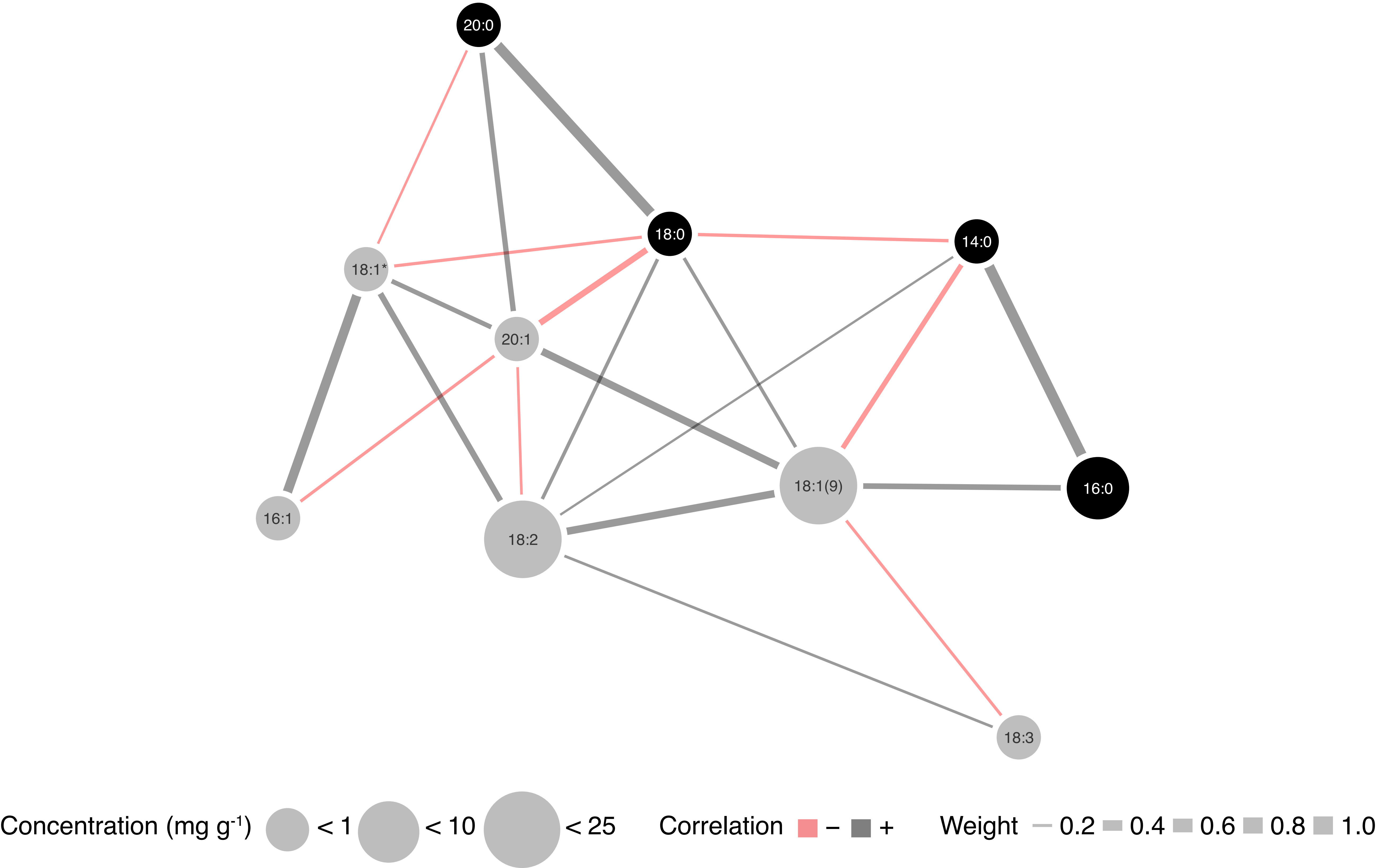

### Supplemental Figure S6

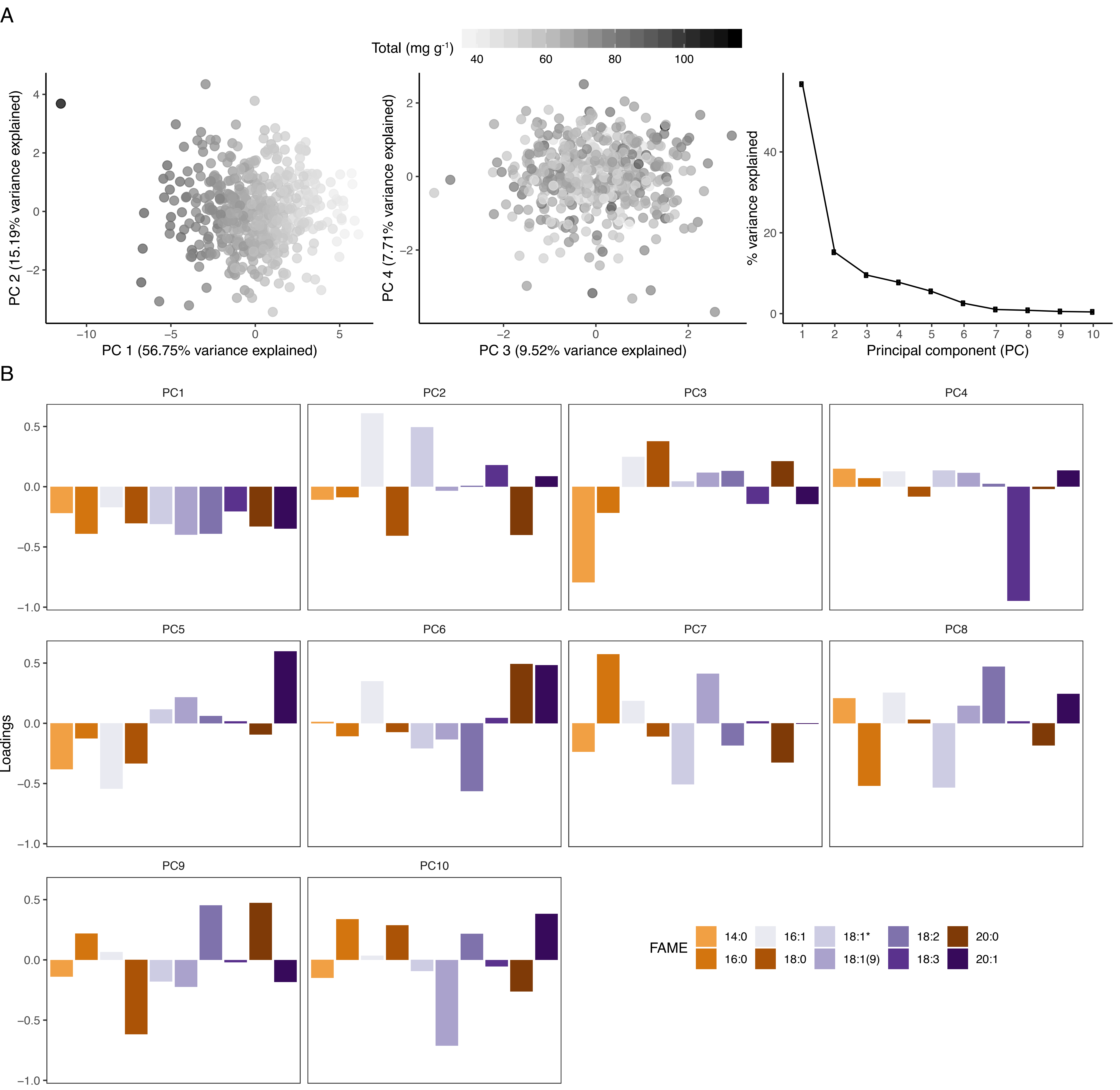

### Supplemental Figure S7

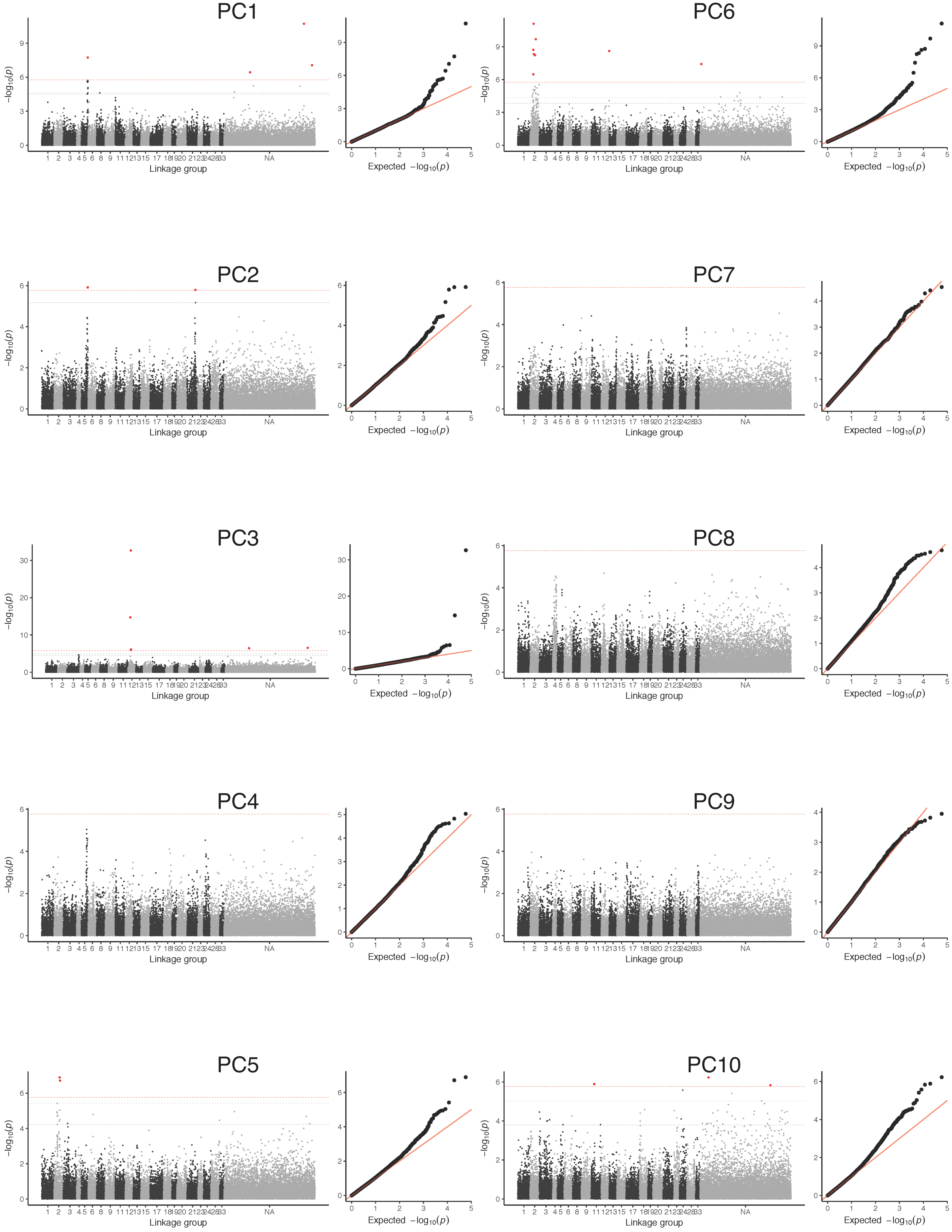

### Supplemental Figure S8

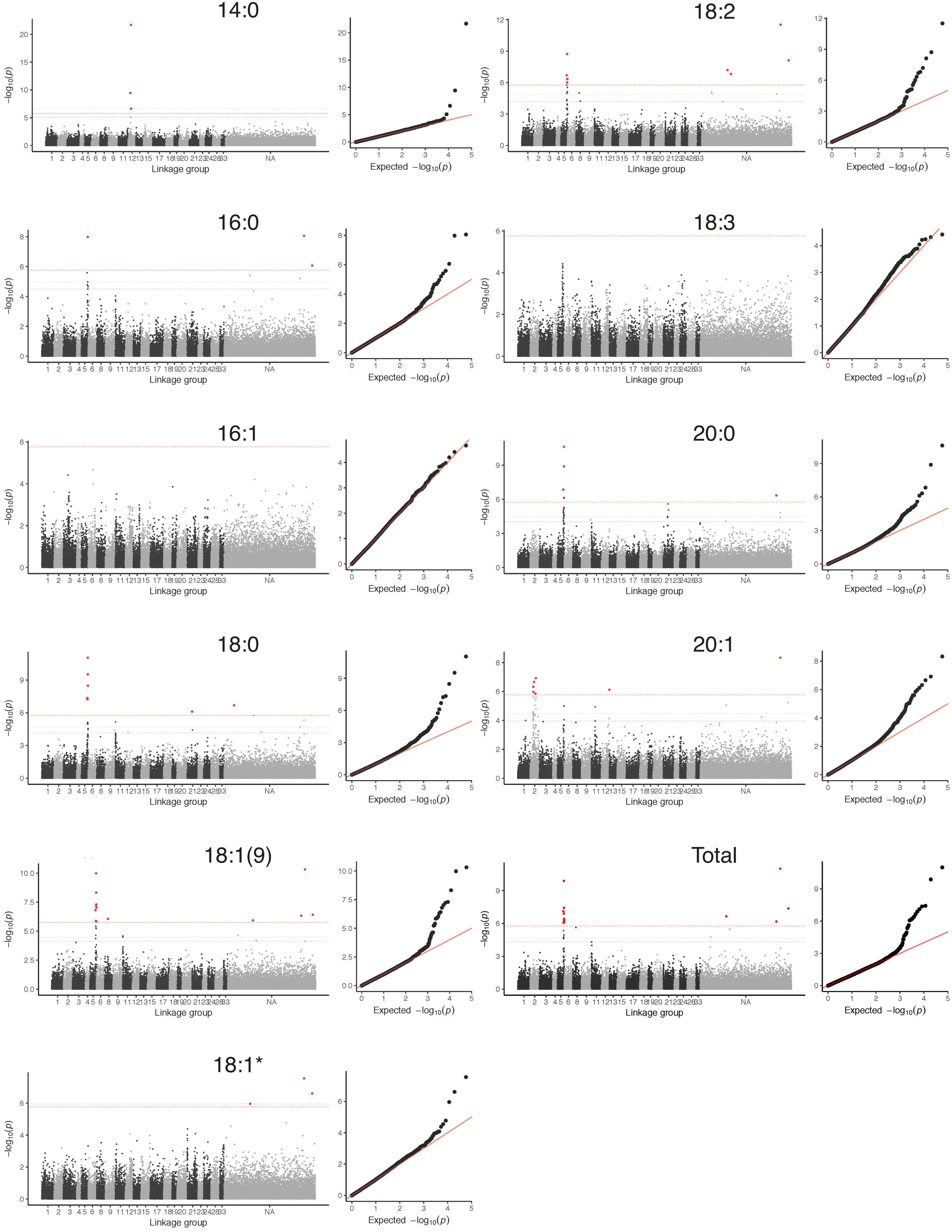

### Supplemental Figure S9

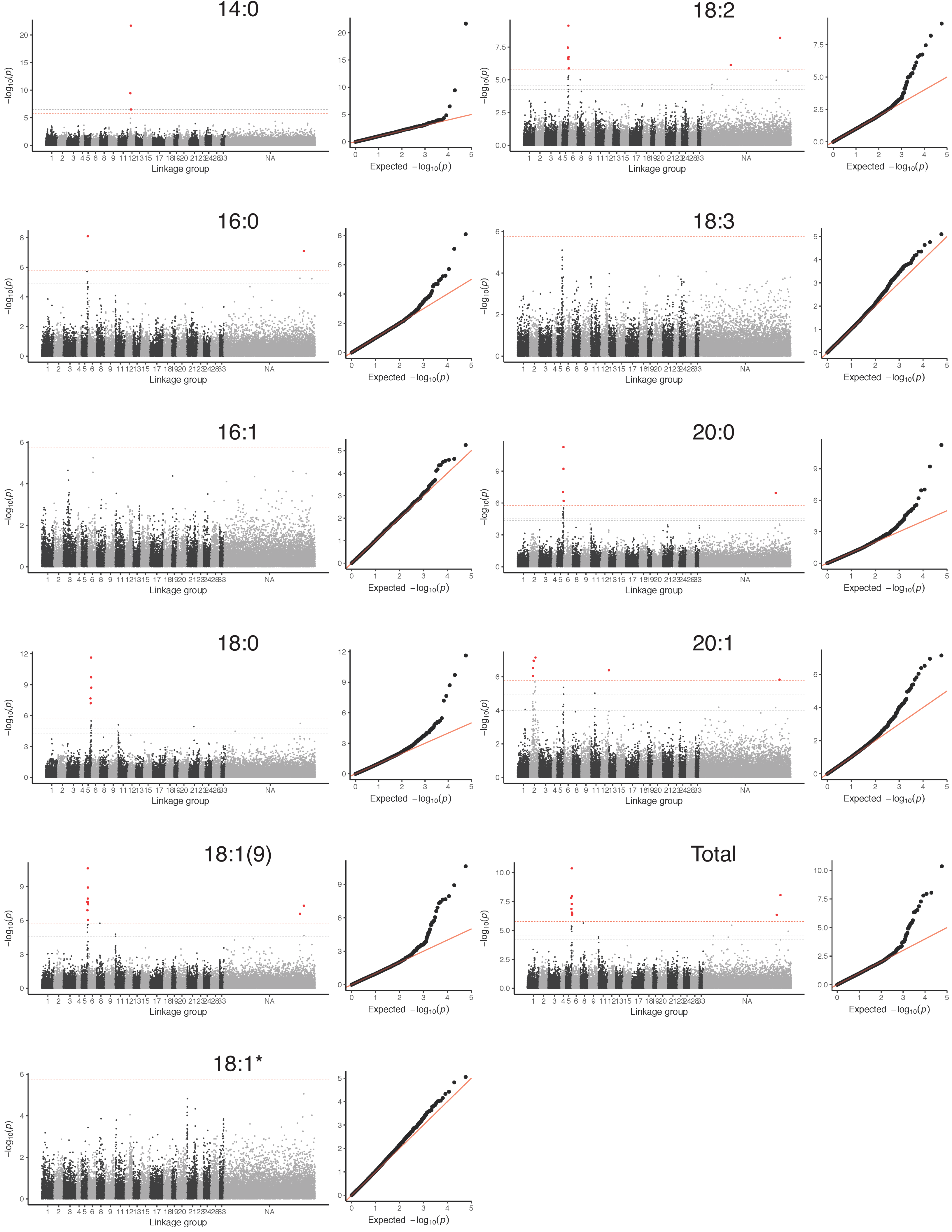
