## Supplemental Figure S10 for "Multivariate Genome-wide Association Analyses Reveal the Genetic Basis of Seed Fatty Acid Composition in Oat (*Avena sativa* L.)"

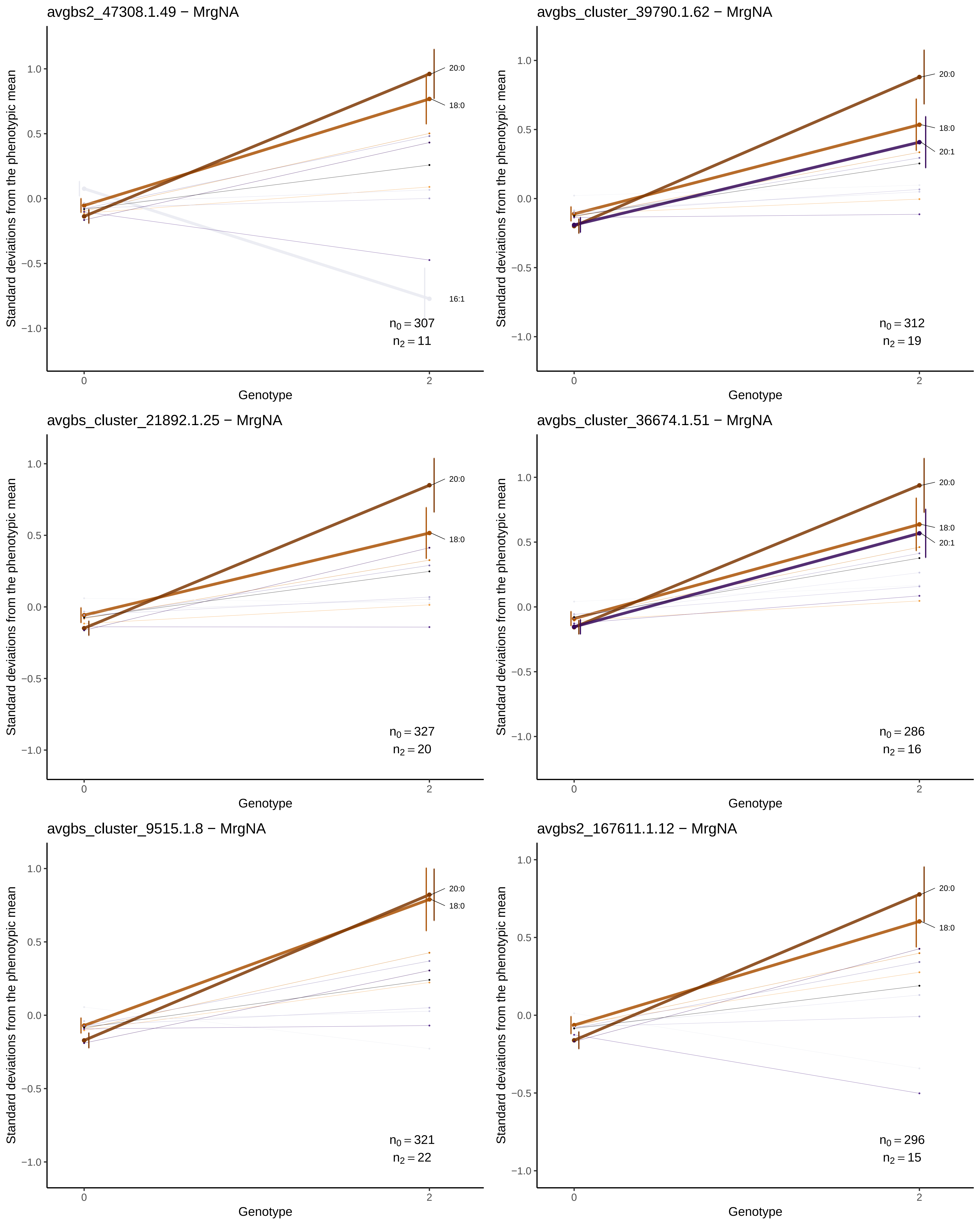
avgbs2_47308.1.49 − MrgNA
Standard deviations from the phenotypic mean
Genotype
1.0
0.5
0.0
−0.5
−1.0
0
2
20:0
18:0
16:1
$n_0 = 307$
$n_2 = 11$
avgbs_cluster_39790.1.62 − MrgNA
Standard deviations from the phenotypic mean
Genotype
1.0
0.5
0.0
−0.5
−1.0
0
2
20:0
18:0
20:1
$n_0 = 312$
$n_2 = 19$
avgbs_cluster_21892.1.25 − MrgNA
Standard deviations from the phenotypic mean
Genotype
1.0
0.5
0.0
−0.5
−1.0
0
2
20:0
18:0
$n_0 = 327$
$n_2 = 20$
avgbs_cluster_36674.1.51 − MrgNA
Standard deviations from the phenotypic mean
Genotype
1.0
0.5
0.0
−0.5
−1.0
0
2
20:0
18:0
20:1
$n_0 = 286$
$n_2 = 16$
avgbs_cluster_9515.1.8 − MrgNA
Standard deviations from the phenotypic mean
Genotype
1.0
0.5
0.0
−0.5
−1.0
0
2
20:0
18:0
$n_0 = 321$
$n_2 = 22$
avgbs2_167611.1.12 − MrgNA
Standard deviations from the phenotypic mean
Genotype
1.0
0.5
0.0
−0.5
−1.0
0
2
20:0
18:0
$n_0 = 296$
$n_2 = 15$

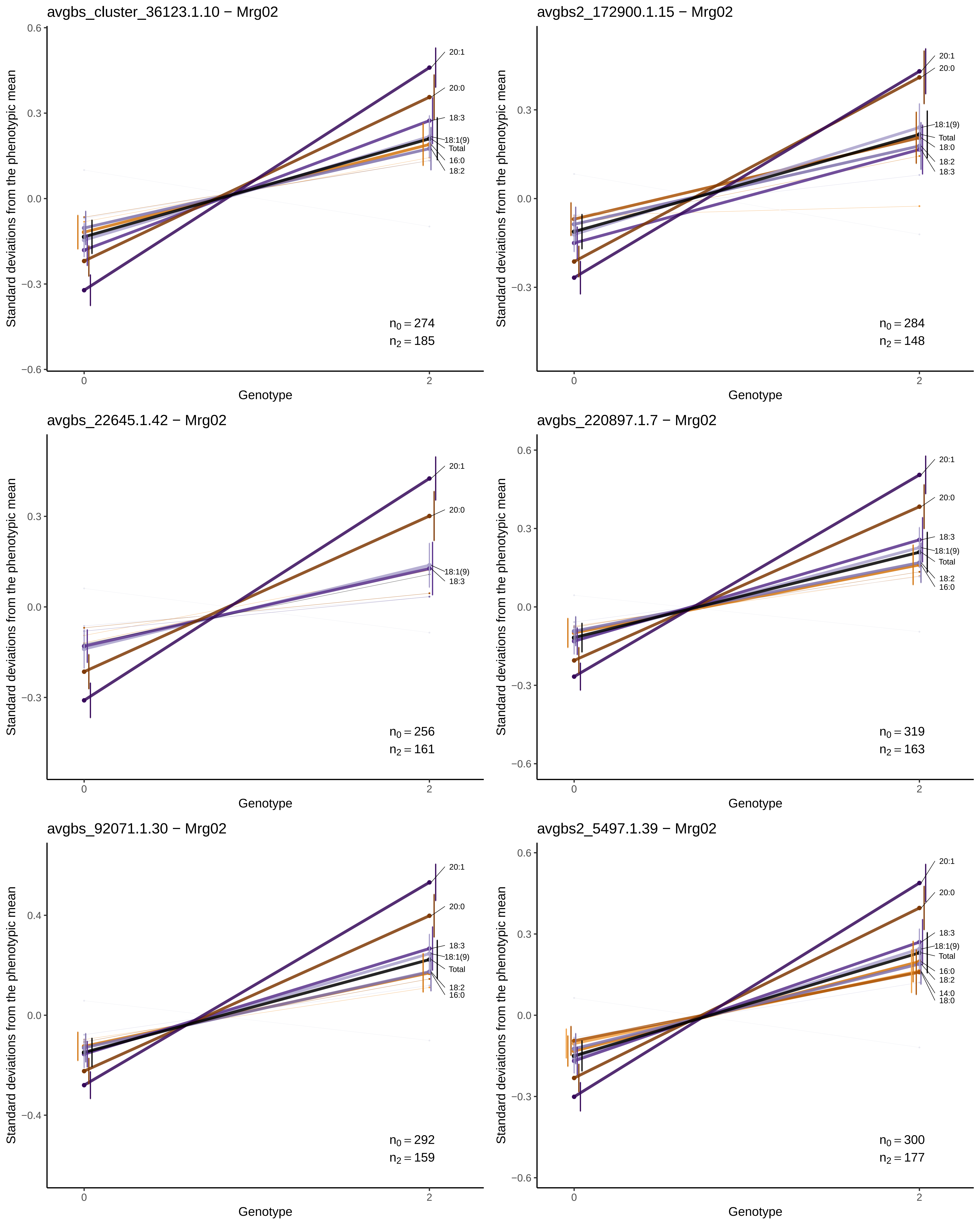

avgbs_cluster_36123.1.10 − Mrg02
Standard deviations from the phenotypic mean
Genotype
20:1
20:0
18:3
18:1(9)
Total
16:0
18:2
$n_0 = 274$
$n_2 = 185$
avgbs2_172900.1.15 − Mrg02
Standard deviations from the phenotypic mean
Genotype
20:1
20:0
18:1(9)
Total
18:0
18:2
18:3
$n_0 = 284$
$n_2 = 148$
avgbs_22645.1.42 − Mrg02
Standard deviations from the phenotypic mean
Genotype
20:1
20:0
18:1(9)
18:3
$n_0 = 256$
$n_2 = 161$
avgbs_220897.1.7 − Mrg02
Standard deviations from the phenotypic mean
Genotype
20:1
20:0
18:3
18:1(9)
Total
18:2
16:0
$n_0 = 319$
$n_2 = 163$
avgbs_92071.1.30 − Mrg02
Standard deviations from the phenotypic mean
Genotype
20:1
20:0
18:3
18:1(9)
Total
18:2
16:0
$n_0 = 292$
$n_2 = 159$
avgbs2_5497.1.39 − Mrg02
Standard deviations from the phenotypic mean
Genotype
20:1
20:0
18:3
18:1(9)
Total
16:0
18:2
14:0
18:0
$n_0 = 300$
$n_2 = 177$

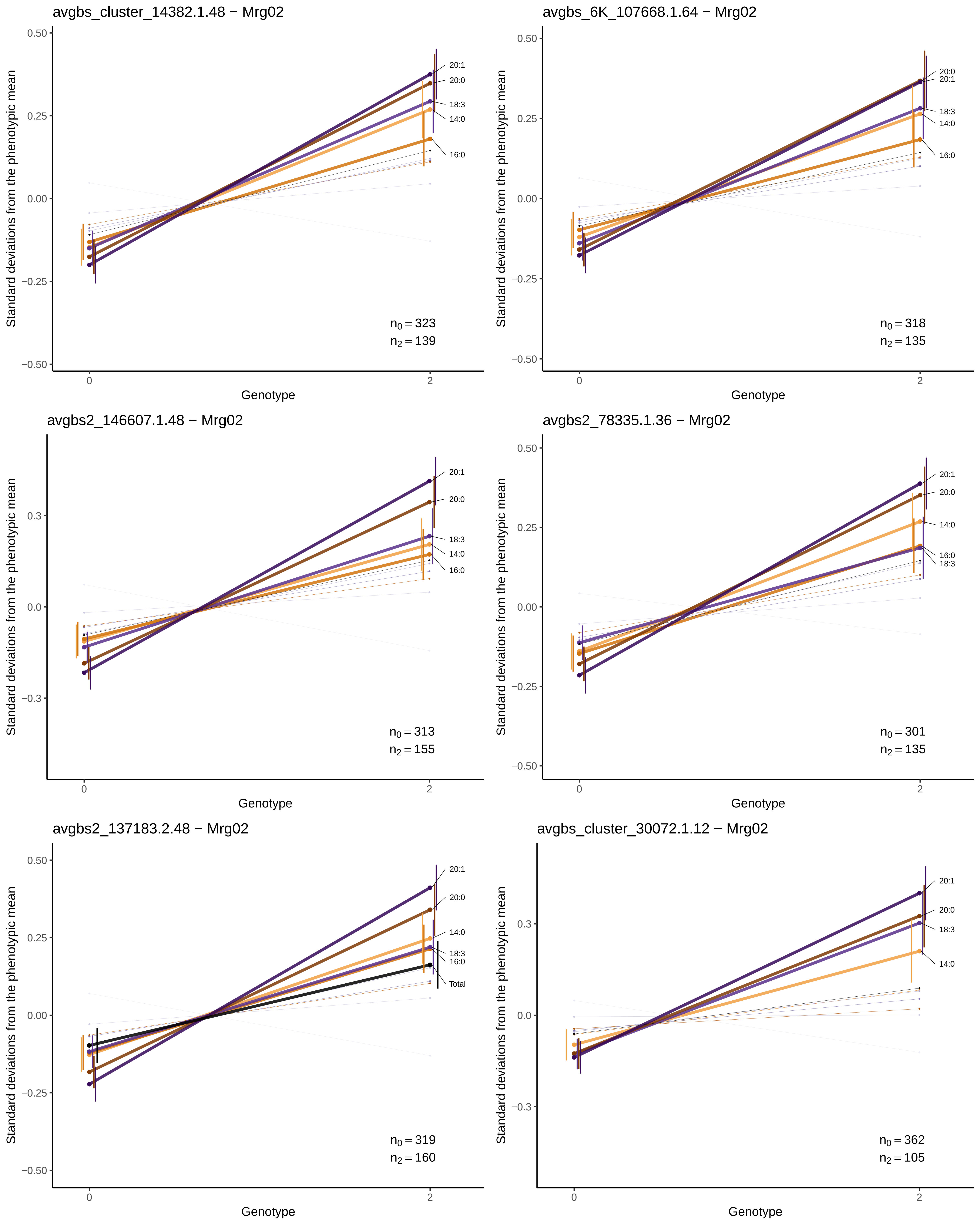
avgbs_cluster_14382.1.48 − Mrg02
Standard deviations from the phenotypic mean
0.50
0.25
0.00
−0.25
−0.50
20:1
20:0
18:3
14:0
16:0
$n_0 = 323$
$n_2 = 139$
0
2
Genotype
avgbs_6K_107668.1.64 − Mrg02
Standard deviations from the phenotypic mean
0.50
0.25
0.00
−0.25
−0.50
20:0
20:1
18:3
14:0
16:0
$n_0 = 318$
$n_2 = 135$
0
2
Genotype
avgbs2_146607.1.48 − Mrg02
Standard deviations from the phenotypic mean
0.3
0.0
-0.3
20:1
20:0
18:3
14:0
16:0
$n_0 = 313$
$n_2 = 155$
0
2
Genotype
avgbs2_78335.1.36 − Mrg02
Standard deviations from the phenotypic mean
0.50
0.25
0.00
−0.25
−0.50
20:1
20:0
14:0
16:0
18:3
$n_0 = 301$
$n_2 = 135$
0
2
Genotype
avgbs2_137183.2.48 − Mrg02
Standard deviations from the phenotypic mean
0.50
0.25
0.00
−0.25
−0.50
20:1
20:0
14:0
18:3
16:0
Total
$n_0 = 319$
$n_2 = 160$
0
2
Genotype
avgbs_cluster_30072.1.12 − Mrg02
Standard deviations from the phenotypic mean
0.3
0.0
-0.3
20:1
20:0
18:3
14:0
$n_0 = 362$
$n_2 = 105$
0
2
Genotype

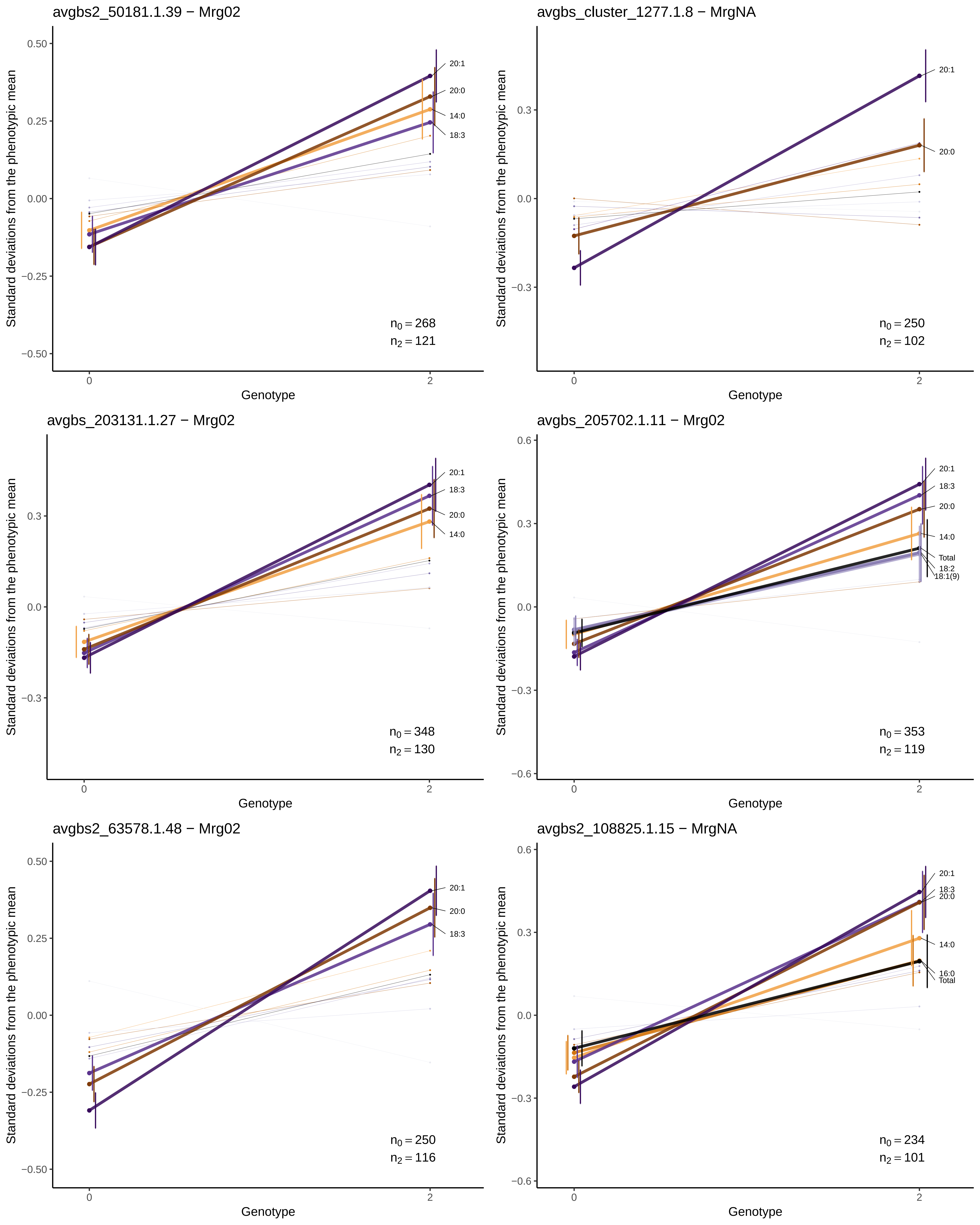
avgbs2_50181.1.39 − Mrg02
Standard deviations from the phenotypic mean
0.50
0.25
0.00
−0.25
−0.50
20:1
20:0
14:0
18:3
$n_0 = 268$
$n_2 = 121$
0
2
Genotype
avgbs_cluster_1277.1.8 − MrgNA
Standard deviations from the phenotypic mean
0.3
0.0
−0.3
20:1
20:0
$n_0 = 250$
$n_2 = 102$
0
2
Genotype
avgbs_203131.1.27 − Mrg02
Standard deviations from the phenotypic mean
0.3
0.0
−0.3
20:1
18:3
20:0
14:0
$n_0 = 348$
$n_2 = 130$
0
2
Genotype
avgbs_205702.1.11 − Mrg02
Standard deviations from the phenotypic mean
0.6
0.3
0.0
−0.3
−0.6
20:1
18:3
20:0
14:0
Total
18:2
18:1(9)
$n_0 = 353$
$n_2 = 119$
0
2
Genotype
avgbs2_63578.1.48 − Mrg02
Standard deviations from the phenotypic mean
0.50
0.25
0.00
−0.25
−0.50
20:1
20:0
18:3
$n_0 = 250$
$n_2 = 116$
0
2
Genotype
avgbs2_108825.1.15 − MrgNA
Standard deviations from the phenotypic mean
0.6
0.3
0.0
−0.3
−0.6
20:1
18:3
20:0
14:0
16:0
Total
$n_0 = 234$
$n_2 = 101$
0
2
Genotype

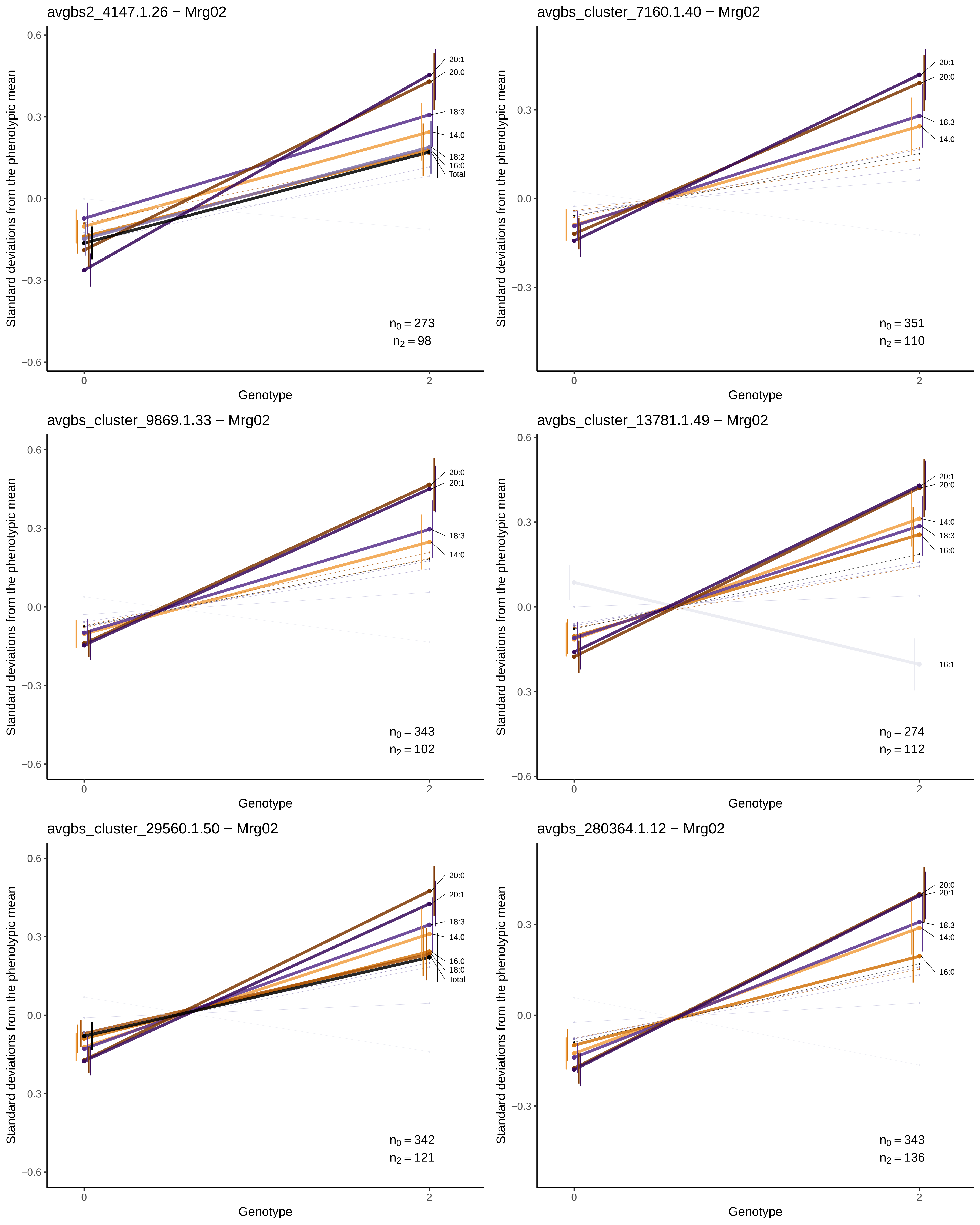
avgbs2_4147.1.26 − Mrg02
Standard deviations from the phenotypic mean
0.6
0.3
0.0
−0.3
−0.6
20:1
20:0
18:3
14:0
18:2
16:0
Total
$n_0 = 273$
$n_2 = 98$
0
2
Genotype
avgbs_cluster_7160.1.40 − Mrg02
Standard deviations from the phenotypic mean
0.3
0.0
−0.3
20:1
20:0
18:3
14:0
$n_0 = 351$
$n_2 = 110$
0
2
Genotype
avgbs_cluster_9869.1.33 − Mrg02
Standard deviations from the phenotypic mean
0.6
0.3
0.0
−0.3
−0.6
20:0
20:1
18:3
14:0
$n_0 = 343$
$n_2 = 102$
0
2
Genotype
avgbs_cluster_13781.1.49 − Mrg02
Standard deviations from the phenotypic mean
0.6
0.3
0.0
−0.3
−0.6
20:1
20:0
14:0
18:3
16:0
16:1
$n_0 = 274$
$n_2 = 112$
0
2
Genotype
avgbs_cluster_29560.1.50 − Mrg02
Standard deviations from the phenotypic mean
0.6
0.3
0.0
−0.3
−0.6
20:0
20:1
18:3
14:0
16:0
18:0
Total
$n_0 = 342$
$n_2 = 121$
0
2
Genotype
avgbs_280364.1.12 − Mrg02
Standard deviations from the phenotypic mean
0.3
0.0
−0.3
20:0
20:1
18:3
14:0
16:0
$n_0 = 343$
$n_2 = 136$
0
2
Genotype

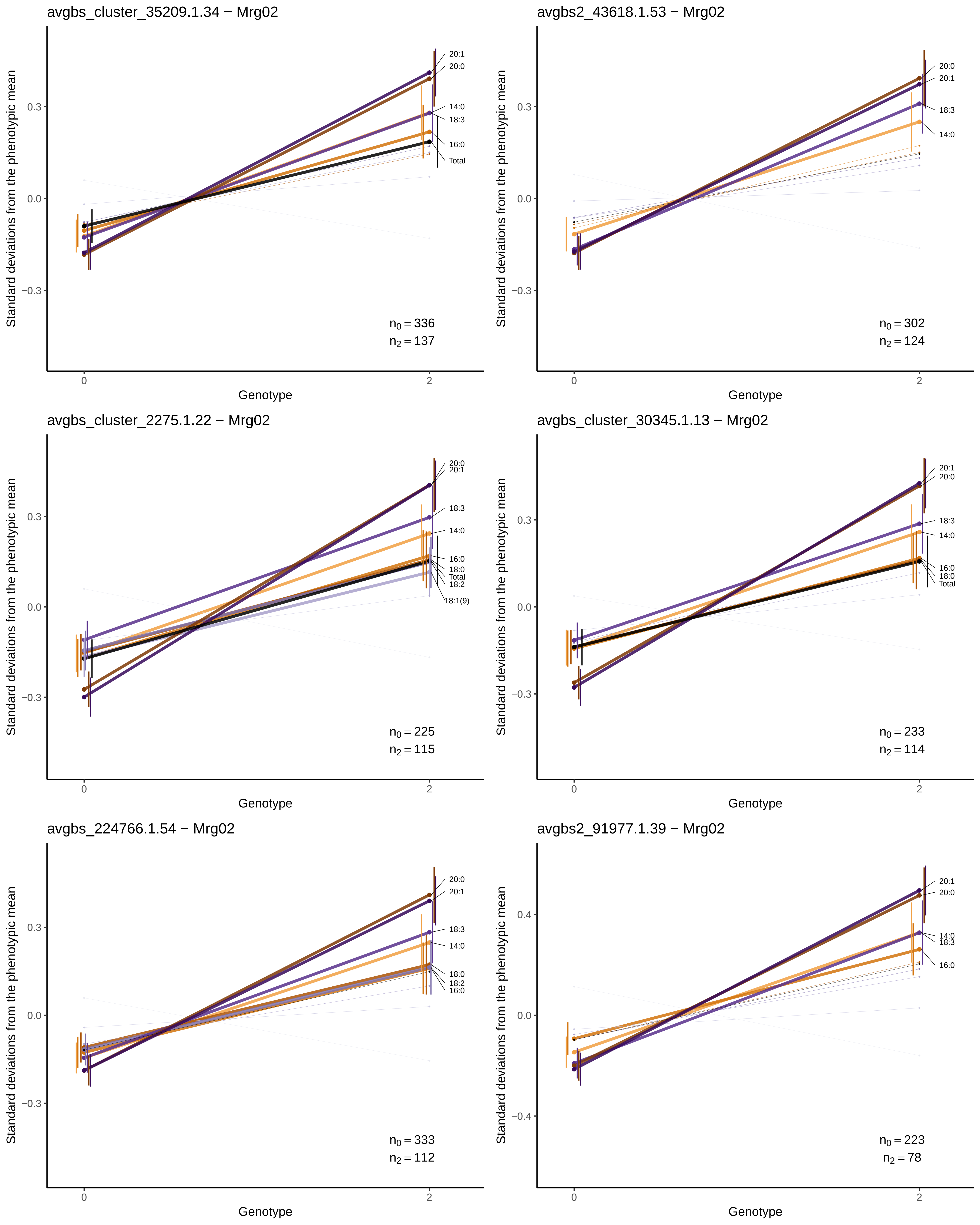
avgbs_cluster_35209.1.34 − Mrg02
Standard deviations from the phenotypic mean
Genotype
20:1
20:0
14:0
18:3
16:0
Total
$n_0 = 336$
$n_2 = 137$
avgbs2_43618.1.53 − Mrg02
Standard deviations from the phenotypic mean
Genotype
20:0
20:1
18:3
14:0
$n_0 = 302$
$n_2 = 124$
avgbs_cluster_2275.1.22 − Mrg02
Standard deviations from the phenotypic mean
Genotype
20:0
20:1
18:3
14:0
16:0
18:0
Total
18:2
18:1(9)
$n_0 = 225$
$n_2 = 115$
avgbs_cluster_30345.1.13 − Mrg02
Standard deviations from the phenotypic mean
Genotype
20:1
20:0
18:3
14:0
16:0
18:0
Total
$n_0 = 233$
$n_2 = 114$
avgbs_224766.1.54 − Mrg02
Standard deviations from the phenotypic mean
Genotype
20:0
20:1
18:3
14:0
18:0
18:2
16:0
$n_0 = 333$
$n_2 = 112$
avgbs2_91977.1.39 − Mrg02
Standard deviations from the phenotypic mean
Genotype
20:1
20:0
14:0
18:3
16:0
$n_0 = 223$
$n_2 = 78$

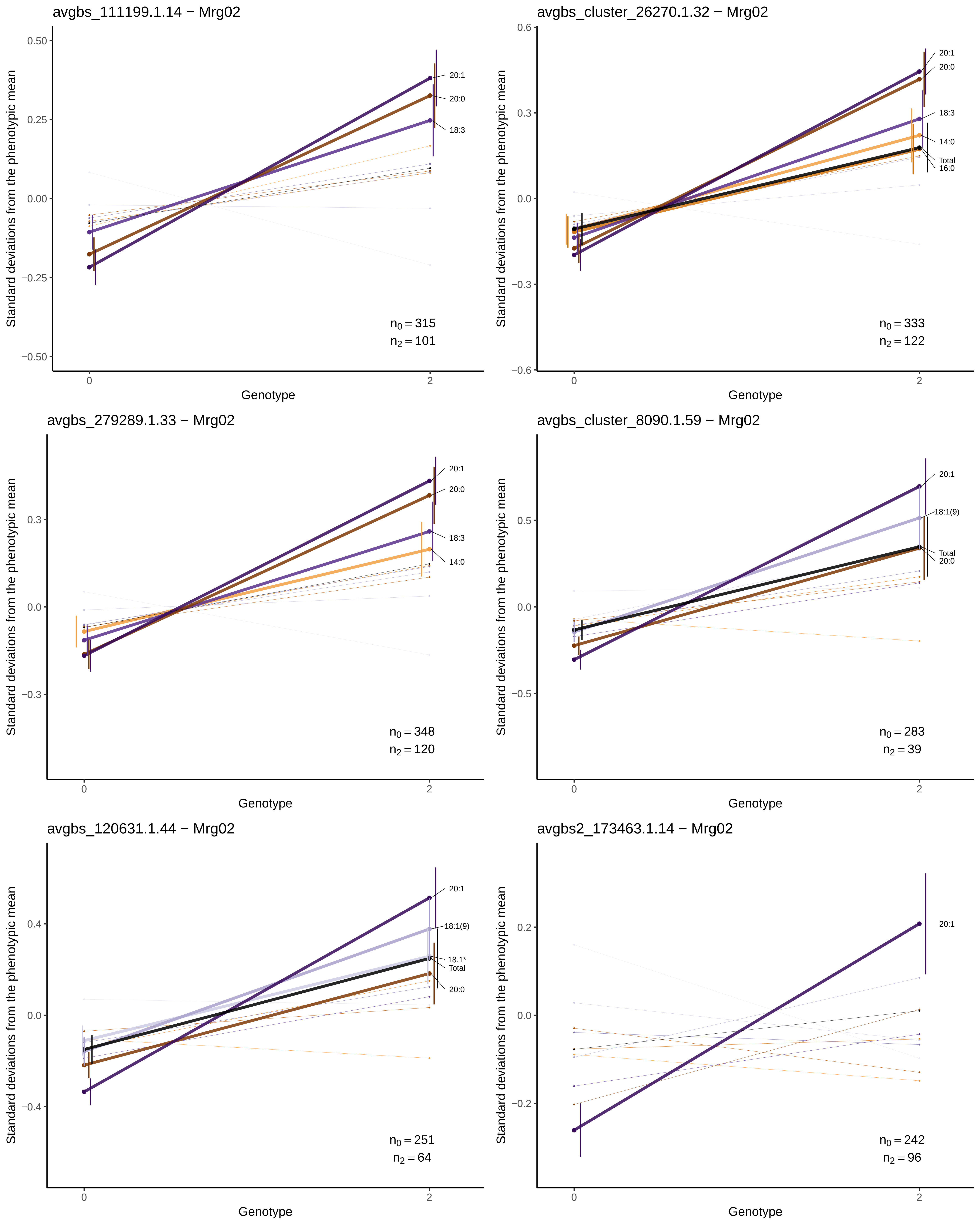
avgbs_111199.1.14 − Mrg02
Standard deviations from the phenotypic mean
Genotype
20:1
20:0
18:3
$n_0 = 315$
$n_2 = 101$
avgbs_cluster_26270.1.32 − Mrg02
20:1
20:0
18:3
14:0
Total
16:0
$n_0 = 333$
$n_2 = 122$
avgbs_279289.1.33 − Mrg02
20:1
20:0
18:3
14:0
$n_0 = 348$
$n_2 = 120$
avgbs_cluster_8090.1.59 − Mrg02
20:1
18:1(9)
Total
20:0
$n_0 = 283$
$n_2 = 39$
avgbs_120631.1.44 − Mrg02
20:1
18:1(9)
18:1*
Total
20:0
$n_0 = 251$
$n_2 = 64$
avgbs2_173463.1.14 − Mrg02
20:1
$n_0 = 242$
$n_2 = 96$

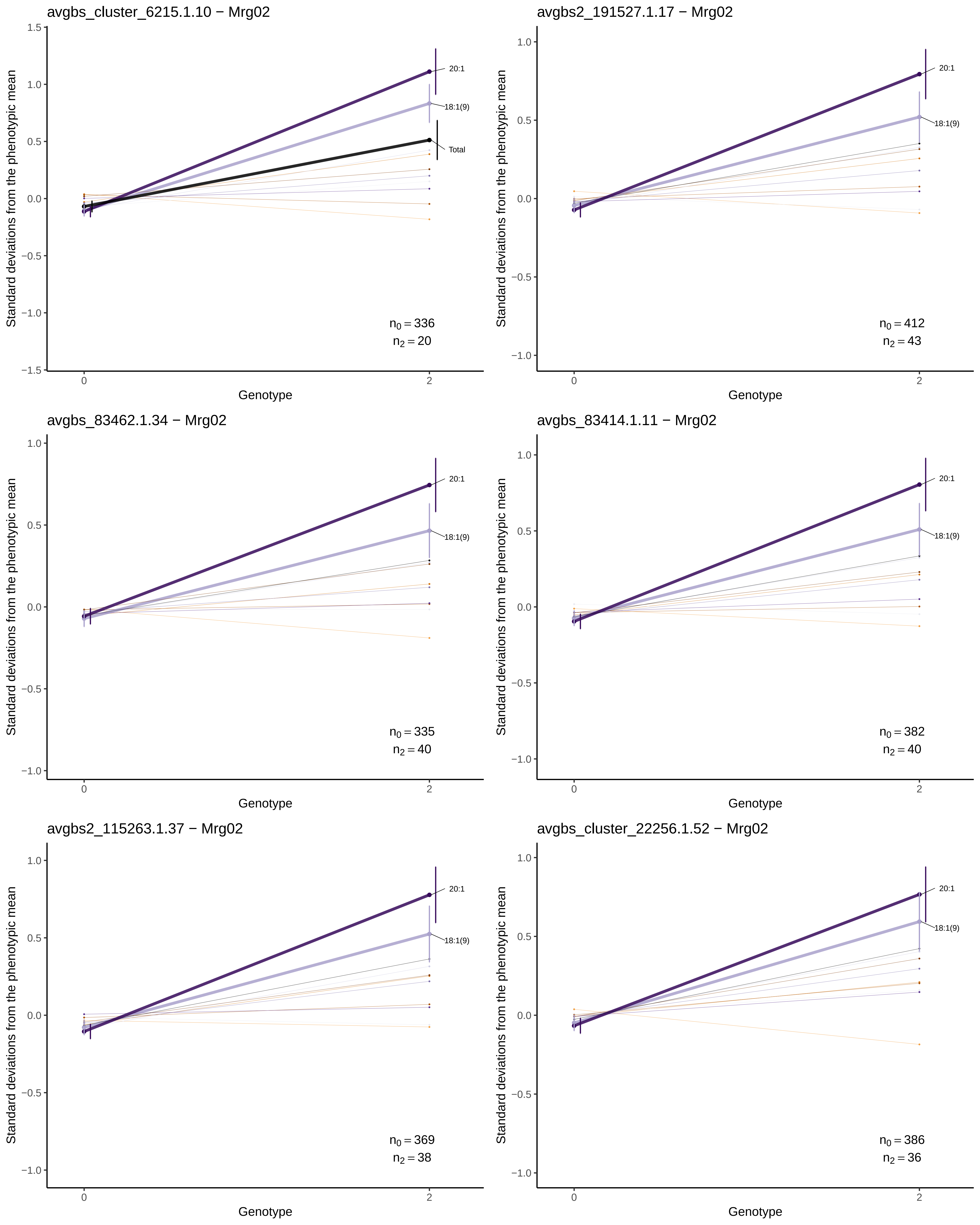
avgbs_cluster_6215.1.10 − Mrg02
20:1
18:1(9)
Total
n0 = 336
n2 = 20
avgbs2_191527.1.17 − Mrg02
20:1
18:1(9)
n0 = 412
n2 = 43
avgbs_83462.1.34 − Mrg02
20:1
18:1(9)
n0 = 335
n2 = 40
avgbs_83414.1.11 − Mrg02
20:1
18:1(9)
n0 = 382
n2 = 40
avgbs2_115263.1.37 − Mrg02
20:1
18:1(9)
n0 = 369
n2 = 38
avgbs_cluster_22256.1.52 − Mrg02
20:1
18:1(9)
n0 = 386
n2 = 36
Standard deviations from the phenotypic mean
Genotype
0
2

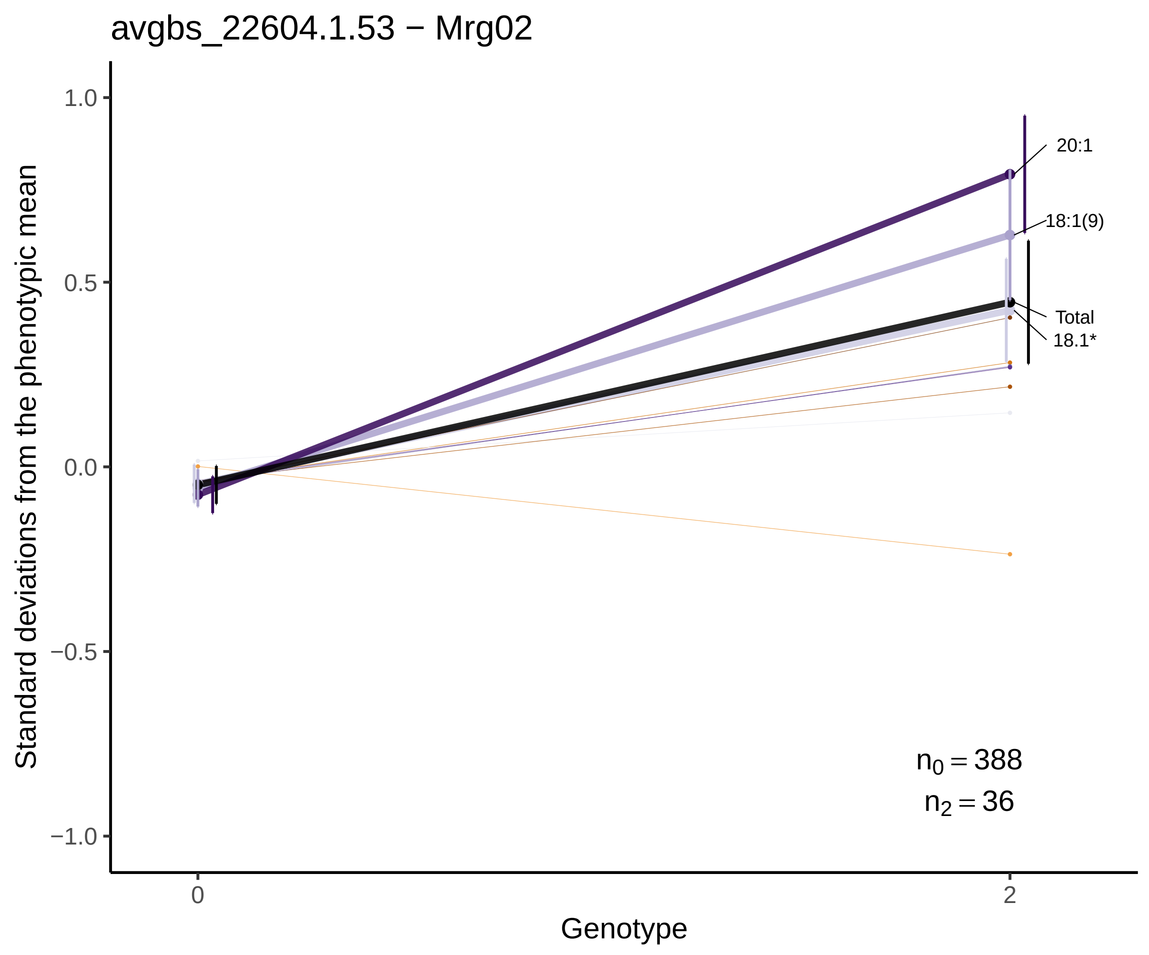
avgbs_22604.1.53 − Mrg02
Standard deviations from the phenotypic mean
1.0
0.5
0.0
−0.5
−1.0
20:1
18:1(9)
Total
18.1*
$n_0 = 388$
$n_2 = 36$
0
2
Genotype

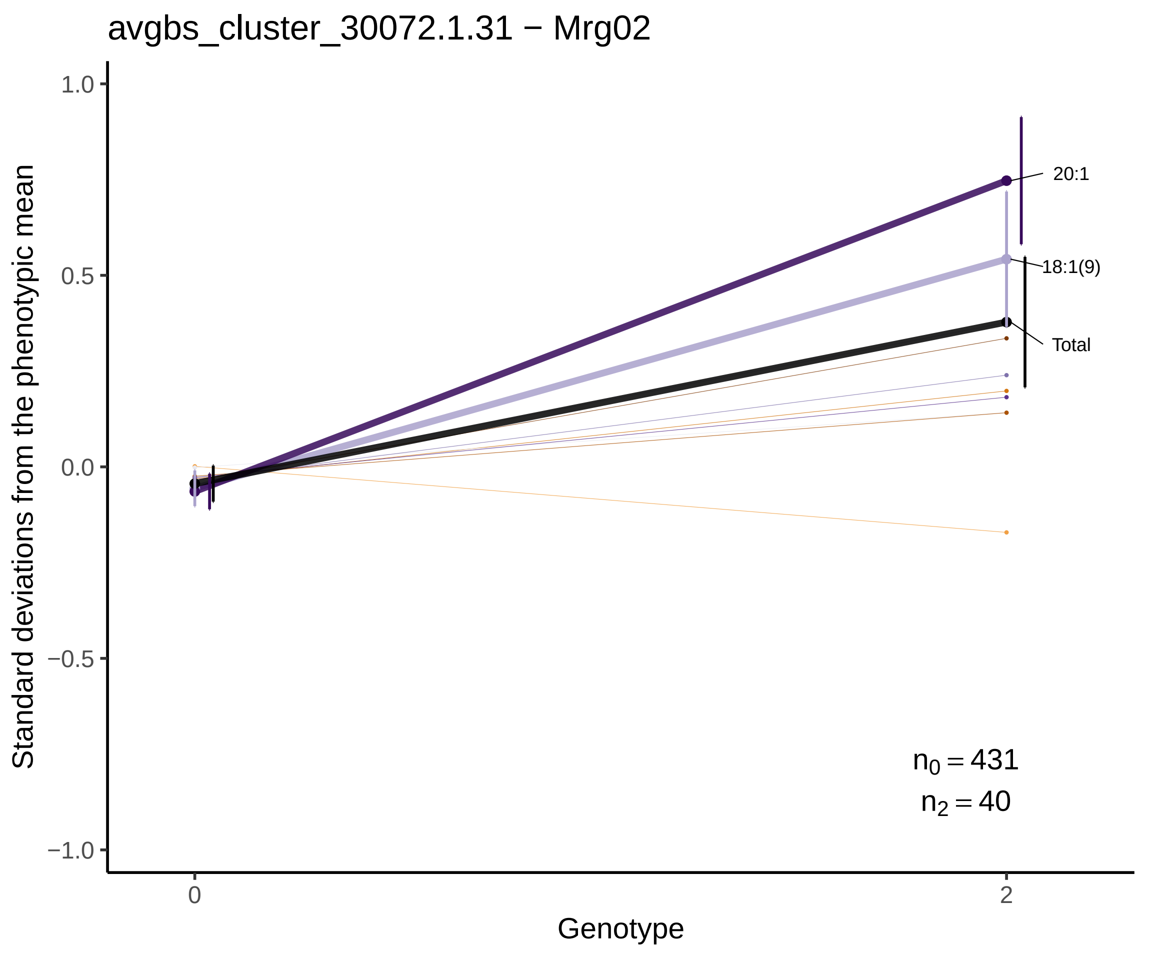
avgbs_cluster_30072.1.31 − Mrg02
Standard deviations from the phenotypic mean
1.0
0.5
0.0
−0.5
−1.0
20:1
18:1(9)
Total
$n_0 = 431$
$n_2 = 40$
0
2
Genotype

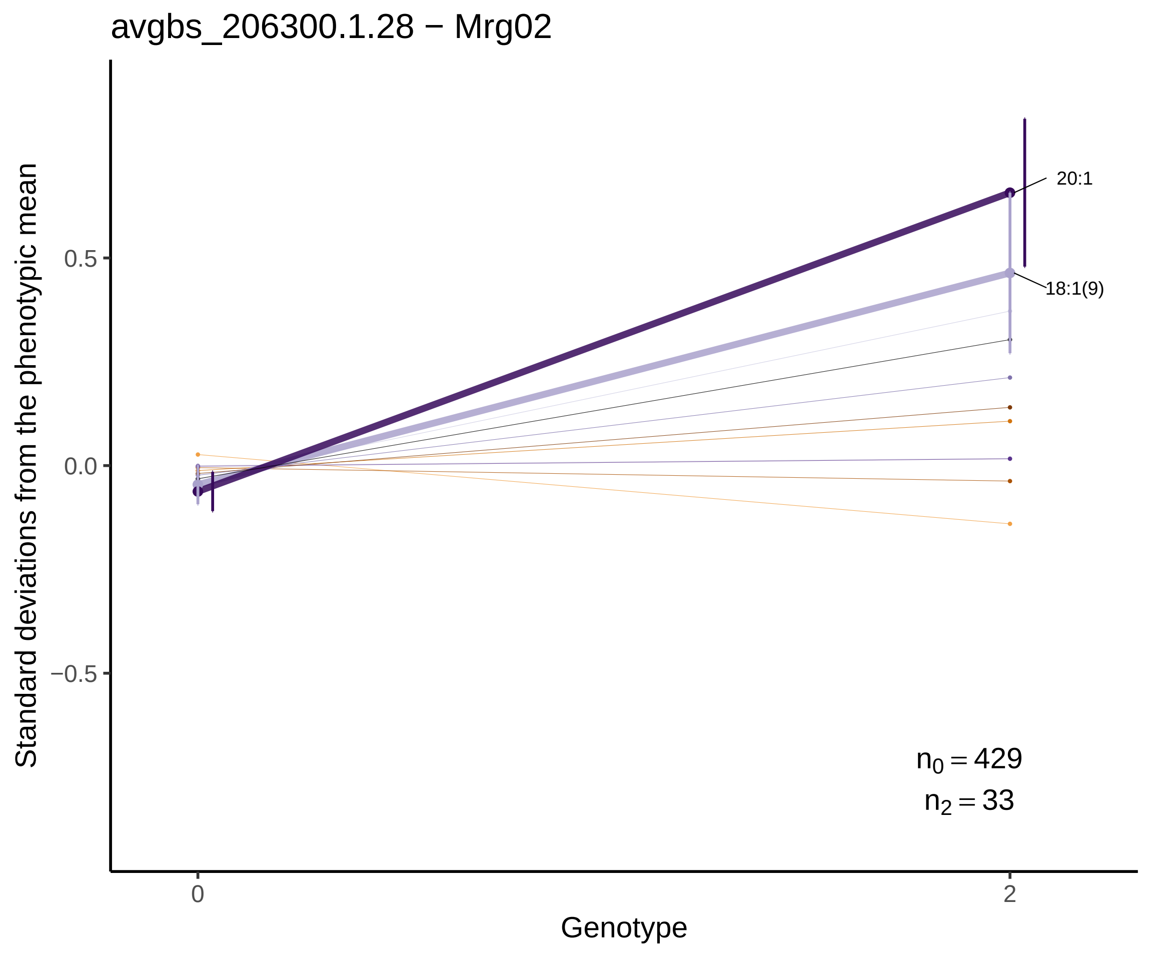
avgbs_206300.1.28 − Mrg02
Standard deviations from the phenotypic mean
0.5
0.0
−0.5
20:1
18:1(9)
$n_0 = 429$
$n_2 = 33$
0
2
Genotype

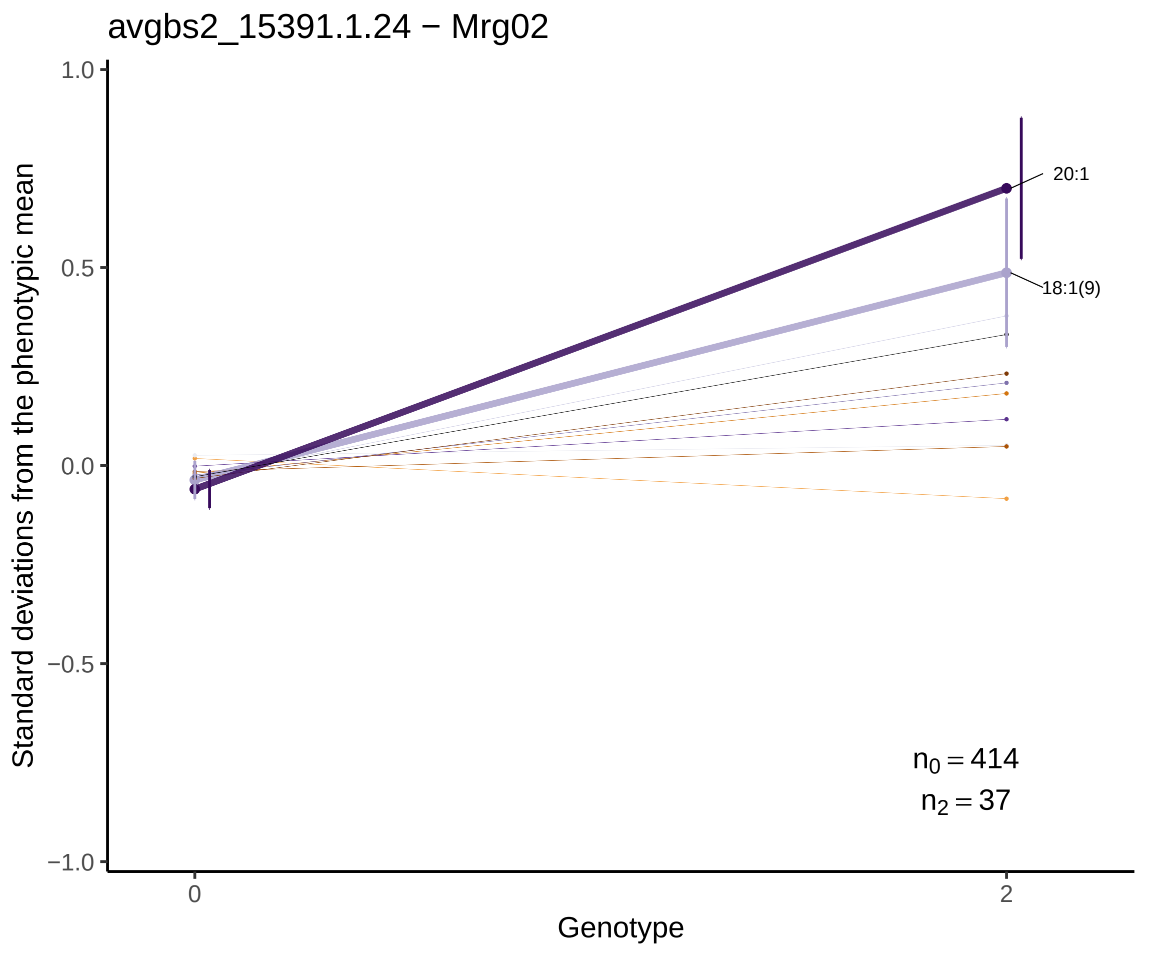
avgbs2_15391.1.24 − Mrg02
Standard deviations from the phenotypic mean
1.0
0.5
0.0
−0.5
−1.0
20:1
18:1(9)
$n_0 = 414$
$n_2 = 37$
0
2
Genotype

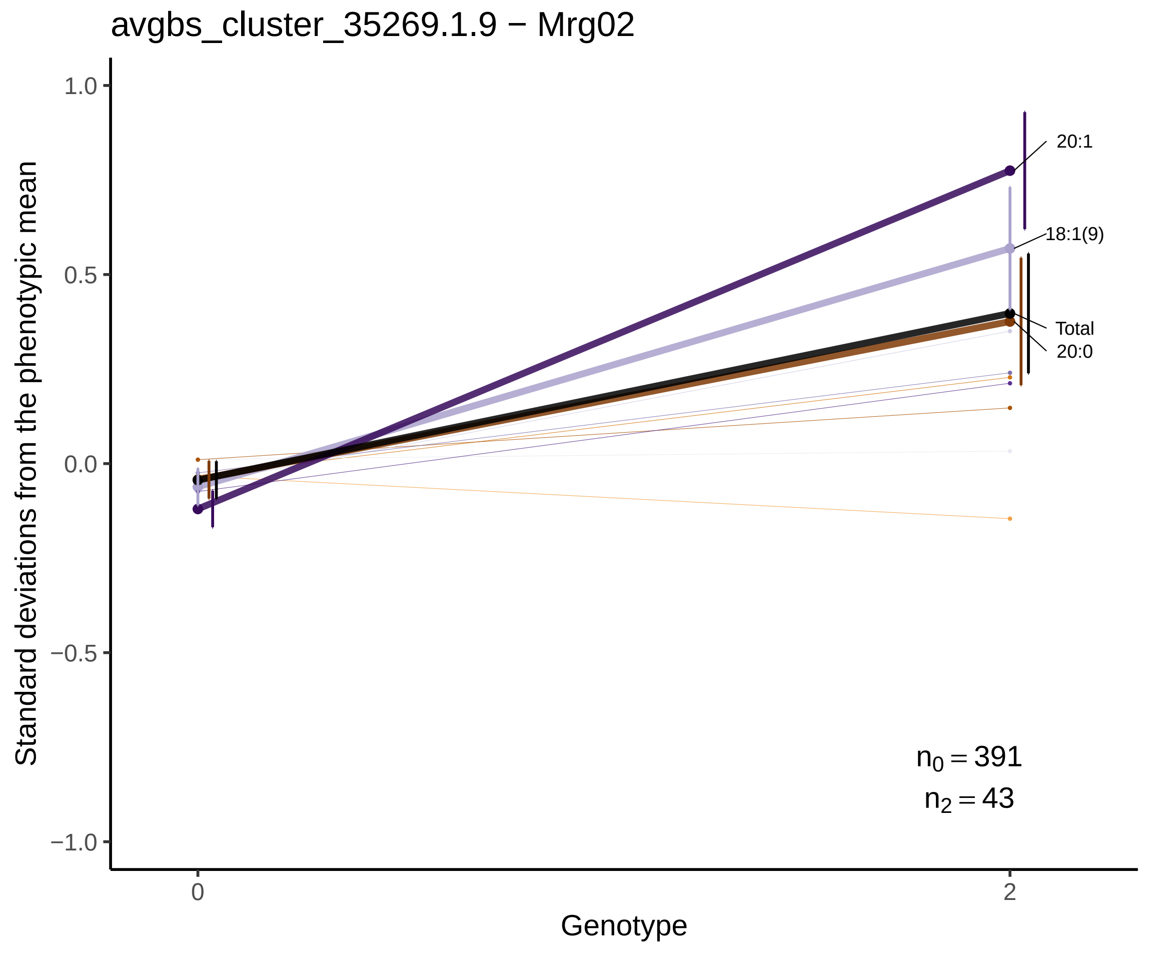
avgbs_cluster_35269.1.9 − Mrg02
Standard deviations from the phenotypic mean
1.0
0.5
0.0
−0.5
−1.0
20:1
18:1(9)
Total
20:0
$n_0 = 391$
$n_2 = 43$
0
2
Genotype

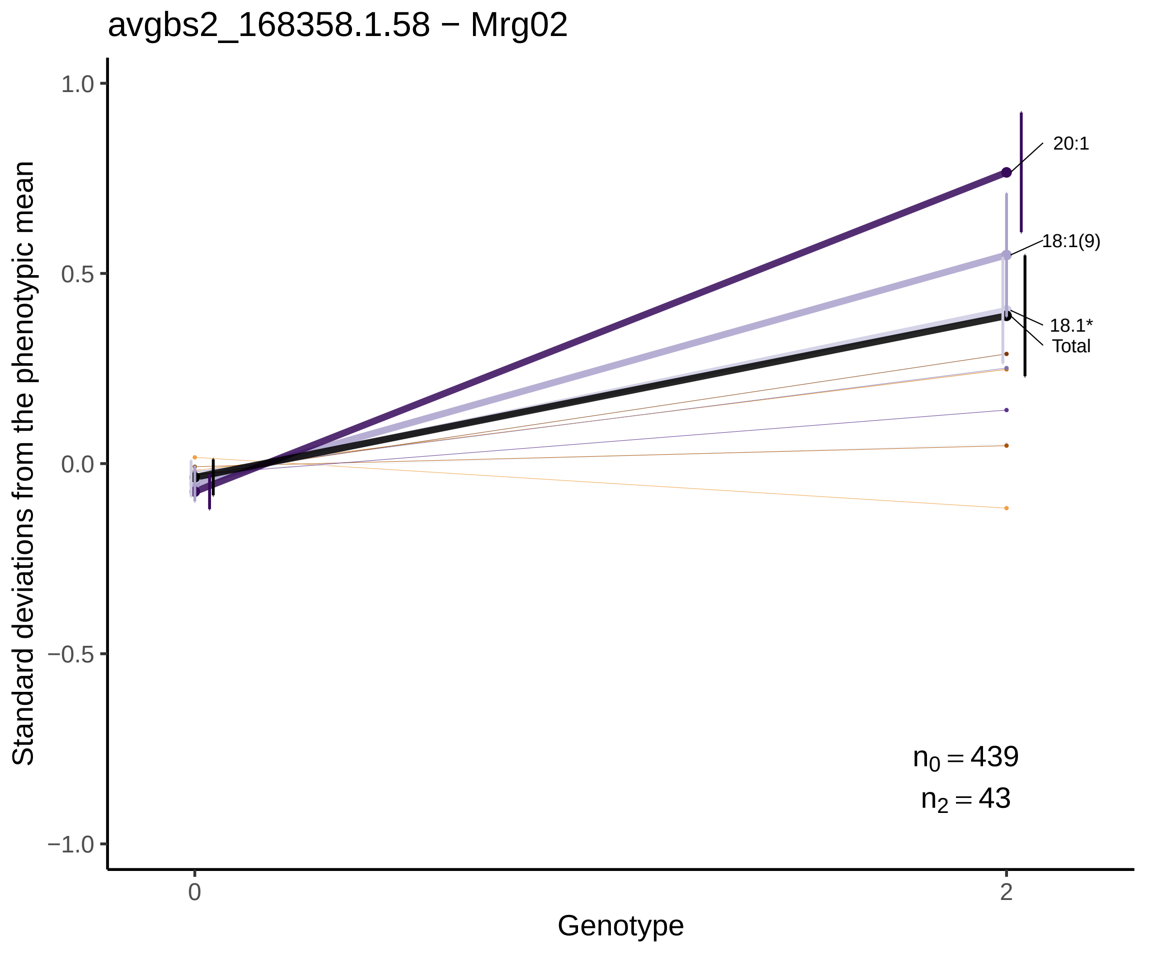
avgbs2_168358.1.58 − Mrg02
Standard deviations from the phenotypic mean
1.0
0.5
0.0
−0.5
−1.0
20:1
18:1(9)
18.1*
Total
$n_0 = 439$
$n_2 = 43$
0
2
Genotype

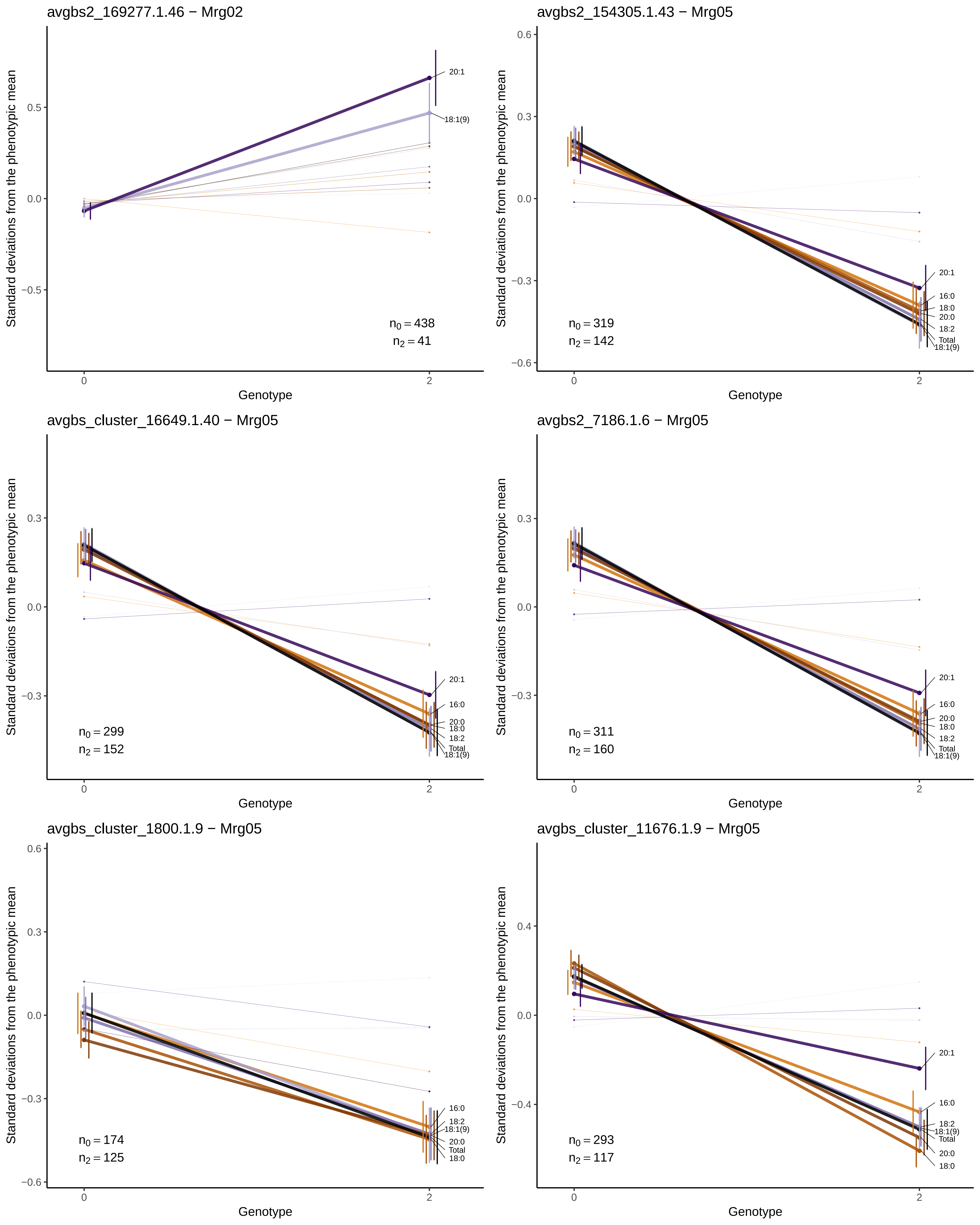

avgbs2_169277.1.46 − Mrg02
Standard deviations from the phenotypic mean
0.5
0.0
−0.5
20:1
18:1(9)
$n_0 = 438$
$n_2 = 41$
0
2
Genotype
avgbs2_154305.1.43 − Mrg05
Standard deviations from the phenotypic mean
0.6
0.3
0.0
−0.3
−0.6
20:1
16:0
18:0
20:0
18:2
Total
18:1(9)
$n_0 = 319$
$n_2 = 142$
0
2
Genotype
avgbs_cluster_16649.1.40 − Mrg05
Standard deviations from the phenotypic mean
0.3
0.0
−0.3
20:1
16:0
20:0
18:0
18:2
Total
18:1(9)
$n_0 = 299$
$n_2 = 152$
0
2
Genotype
avgbs2_7186.1.6 − Mrg05
Standard deviations from the phenotypic mean
0.3
0.0
−0.3
20:1
16:0
20:0
18:0
18:2
Total
18:1(9)
$n_0 = 311$
$n_2 = 160$
0
2
Genotype
avgbs_cluster_1800.1.9 − Mrg05
Standard deviations from the phenotypic mean
0.6
0.3
0.0
−0.3
−0.6
16:0
18:2
18:1(9)
20:0
Total
18:0
$n_0 = 174$
$n_2 = 125$
0
2
Genotype
avgbs_cluster_11676.1.9 − Mrg05
Standard deviations from the phenotypic mean
0.4
0.0
−0.4
20:1
16:0
18:2
18:1(9)
Total
20:0
18:0
$n_0 = 293$
$n_2 = 117$
0
2
Genotype

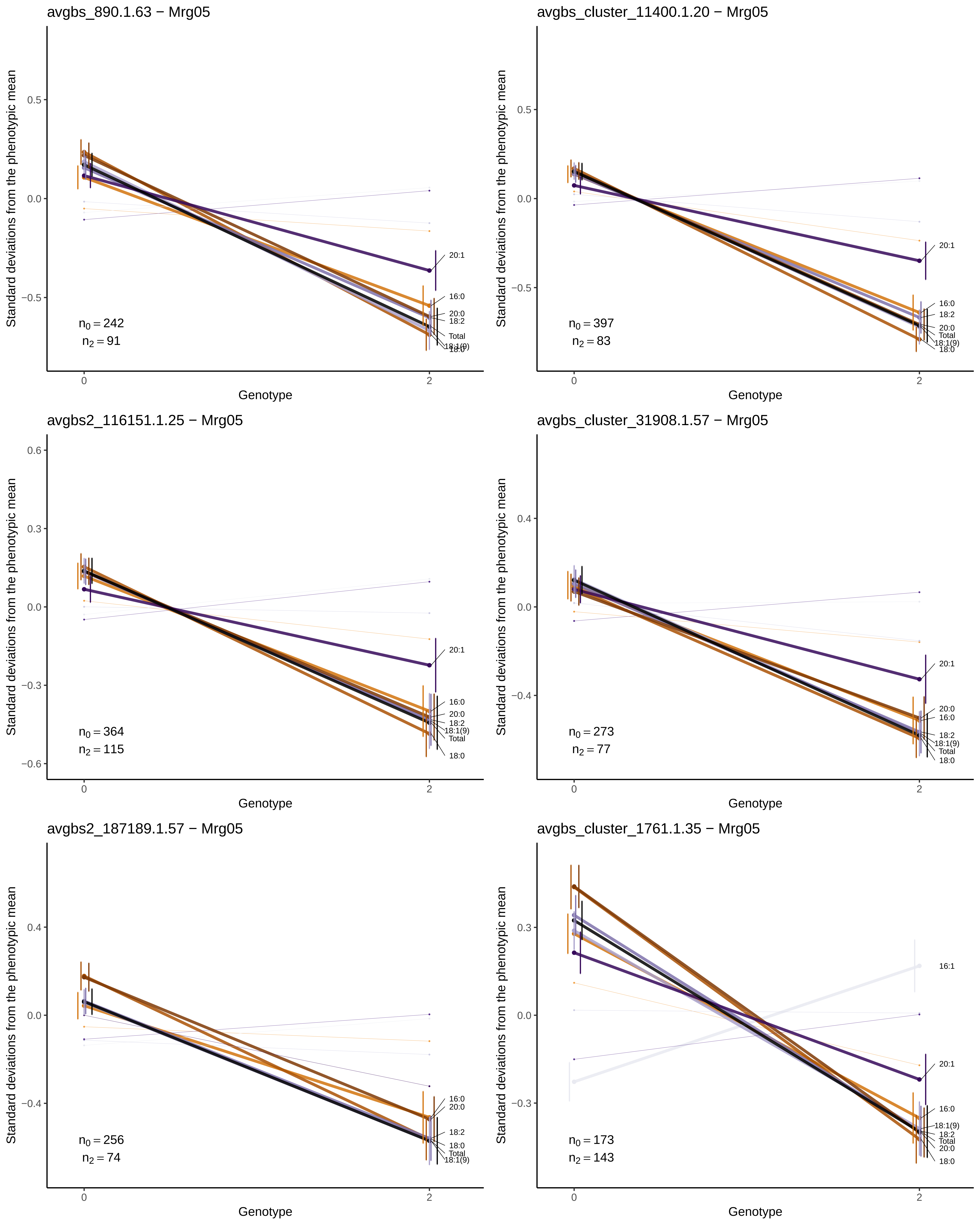

avgbs_890.1.63 − Mrg05
Standard deviations from the phenotypic mean
0.5
0.0
−0.5
20:1
16:0
20:0
18:2
Total
18:1(9)
18:0
$n_0 = 242$
$n_2 = 91$
0
2
Genotype
avgbs_cluster_11400.1.20 − Mrg05
Standard deviations from the phenotypic mean
0.5
0.0
−0.5
20:1
16:0
18:2
20:0
Total
18:1(9)
18:0
$n_0 = 397$
$n_2 = 83$
0
2
Genotype
avgbs2_116151.1.25 − Mrg05
Standard deviations from the phenotypic mean
0.6
0.3
0.0
−0.3
−0.6
20:1
16:0
20:0
18:2
18:1(9)
Total
18:0
$n_0 = 364$
$n_2 = 115$
0
2
Genotype
avgbs_cluster_31908.1.57 − Mrg05
Standard deviations from the phenotypic mean
0.4
0.0
−0.4
20:1
20:0
16:0
18:2
18:1(9)
Total
18:0
$n_0 = 273$
$n_2 = 77$
0
2
Genotype
avgbs2_187189.1.57 − Mrg05
Standard deviations from the phenotypic mean
0.4
0.0
−0.4
16:0
20:0
18:2
18:0
Total
18:1(9)
$n_0 = 256$
$n_2 = 74$
0
2
Genotype
avgbs_cluster_1761.1.35 − Mrg05
Standard deviations from the phenotypic mean
0.3
0.0
−0.3
16:1
20:1
16:0
18:1(9)
18:2
Total
20:0
18:0
$n_0 = 173$
$n_2 = 143$
0
2
Genotype

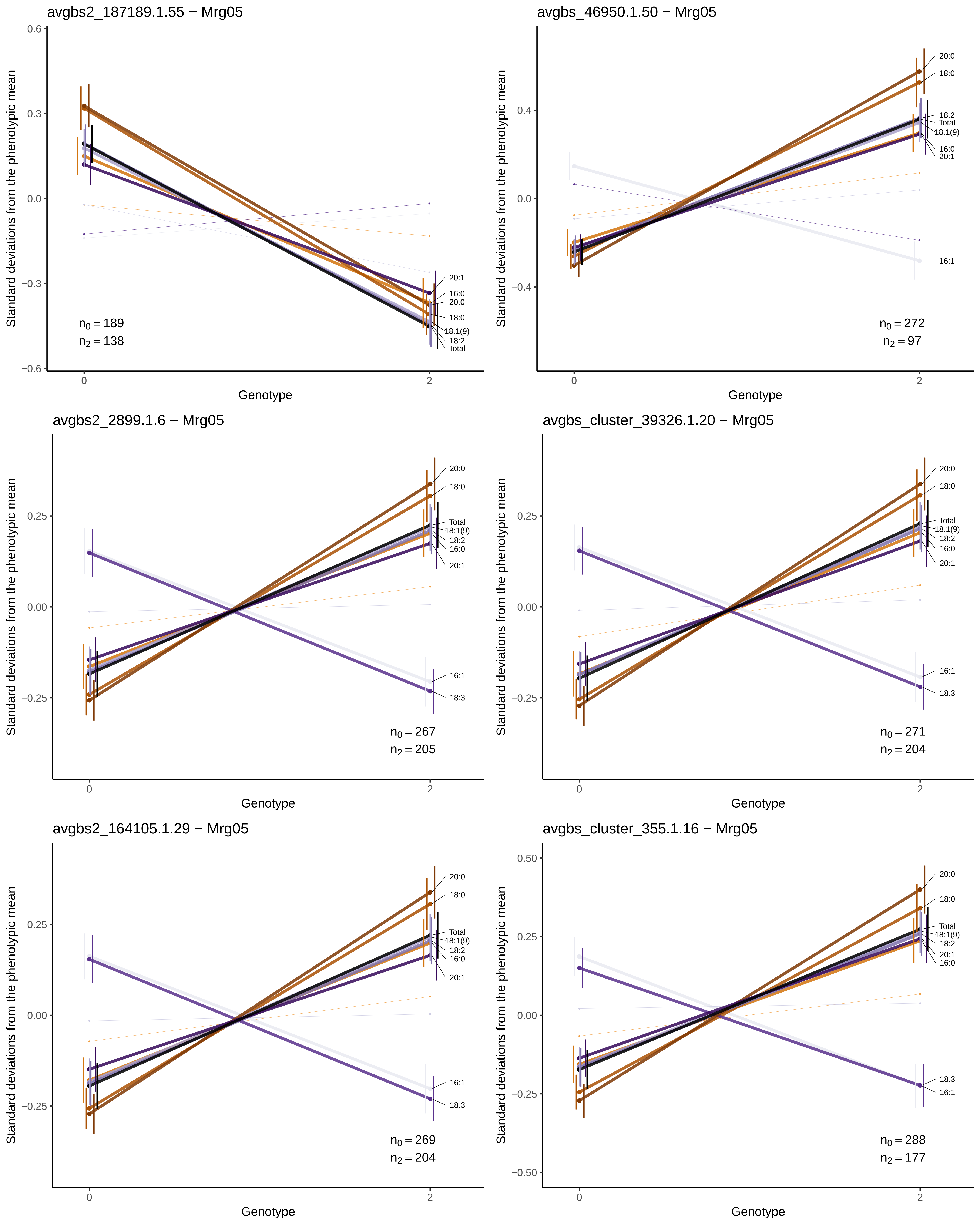
avgbs2_187189.1.55 − Mrg05
Standard deviations from the phenotypic mean
0.6
0.3
0.0
−0.3
−0.6
20:1
16:0
20:0
18:0
18:1(9)
18:2
Total
$n_0 = 189$
$n_2 = 138$
0
2
Genotype
avgbs_46950.1.50 − Mrg05
Standard deviations from the phenotypic mean
0.4
0.0
−0.4
20:0
18:0
18:2
Total
18:1(9)
16:0
20:1
16:1
$n_0 = 272$
$n_2 = 97$
0
2
Genotype
avgbs2_2899.1.6 − Mrg05
Standard deviations from the phenotypic mean
0.25
0.00
−0.25
20:0
18:0
Total
18:1(9)
18:2
16:0
20:1
16:1
18:3
$n_0 = 267$
$n_2 = 205$
0
2
Genotype
avgbs_cluster_39326.1.20 − Mrg05
Standard deviations from the phenotypic mean
0.25
0.00
−0.25
20:0
18:0
Total
18:1(9)
18:2
16:0
20:1
16:1
18:3
$n_0 = 271$
$n_2 = 204$
0
2
Genotype
avgbs2_164105.1.29 − Mrg05
Standard deviations from the phenotypic mean
0.25
0.00
−0.25
20:0
18:0
Total
18:1(9)
18:2
16:0
20:1
16:1
18:3
$n_0 = 269$
$n_2 = 204$
0
2
Genotype
avgbs_cluster_355.1.16 − Mrg05
Standard deviations from the phenotypic mean
0.50
0.25
0.00
−0.25
−0.50
20:0
18:0
Total
18:1(9)
18:2
20:1
16:0
18:3
16:1
$n_0 = 288$
$n_2 = 177$
0
2
Genotype

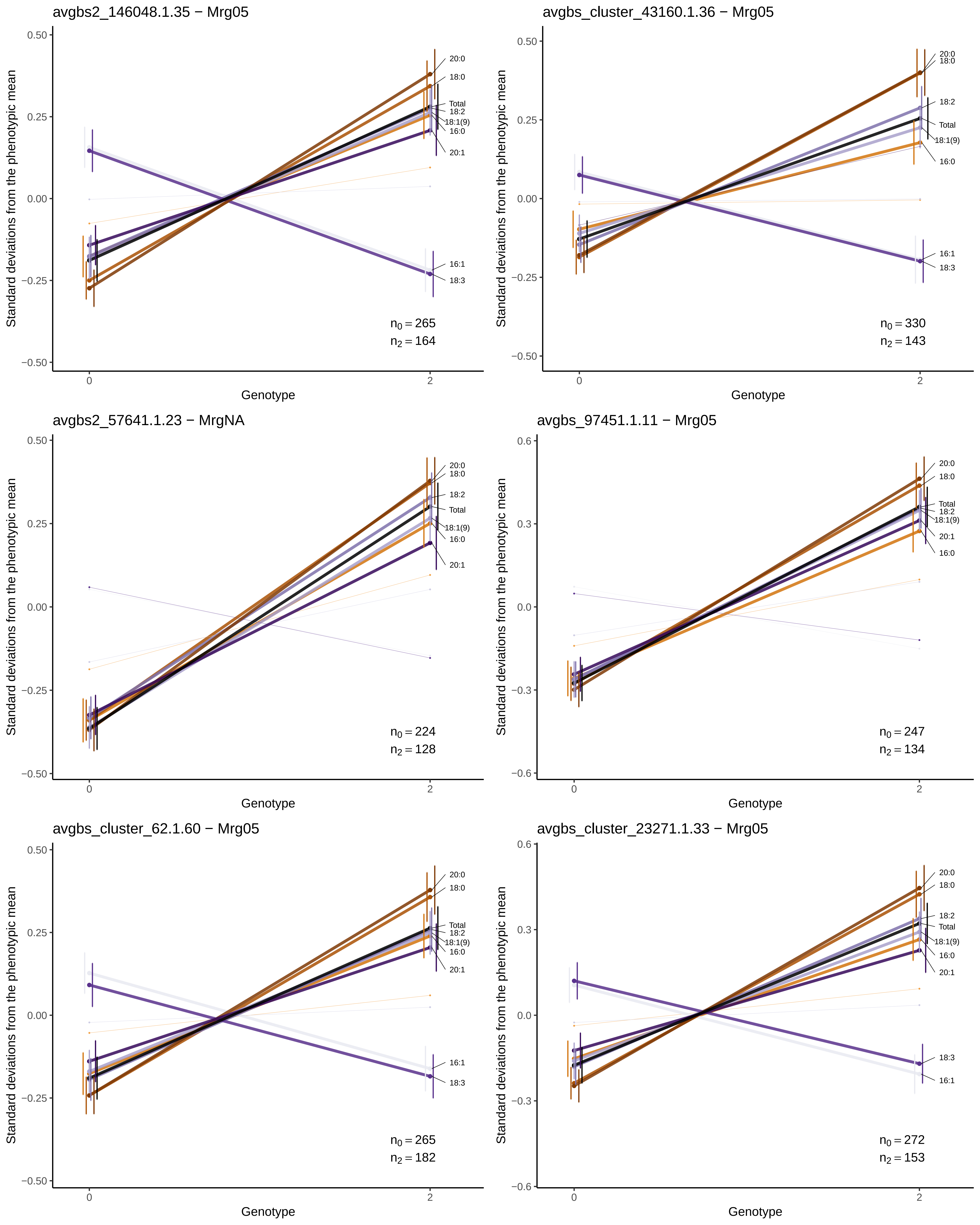

avgbs2_146048.1.35 − Mrg05
Standard deviations from the phenotypic mean
0.50
0.25
0.00
−0.25
−0.50
Genotype
0
2
20:0
18:0
Total
18:2
18:1(9)
16:0
20:1
16:1
18:3
$n_0 = 265$
$n_2 = 164$
avgbs_cluster_43160.1.36 − Mrg05
Standard deviations from the phenotypic mean
0.50
0.25
0.00
−0.25
−0.50
Genotype
0
2
20:0
18:0
18:2
Total
18:1(9)
16:0
16:1
18:3
$n_0 = 330$
$n_2 = 143$
avgbs2_57641.1.23 − MrgNA
Standard deviations from the phenotypic mean
0.50
0.25
0.00
−0.25
−0.50
Genotype
0
2
20:0
18:0
18:2
Total
18:1(9)
16:0
20:1
$n_0 = 224$
$n_2 = 128$
avgbs_97451.1.11 − Mrg05
Standard deviations from the phenotypic mean
0.6
0.3
0.0
−0.3
−0.6
Genotype
0
2
20:0
18:0
Total
18:2
18:1(9)
20:1
16:0
$n_0 = 247$
$n_2 = 134$
avgbs_cluster_62.1.60 − Mrg05
Standard deviations from the phenotypic mean
0.50
0.25
0.00
−0.25
−0.50
Genotype
0
2
20:0
18:0
Total
18:2
18:1(9)
16:0
20:1
16:1
18:3
$n_0 = 265$
$n_2 = 182$
avgbs_cluster_23271.1.33 − Mrg05
Standard deviations from the phenotypic mean
0.6
0.3
0.0
−0.3
−0.6
Genotype
0
2
20:0
18:0
18:2
Total
18:1(9)
16:0
20:1
18:3
16:1
$n_0 = 272$
$n_2 = 153$

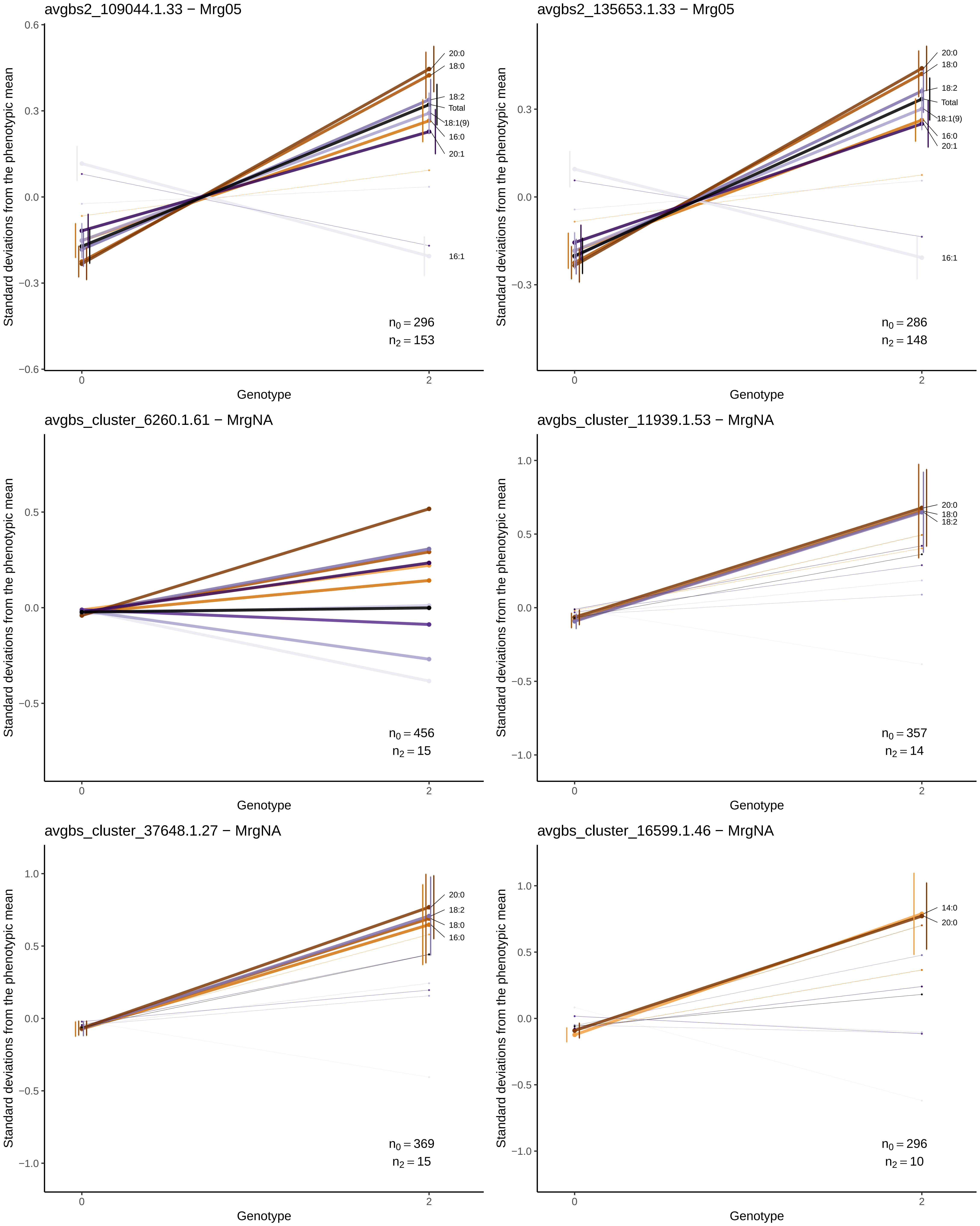

avgbs2_109044.1.33 − Mrg05
Standard deviations from the phenotypic mean
Genotype
20:0
18:0
18:2
Total
18:1(9)
16:0
20:1
16:1
$n_0 = 296$
$n_2 = 153$
avgbs2_135653.1.33 − Mrg05
Standard deviations from the phenotypic mean
Genotype
20:0
18:0
18:2
Total
18:1(9)
16:0
20:1
16:1
$n_0 = 286$
$n_2 = 148$
avgbs_cluster_6260.1.61 − MrgNA
Standard deviations from the phenotypic mean
Genotype
$n_0 = 456$
$n_2 = 15$
avgbs_cluster_11939.1.53 − MrgNA
Standard deviations from the phenotypic mean
Genotype
20:0
18:0
18:2
$n_0 = 357$
$n_2 = 14$
avgbs_cluster_37648.1.27 − MrgNA
Standard deviations from the phenotypic mean
Genotype
20:0
18:2
18:0
16:0
$n_0 = 369$
$n_2 = 15$
avgbs_cluster_16599.1.46 − MrgNA
Standard deviations from the phenotypic mean
Genotype
14:0
20:0
$n_0 = 296$
$n_2 = 10$

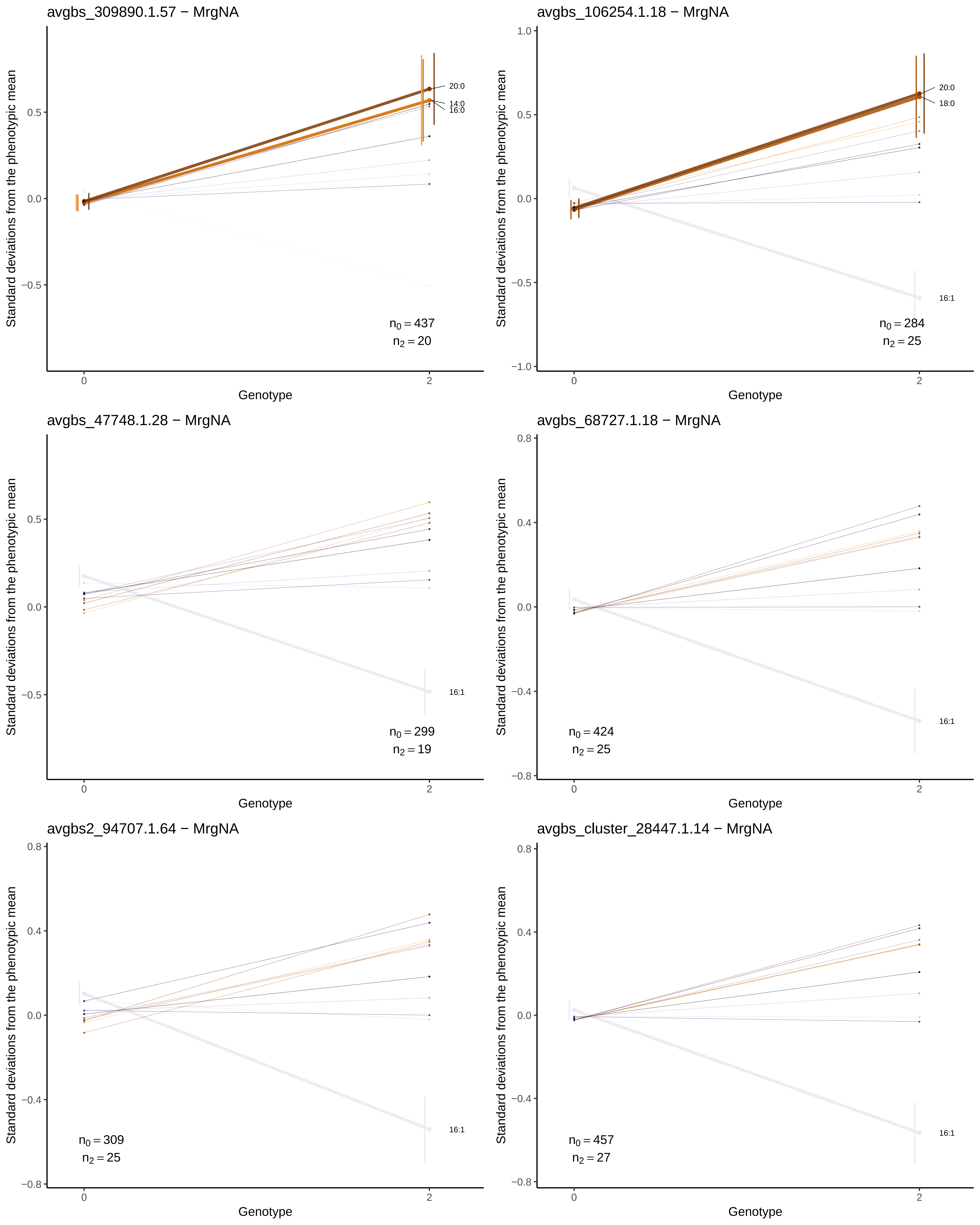
avgbs_309890.1.57 − MrgNA
Standard deviations from the phenotypic mean
0.5
0.0
−0.5
20:0
14:0
16:0
$n_0 = 437$
$n_2 = 20$
0
2
Genotype
avgbs_106254.1.18 − MrgNA
Standard deviations from the phenotypic mean
1.0
0.5
0.0
−0.5
−1.0
20:0
18:0
16:1
$n_0 = 284$
$n_2 = 25$
0
2
Genotype
avgbs_47748.1.28 − MrgNA
Standard deviations from the phenotypic mean
0.5
0.0
−0.5
16:1
$n_0 = 299$
$n_2 = 19$
0
2
Genotype
avgbs_68727.1.18 − MrgNA
Standard deviations from the phenotypic mean
0.8
0.4
0.0
−0.4
−0.8
16:1
$n_0 = 424$
$n_2 = 25$
0
2
Genotype
avgbs2_94707.1.64 − MrgNA
Standard deviations from the phenotypic mean
0.8
0.4
0.0
−0.4
−0.8
16:1
$n_0 = 309$
$n_2 = 25$
0
2
Genotype
avgbs_cluster_28447.1.14 − MrgNA
Standard deviations from the phenotypic mean
0.8
0.4
0.0
−0.4
−0.8
16:1
$n_0 = 457$
$n_2 = 27$
0
2
Genotype

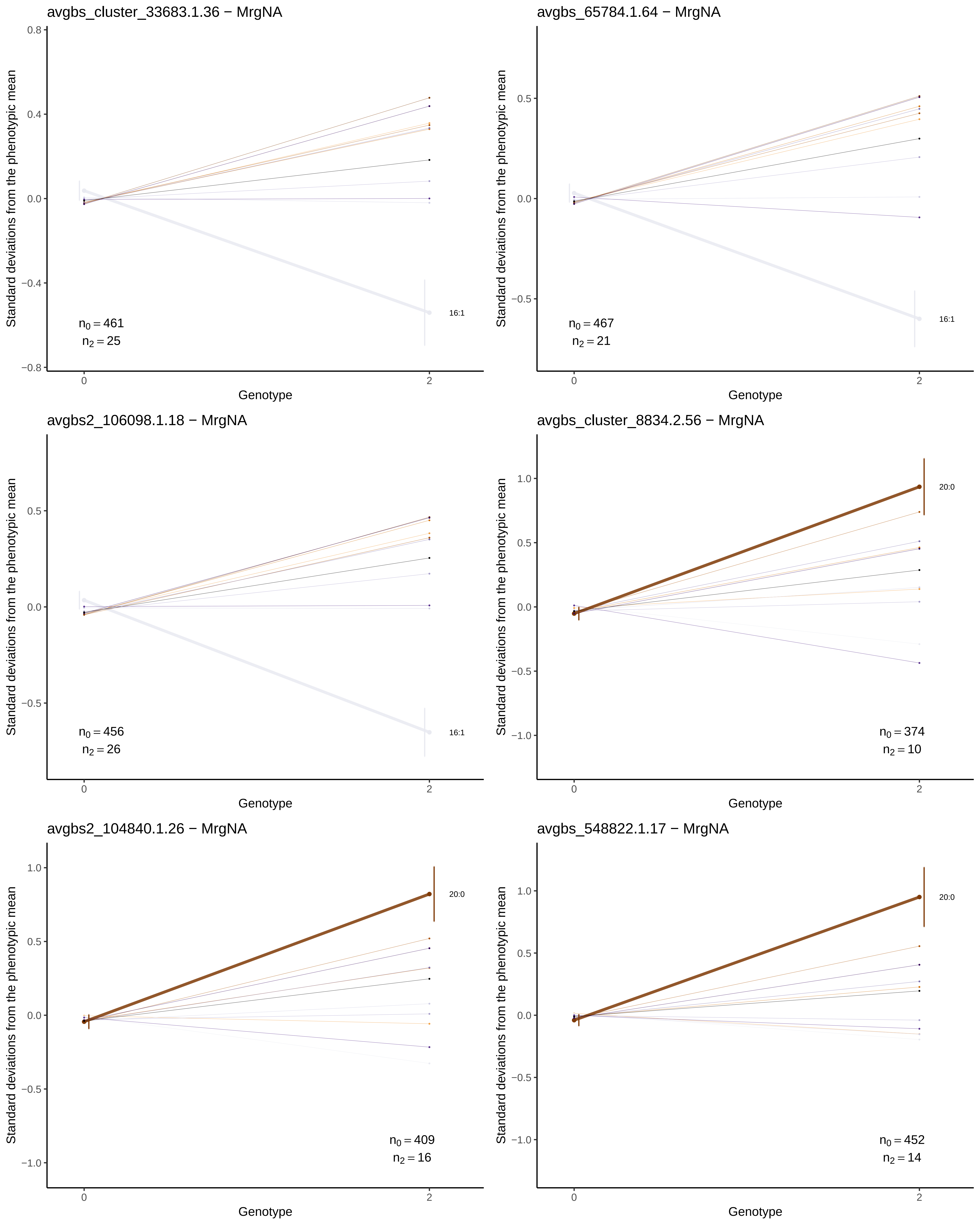

avgbs_cluster_33683.1.36 − MrgNA
16:1
$n_0 = 461$
$n_2 = 25$
avgbs_65784.1.64 − MrgNA
16:1
$n_0 = 467$
$n_2 = 21$
avgbs2_106098.1.18 − MrgNA
16:1
$n_0 = 456$
$n_2 = 26$
avgbs_cluster_8834.2.56 − MrgNA
20:0
$n_0 = 374$
$n_2 = 10$
avgbs2_104840.1.26 − MrgNA
20:0
$n_0 = 409$
$n_2 = 16$
avgbs_548822.1.17 − MrgNA
20:0
$n_0 = 452$
$n_2 = 14$
Standard deviations from the phenotypic mean
Genotype
0
2

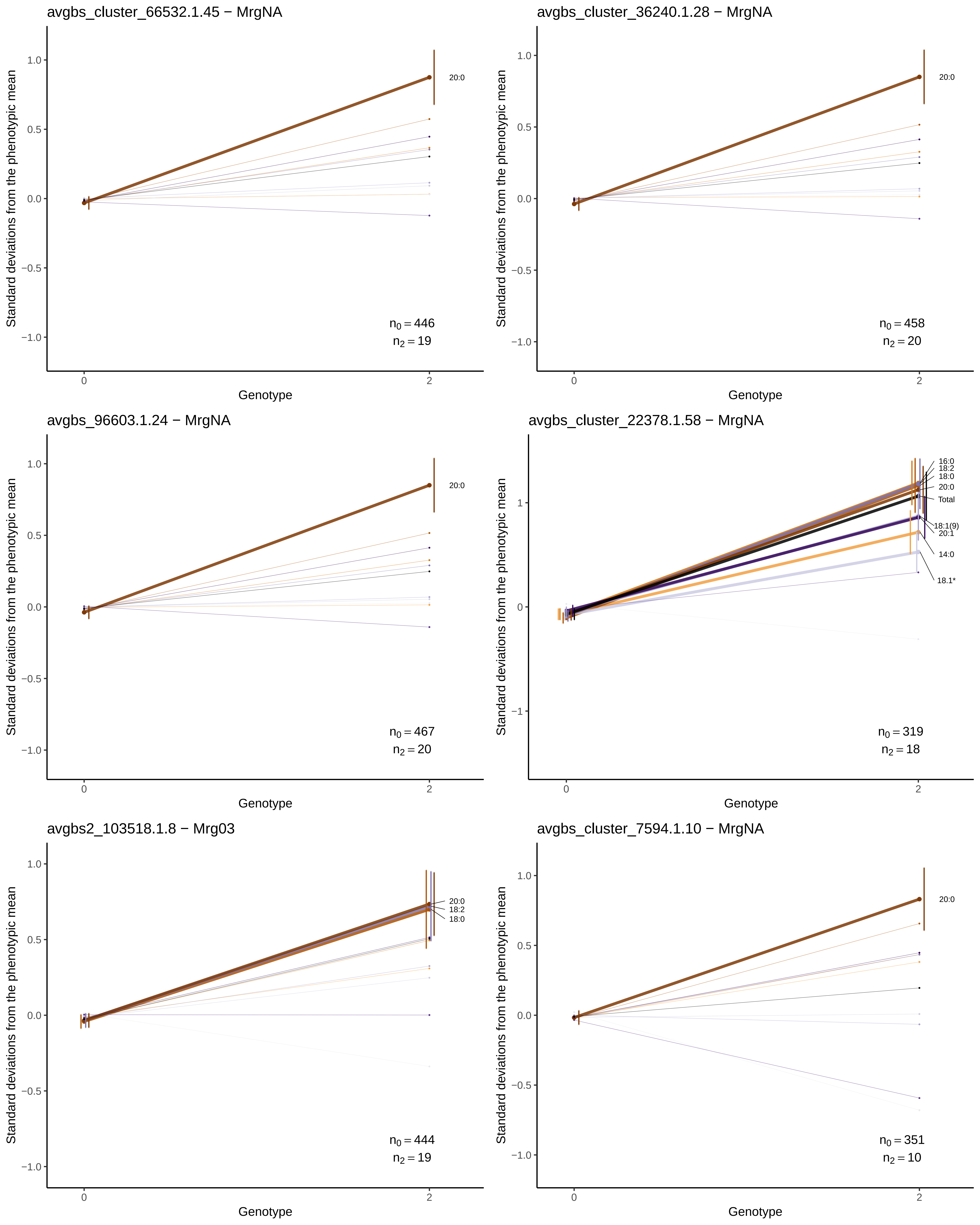
avgbs_cluster_66532.1.45 − MrgNA
20:0
n0 = 446
n2 = 19
avgbs_cluster_36240.1.28 − MrgNA
20:0
n0 = 458
n2 = 20
avgbs_96603.1.24 − MrgNA
20:0
n0 = 467
n2 = 20
avgbs_cluster_22378.1.58 − MrgNA
16:0
18:2
18:0
20:0
Total
18:1(9)
20:1
14:0
18:1*
n0 = 319
n2 = 18
avgbs2_103518.1.8 − Mrg03
20:0
18:2
18:0
n0 = 444
n2 = 19
avgbs_cluster_7594.1.10 − MrgNA
20:0
n0 = 351
n2 = 10
Standard deviations from the phenotypic mean
Genotype

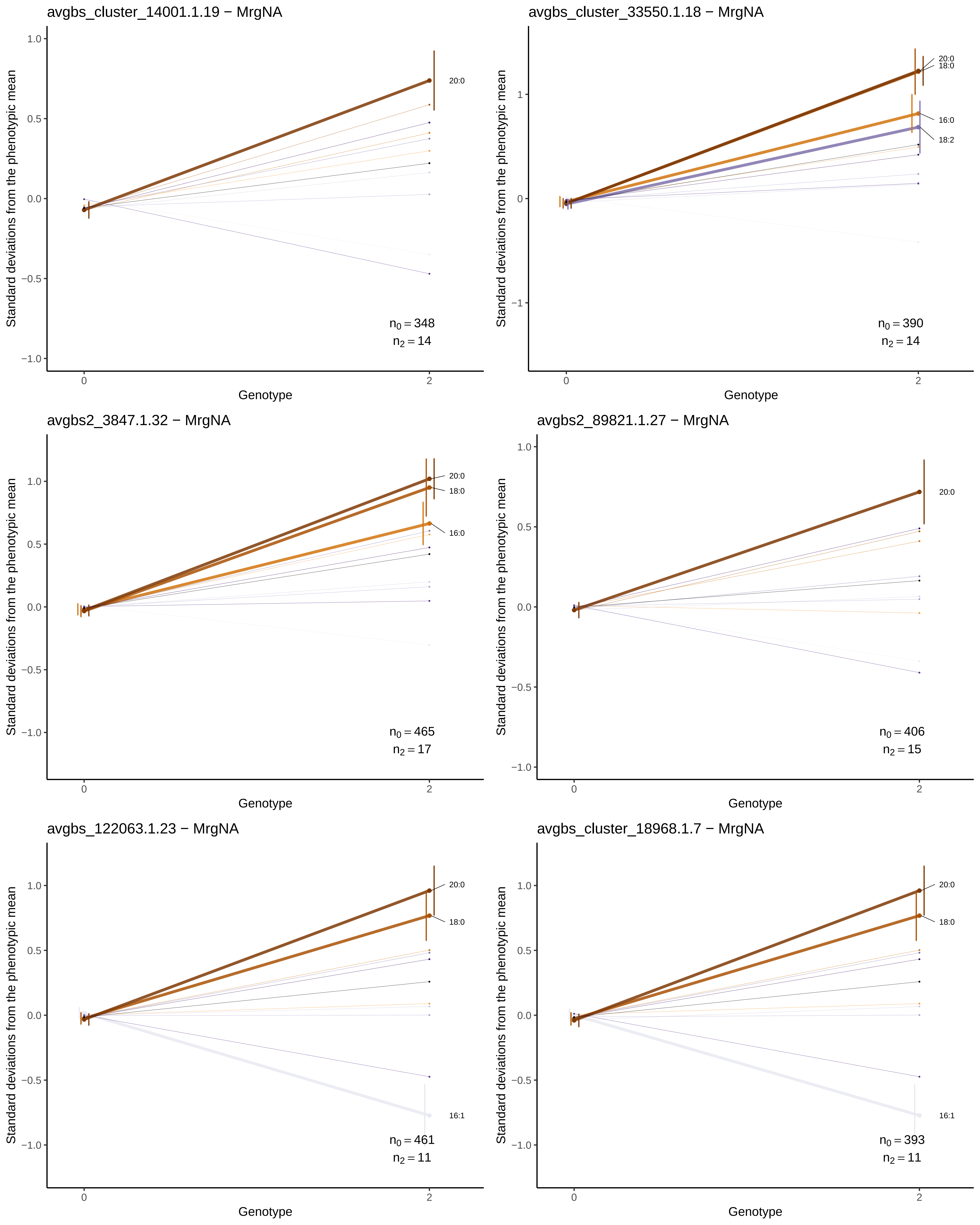

avgbs_cluster_14001.1.19 − MrgNA
Standard deviations from the phenotypic mean
Genotype
20:0
$n_0 = 348$
$n_2 = 14$
avgbs_cluster_33550.1.18 − MrgNA
Standard deviations from the phenotypic mean
Genotype
20:0
18:0
16:0
18:2
$n_0 = 390$
$n_2 = 14$
avgbs2_3847.1.32 − MrgNA
Standard deviations from the phenotypic mean
Genotype
20:0
18:0
16:0
$n_0 = 465$
$n_2 = 17$
avgbs2_89821.1.27 − MrgNA
Standard deviations from the phenotypic mean
Genotype
20:0
$n_0 = 406$
$n_2 = 15$
avgbs_122063.1.23 − MrgNA
Standard deviations from the phenotypic mean
Genotype
20:0
18:0
16:1
$n_0 = 461$
$n_2 = 11$
avgbs_cluster_18968.1.7 − MrgNA
Standard deviations from the phenotypic mean
Genotype
20:0
18:0
16:1
$n_0 = 393$
$n_2 = 11$

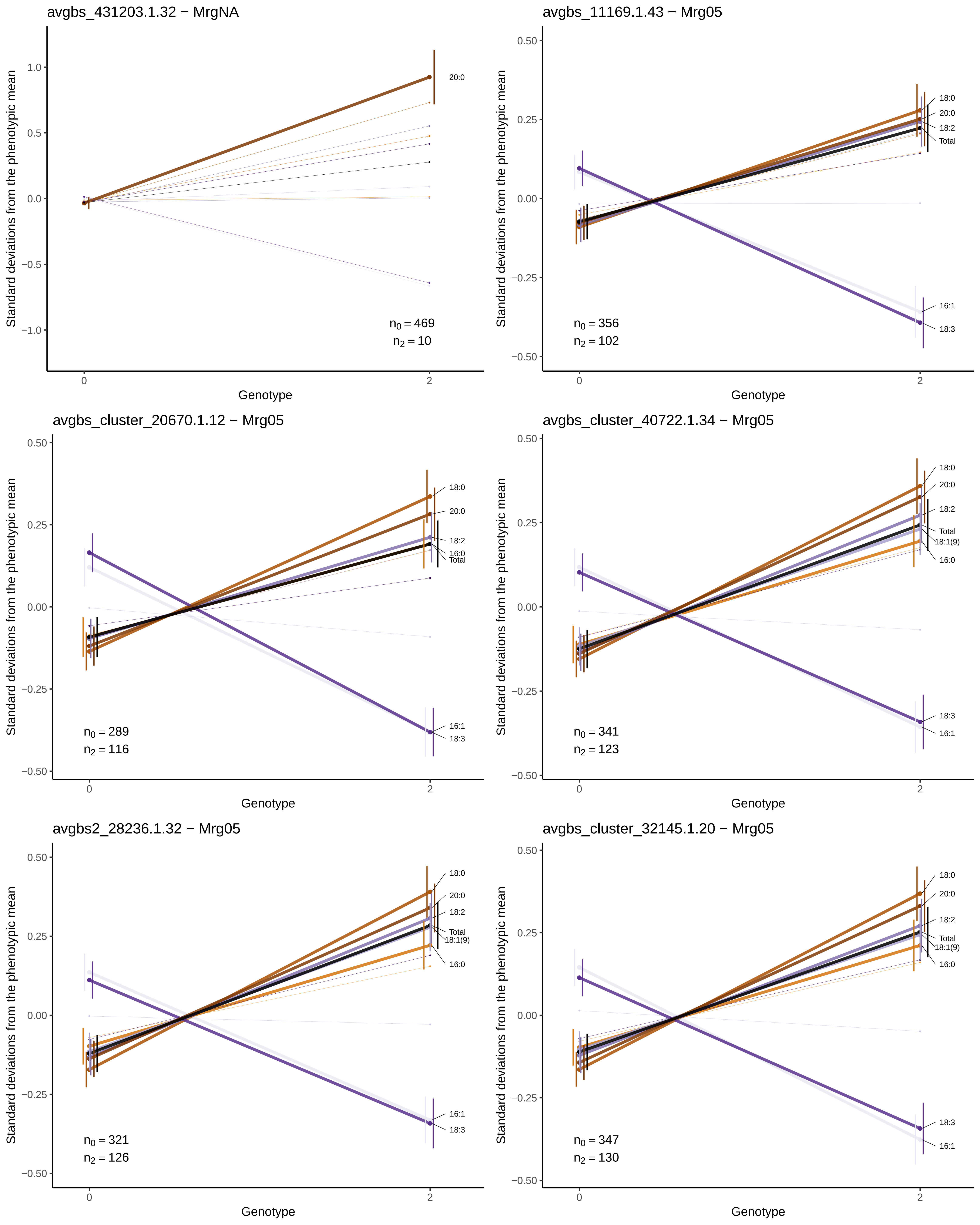
avgbs_431203.1.32 − MrgNA
Standard deviations from the phenotypic mean
1.0
0.5
0.0
−0.5
−1.0
20:0
$n_0 = 469$
$n_2 = 10$
0
2
Genotype
avgbs_11169.1.43 − Mrg05
Standard deviations from the phenotypic mean
0.50
0.25
0.00
−0.25
−0.50
18:0
20:0
18:2
Total
16:1
18:3
$n_0 = 356$
$n_2 = 102$
0
2
Genotype
avgbs_cluster_20670.1.12 − Mrg05
Standard deviations from the phenotypic mean
0.50
0.25
0.00
−0.25
−0.50
18:0
20:0
18:2
16:0
Total
16:1
18:3
$n_0 = 289$
$n_2 = 116$
0
2
Genotype
avgbs_cluster_40722.1.34 − Mrg05
Standard deviations from the phenotypic mean
0.50
0.25
0.00
−0.25
−0.50
18:0
20:0
18:2
Total
18:1(9)
16:0
18:3
16:1
$n_0 = 341$
$n_2 = 123$
0
2
Genotype
avgbs2_28236.1.32 − Mrg05
Standard deviations from the phenotypic mean
0.50
0.25
0.00
−0.25
−0.50
18:0
20:0
18:2
Total
18:1(9)
16:0
16:1
18:3
$n_0 = 321$
$n_2 = 126$
0
2
Genotype
avgbs_cluster_32145.1.20 − Mrg05
Standard deviations from the phenotypic mean
0.50
0.25
0.00
−0.25
−0.50
18:0
20:0
18:2
Total
18:1(9)
16:0
18:3
16:1
$n_0 = 347$
$n_2 = 130$
0
2
Genotype

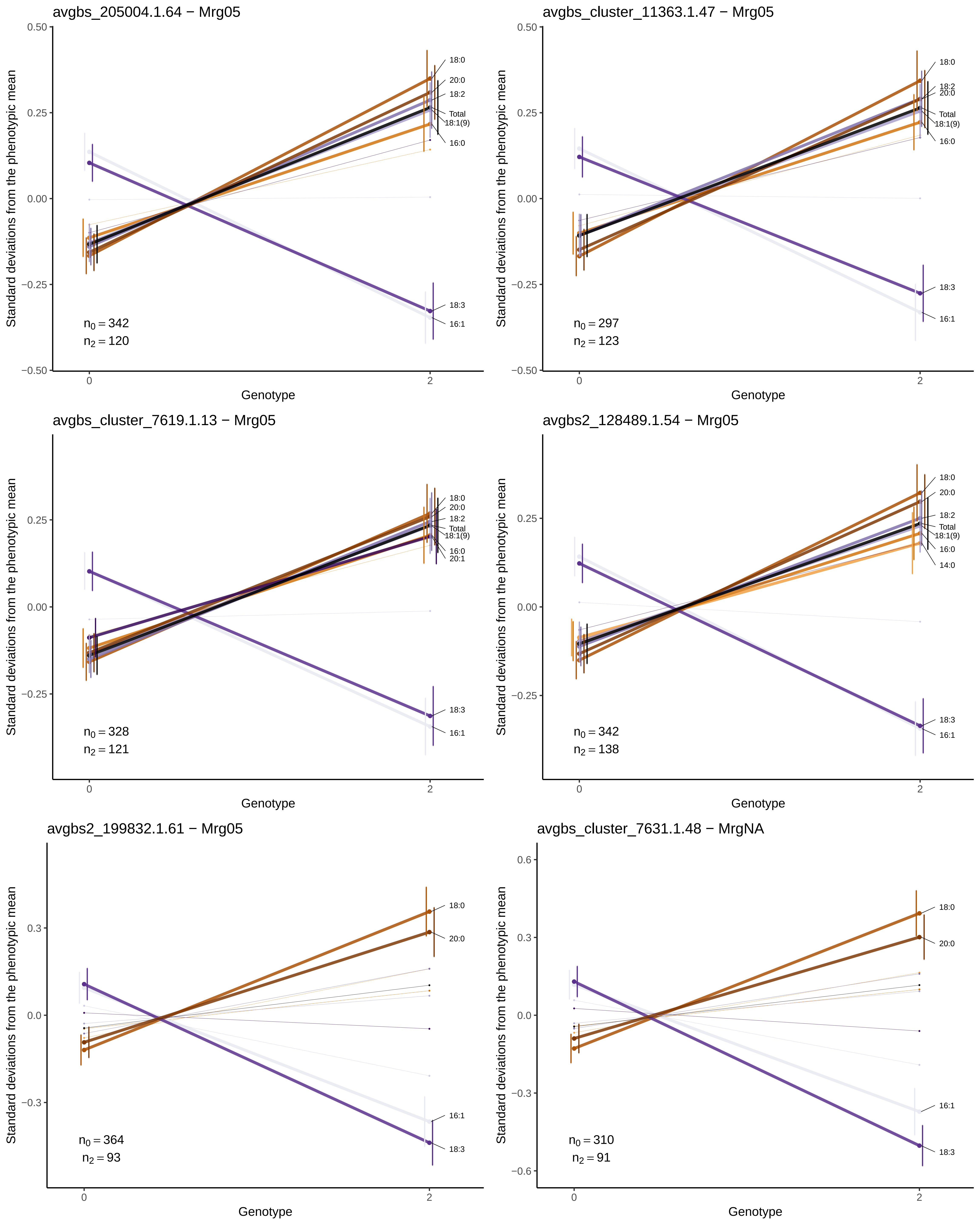
avgbs_205004.1.64 − Mrg05
18:0
20:0
18:2
Total
18:1(9)
16:0
18:3
16:1
$n_0 = 342$
$n_2 = 120$
avgbs_cluster_11363.1.47 − Mrg05
18:0
18:2
20:0
Total
18:1(9)
16:0
18:3
16:1
$n_0 = 297$
$n_2 = 123$
avgbs_cluster_7619.1.13 − Mrg05
18:0
20:0
18:2
Total
18:1(9)
16:0
20:1
18:3
16:1
$n_0 = 328$
$n_2 = 121$
avgbs2_128489.1.54 − Mrg05
18:0
20:0
18:2
Total
18:1(9)
16:0
14:0
18:3
16:1
$n_0 = 342$
$n_2 = 138$
avgbs2_199832.1.61 − Mrg05
18:0
20:0
16:1
18:3
$n_0 = 364$
$n_2 = 93$
avgbs_cluster_7631.1.48 − MrgNA
18:0
20:0
16:1
18:3
$n_0 = 310$
$n_2 = 91$
Standard deviations from the phenotypic mean
Genotype
0
2
0.50
0.25
0.00
−0.25
−0.50
0.6
0.3
0.0
−0.3
−0.6

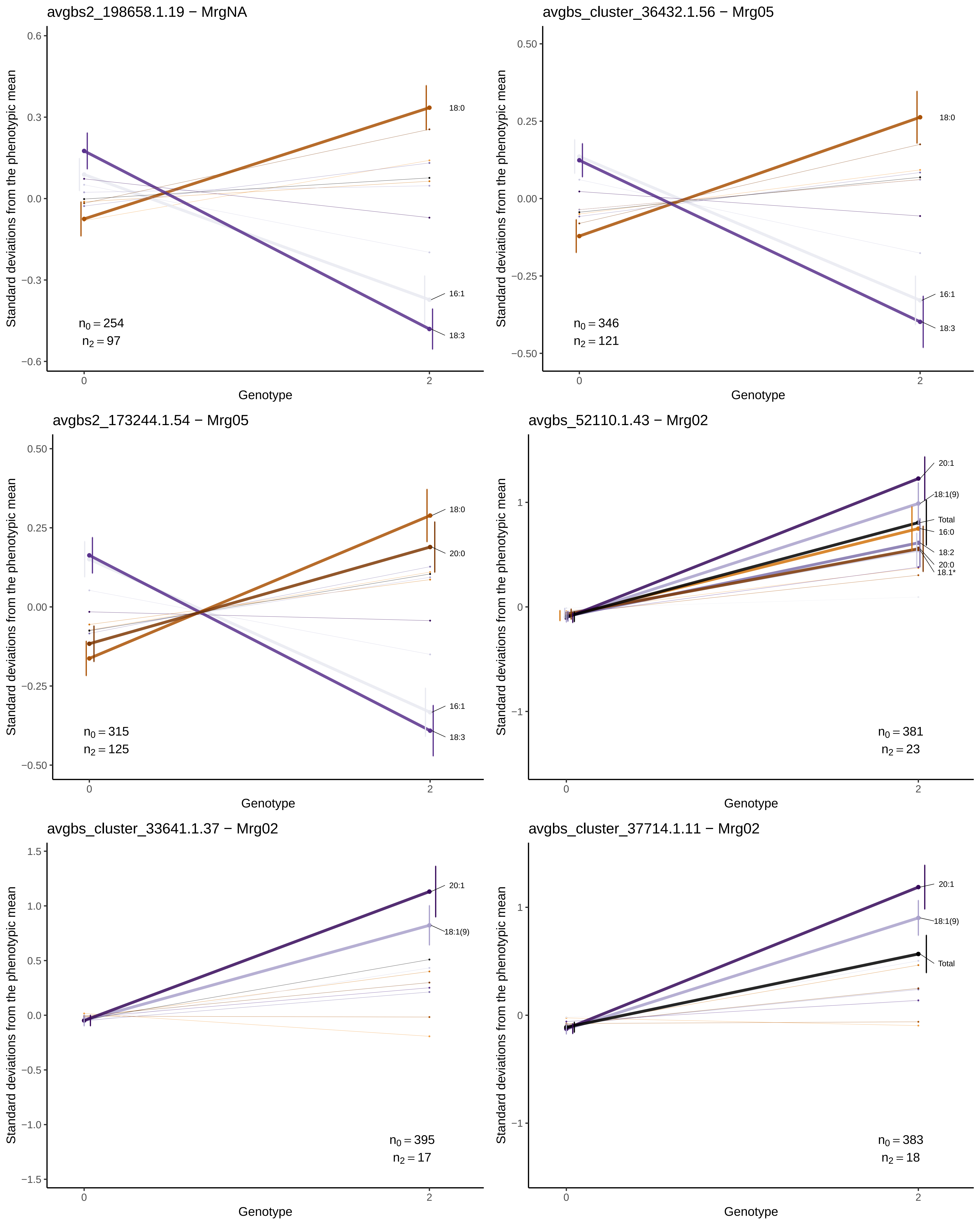
avgbs2_198658.1.19 − MrgNA
Standard deviations from the phenotypic mean
Genotype
18:0
16:1
18:3
$n_0 = 254$
$n_2 = 97$
avgbs_cluster_36432.1.56 − Mrg05
Standard deviations from the phenotypic mean
Genotype
18:0
16:1
18:3
$n_0 = 346$
$n_2 = 121$
avgbs2_173244.1.54 − Mrg05
Standard deviations from the phenotypic mean
Genotype
18:0
20:0
16:1
18:3
$n_0 = 315$
$n_2 = 125$
avgbs_52110.1.43 − Mrg02
Standard deviations from the phenotypic mean
Genotype
20:1
18:1(9)
Total
16:0
18:2
20:0
18.1*
$n_0 = 381$
$n_2 = 23$
avgbs_cluster_33641.1.37 − Mrg02
Standard deviations from the phenotypic mean
Genotype
20:1
18:1(9)
$n_0 = 395$
$n_2 = 17$
avgbs_cluster_37714.1.11 − Mrg02
Standard deviations from the phenotypic mean
Genotype
20:1
18:1(9)
Total
$n_0 = 383$
$n_2 = 18$

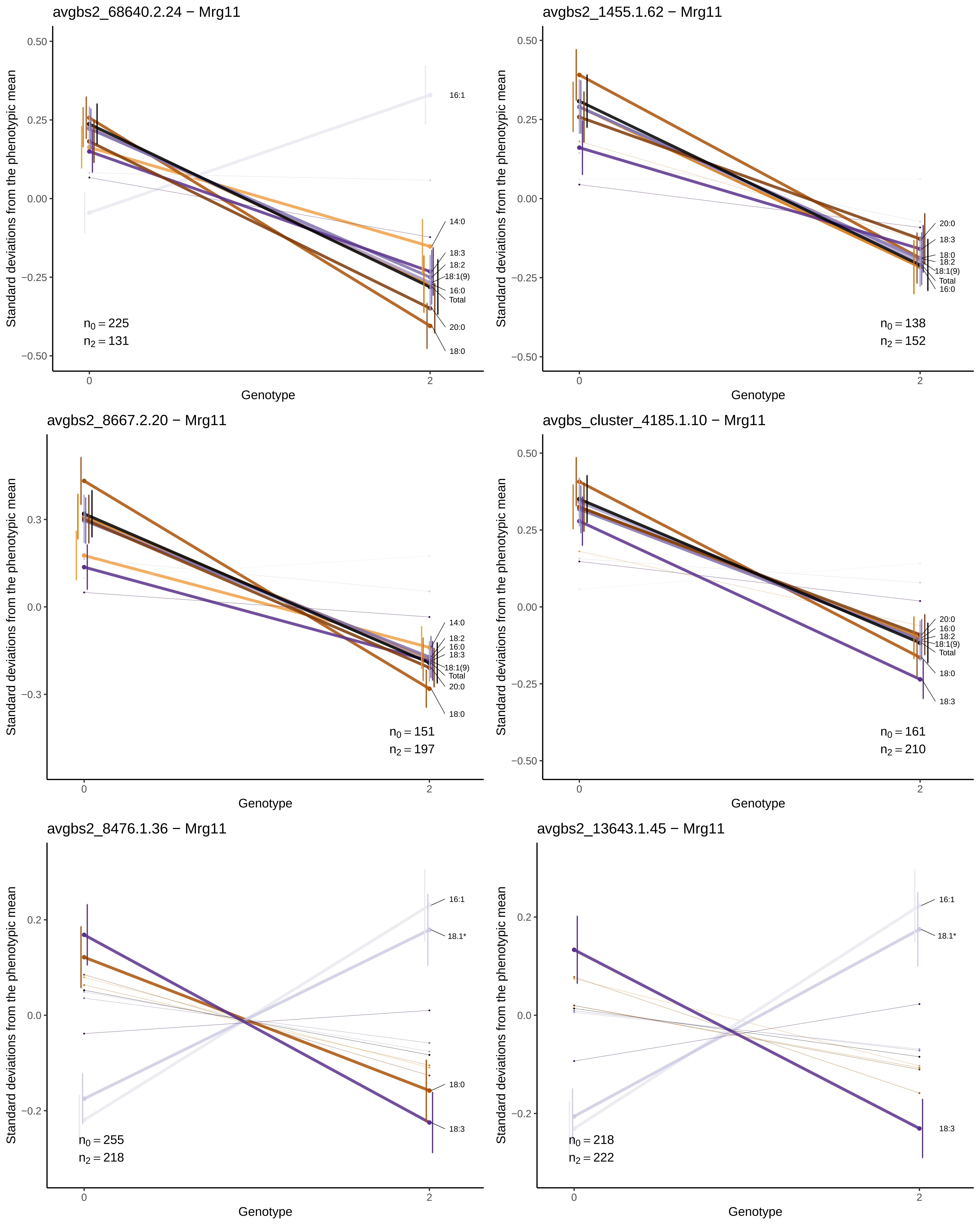

avgbs2_68640.2.24 − Mrg11
Standard deviations from the phenotypic mean
Genotype
16:1
14:0
18:3
18:2
18:1(9)
16:0
Total
20:0
18:0
$n_0 = 225$
$n_2 = 131$
avgbs2_1455.1.62 − Mrg11
20:0
18:3
18:0
18:2
18:1(9)
Total
16:0
$n_0 = 138$
$n_2 = 152$
avgbs2_8667.2.20 − Mrg11
14:0
18:2
16:0
18:3
18:1(9)
Total
20:0
18:0
$n_0 = 151$
$n_2 = 197$
avgbs_cluster_4185.1.10 − Mrg11
20:0
16:0
18:2
18:1(9)
Total
18:0
18:3
$n_0 = 161$
$n_2 = 210$
avgbs2_8476.1.36 − Mrg11
16:1
18.1*
18:0
18:3
$n_0 = 255$
$n_2 = 218$
avgbs2_13643.1.45 − Mrg11
16:1
18.1*
18:3
$n_0 = 218$
$n_2 = 222$

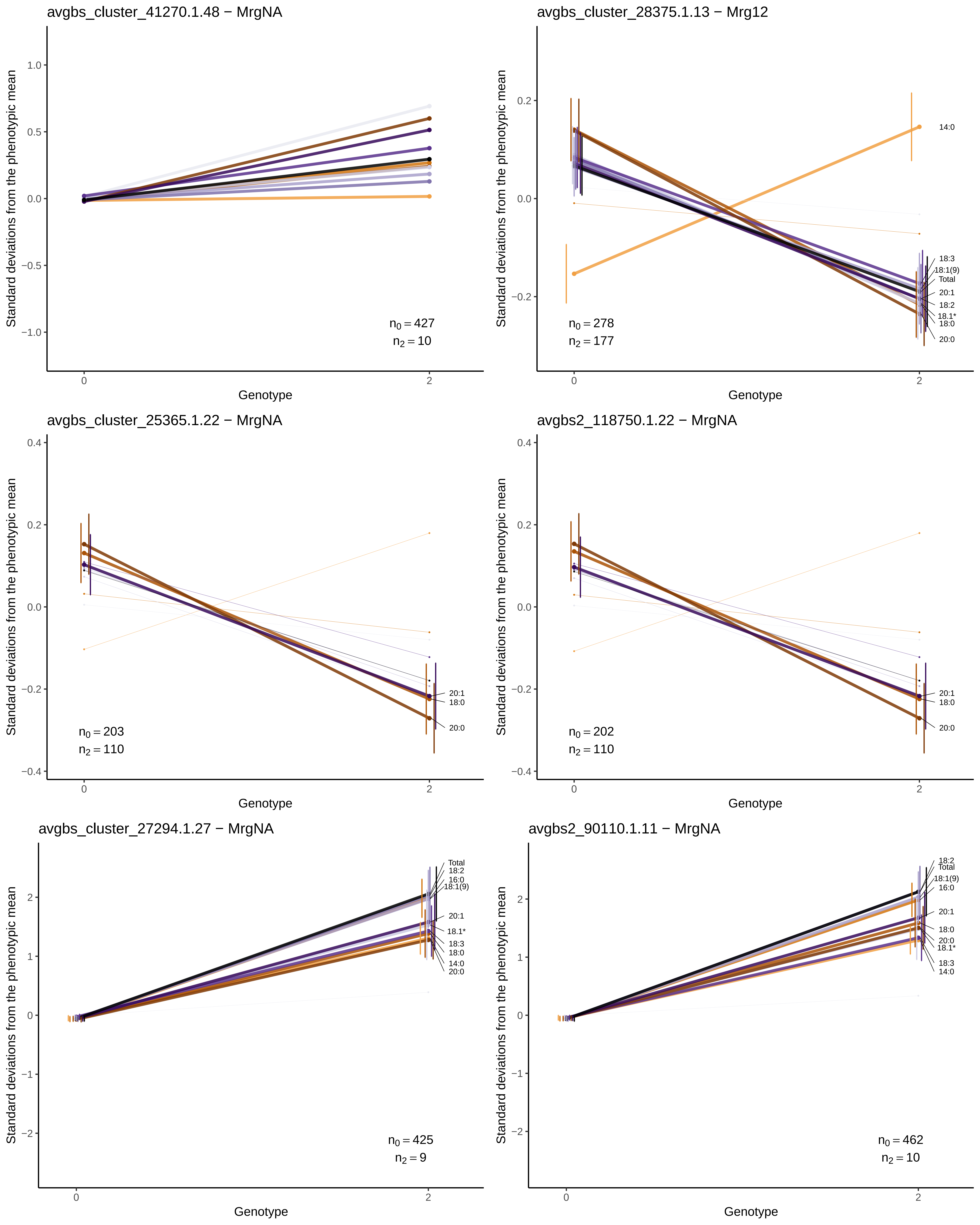
avgbs_cluster_41270.1.48 − MrgNA
$n_0 = 427$
$n_2 = 10$
avgbs_cluster_28375.1.13 − Mrg12
14:0
18:3
18:1(9)
Total
20:1
18:2
18.1*
18:0
20:0
$n_0 = 278$
$n_2 = 177$
avgbs_cluster_25365.1.22 − MrgNA
20:1
18:0
20:0
$n_0 = 203$
$n_2 = 110$
avgbs2_118750.1.22 − MrgNA
20:1
18:0
20:0
$n_0 = 202$
$n_2 = 110$
avgbs_cluster_27294.1.27 − MrgNA
Total
18:2
16:0
18:1(9)
20:1
18.1*
18:3
18:0
14:0
20:0
$n_0 = 425$
$n_2 = 9$
avgbs2_90110.1.11 − MrgNA
18:2
Total
18:1(9)
16:0
20:1
18:0
20:0
18.1*
18:3
14:0
$n_0 = 462$
$n_2 = 10$
Standard deviations from the phenotypic mean
Genotype

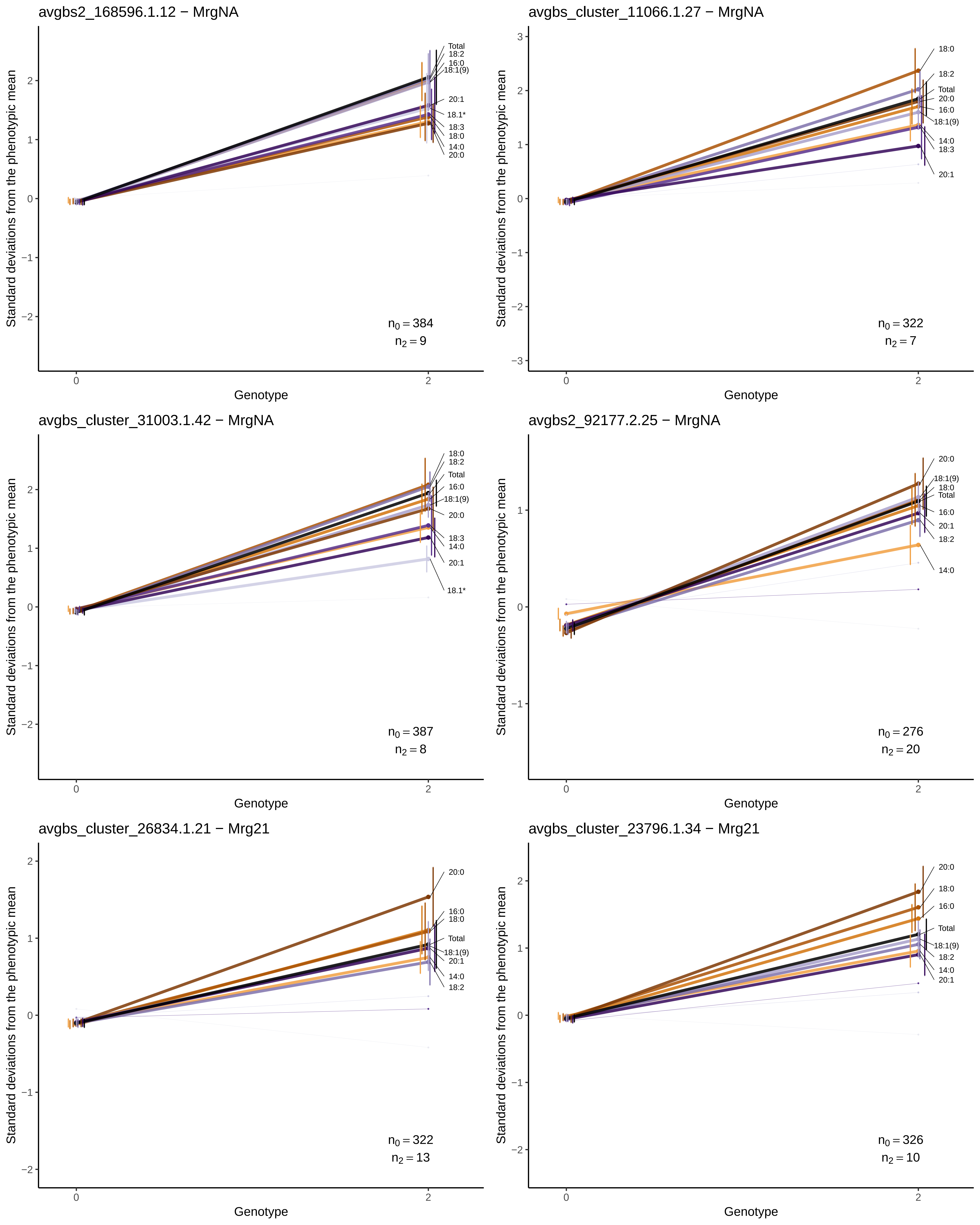
avgbs2_168596.1.12 − MrgNA
Standard deviations from the phenotypic mean
Genotype
Total
18:2
16:0
18:1(9)
20:1
18.1*
18:3
18:0
14:0
20:0
$n_0 = 384$
$n_2 = 9$
avgbs_cluster_11066.1.27 − MrgNA
Standard deviations from the phenotypic mean
Genotype
18:0
18:2
Total
20:0
16:0
18:1(9)
14:0
18:3
20:1
$n_0 = 322$
$n_2 = 7$
avgbs_cluster_31003.1.42 − MrgNA
Standard deviations from the phenotypic mean
Genotype
18:0
18:2
Total
16:0
18:1(9)
20:0
18:3
14:0
20:1
18.1*
$n_0 = 387$
$n_2 = 8$
avgbs2_92177.2.25 − MrgNA
Standard deviations from the phenotypic mean
Genotype
20:0
18:1(9)
18:0
Total
16:0
20:1
18:2
14:0
$n_0 = 276$
$n_2 = 20$
avgbs_cluster_26834.1.21 − Mrg21
Standard deviations from the phenotypic mean
Genotype
20:0
16:0
18:0
Total
18:1(9)
20:1
14:0
18:2
$n_0 = 322$
$n_2 = 13$
avgbs_cluster_23796.1.34 − Mrg21
Standard deviations from the phenotypic mean
Genotype
20:0
18:0
16:0
Total
18:1(9)
18:2
14:0
20:1
$n_0 = 326$
$n_2 = 10$

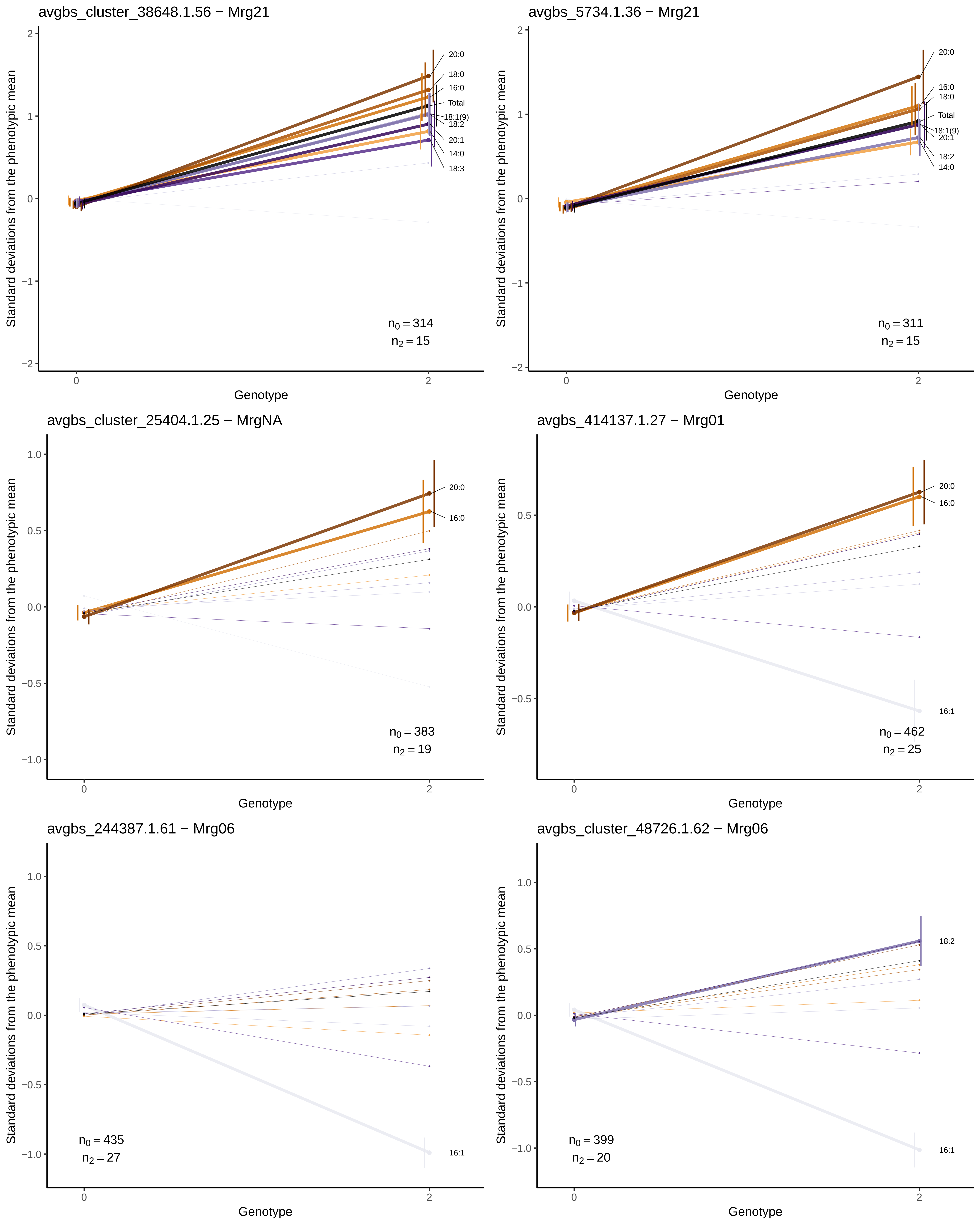

avgbs_cluster_38648.1.56 − Mrg21
avgbs_5734.1.36 − Mrg21
avgbs_cluster_25404.1.25 − MrgNA
avgbs_414137.1.27 − Mrg01
avgbs_244387.1.61 − Mrg06
avgbs_cluster_48726.1.62 − Mrg06
Standard deviations from the phenotypic mean
Genotype
20:0
18:0
16:0
Total
18:1(9)
18:2
20:1
14:0
18:3
16:1
$n_0 = 314$
$n_2 = 15$
$n_0 = 311$
$n_2 = 15$
$n_0 = 383$
$n_2 = 19$
$n_0 = 462$
$n_2 = 25$
$n_0 = 435$
$n_2 = 27$
$n_0 = 399$
$n_2 = 20$

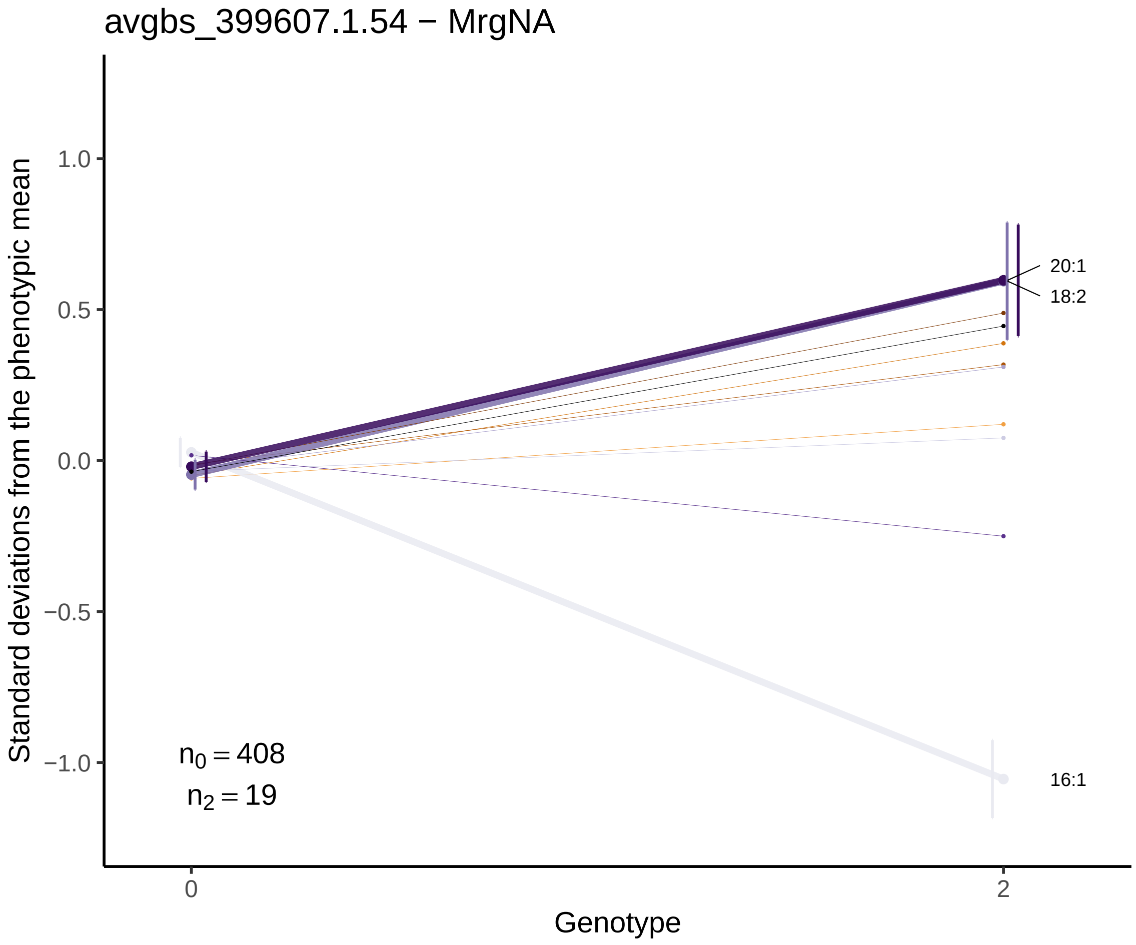
avgbs_399607.1.54 − MrgNA
Standard deviations from the phenotypic mean
1.0
0.5
0.0
−0.5
−1.0
20:1
18:2
16:1
$n_0 = 408$
$n_2 = 19$
0
2
Genotype

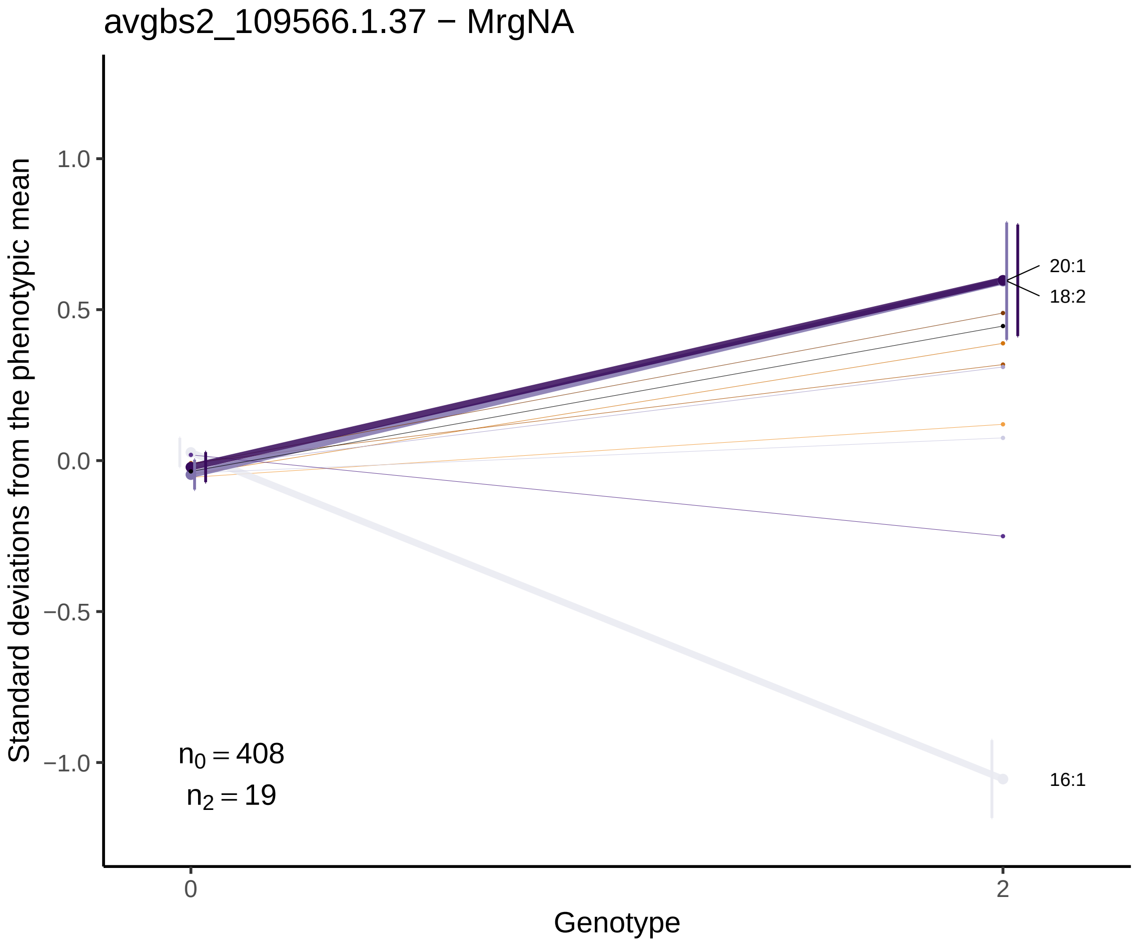
avgbs2_109566.1.37 − MrgNA
Standard deviations from the phenotypic mean
1.0
0.5
0.0
−0.5
−1.0
20:1
18:2
16:1
$n_0 = 408$
$n_2 = 19$
0
2
Genotype
